# Application of a French cattle pangenome, from structural variant discovery to association studies on key phenotypes

**DOI:** 10.1101/2025.04.15.648672

**Authors:** Valentin Sorin, Maulana Mughitz Naji, Clément Birbes, Cécile Grohs, Clémentine Escouflaire, Sébastien Fritz, Camille Eché, Camille Marcuzzo, Amandine Suin, Cécile Donnadieu, Christine Gaspin, Carole Iampietro, Denis Milan, Laurence Drouilhet, Gwenola Tosser-Klopp, Didier Boichard, Christophe Klopp, Marie-Pierre Sanchez, Mekki Boussaha

**Affiliations:** Université Paris Saclay, INRAE, AgroParisTech, GABI, 78350 Jouy en Josas, France; Genotoul Bioinfo, BioInfoMics, MIAT UR875, Sigenae, INRAE, Castanet-Tolosan, France; Eliance, 149 Rue de Bercy, 75012 Paris, France; INRAE, US 1426, GeT-PlaGe, Genotoul, France Génomique, Université de Toulouse, Castanet-Tolosan, France; GenPhySE, Université de Toulouse, INRAE, ENVT, 31326 Castanet-Tolosan, France

## Abstract

**Background:** The current cattle reference genome assembly, a pseudo-linear sequence produced using sequences from a single Hereford cow, represent a limit when performing genetic studies, especially when investigating the whole spectrum of genetic variations within the species. Detecting structural variations (SVs) poses significant challenges when relying solely on conventional methods of short or long-read sequence mapping to the current bovine genome assembly.

**Results:** In this study, we used long-reads (LR) and bioinformatic tools to construct a comprehensive bovine pangenome incorporating genetic diversity of 64 good quality *de novo* genome assemblies representing 14 French dairy and beef cattle breeds. Using a combination of complementary approaches, we explored the pangenome graph and identified 2.563 Gb of sequences common to all samples, and cumulated 0.295 Gb of variable sequences. Notably, we discovered 0.159 Gb of novel sequences not present in the current Hereford reference genome assembly. Our analysis also revealed 109,275 SVs, of which 84,612 were bi-allelic, including 21,840 insertions and 21,340 deletions. Genome-wide association studies using SNPs and a panel of 221 SVs, shared between the pangenome and the EuroGMD chip, revealed several well-known QTLs across the genome for the Holstein, Montbéliarde and Normande breeds. Among those, a QTL on chromosome 11 presents an SV with a highly significant effect on stature in the Holstein breed. This SV is a 6.2 kb deletion affecting the 5’UTR, first exon and part of first intron of *MATN3* gene, suggesting a potential regulatory and coding effect.

**Conclusions:** Our study provides new insights into the genetic diversity of 14 French dairy and beef breeds and highlights the utility of pangenome graphs in capturing structural variation. The identified SV associated with stature highlights the importance of integrating SVs into GWAS for a more comprehensive understanding of complex traits.

## Background

In many species, genetic analyses have traditionally relied on a single linear reference assembly, often used for sequence alignments, identification of genomic variations, and genome annotation. However, these reference assemblies are far from perfect; they often lack millions of base pairs and fail to accurately reflect the genetic diversity that can be captured within the species, potentially introducing biases into the genomic studies. The current bovine reference genome assembly (*Bos taurus taurus*) was initially built using whole-genome sequencing of a single inbred Hereford cow, «L1 Dominette 01449» [1]. Over the years, improvements of sequencing technologies, combined with the development of bioinformatic methods for genome assembly, have greatly improved genome contiguity and accuracy, thereby facilitating the characterization of a wide range of genomic variations. However, the current linear reference genome assembly still contains hundreds of gaps [2], including stretches of missing sequence spanning several megabases (Mb) (∼4.6 Mb of N-stretch), and only represents one haplotypic reference of the species [3,4].

Recent advances in long-read (LR) sequencing technologies now enable the production of more continuous genome sequences, with accuracy comparable to those obtained with short-read (SR) sequencing [5]. SR sequencing is especially valuable for the identification and characterization of single nucleotide polymorphisms (SNPs) as well as small insertions and deletions (InDels) in non-repeated part of the assemblies. In contrast, LR sequencing is more effective in detecting complex structural variations (SVs), which includes large deletions, insertions, inversions, duplications, and translocations. Moreover, high-quality LR sequencing also enable the detection of small variant in repeated part of the assemblies. Additionally, LR sequences facilitates the creation of high-quality *de novo* genome assemblies, which serve as the foundation for multi-assembly graphs, commonly referred to as pangenomes [6–8].

Pangenome are graphs build from a combination of assemblies from multiple individuals [9], encompassing both the core genome, which consists of nodes, that represent sequences, common to all individuals, and the flexible genome, which represents genomic sequences found only in a subset of individuals. These pangenome graphs are of particular interest as they provide a more comprehensive representation of the genetic diversity within a species, hence facilitating a more detailed identification of mutations that contribute to phenotypic variations.

In recent years, several computational tools have been developed for the construction of pangenome graphs, each with different specificities. The **Minigraph** software [10] efficiently detects large structural variants (≥ 50 bp) using a reference genome as a backbone but it is not well-suited for studying small genomic variations. The **Minigraph-Cactus** tool [11] extends the previous approach by incorporating smaller DNA variations, such as SNPs and InDels. Both Minigraph and Minigraph-Cactus allow the iterative addition of assemblies based on their phylogenetic proximity to the reference genome. Alternatively, the pangenome graph builder (**pggb**) uses a reference-free approach, thus reducing bias in graph construction [12]. Although computationally demanding, pggb enables fine-scale genetic variant detection across entire chromosomes, similar to Minigraph-Cactus.

In the present study, we used Minigraph to construct a pangenome graph using 64 *de novo* genome assemblies representing 14 French cattle breeds. Using this graph, we carried out a comprehensive analysis of SVs, assessing their contribution to population structure based on genotypes, and investigating their associations with key phenotypes in the three main French dairy cattle breeds.

## Methods

### *De novo* genomes assemblies processing

In this study we used 64 publicly available *de novo* genome assemblies produced by assembling PacBio Continuous Long Reads (CLR) long-read sequences from animals corresponding to 14 dairy and beef breeds. The number of individuals per breed ranged from two (Rouge Flamande) to eight (Holstein) (Table 1 & Additional file 1: Table S1). Our panel of *de novo* genome assemblies contains both widely used breeds (*e.g.* Holstein, Normande and Montbéliarde) and more rustic and local French breeds (*e.g.* Vosgienne, Tarentaise and Abondance).

**Table 1.** Distribution of assemblies per breed.

| **Breed** | **Breed abbreviation** | **Number of assemblies** |
| --- | --- | --- |
| Abondance | ABO | 5 |
| Aubrac | AUB | 7 |
| Blonde d'Aquitaine | BAQ | 4 |
| Brown Swiss | BSW | 5 |
| Charolaise | CHA | 4 |
| Holstein | HOL | 8 |
| Limousine | LIM | 2 |
| Montbéliarde | MON | 5 |
| Normande | NMD | 7 |
| Parthenaise | PAR | 3 |
| Rouge Flamande | RDC | 2 |
| Simmental | SIM | 3 |
| Tarentaise | TAR | 5 |
| Vosgienne | VOS | 4 |
| **Total** |  | **64** |

As PacBio CLR LRs are prone to high-level sequence errors, a multi-step polishing process was applied using both the CLR and Illumina SR raw sequence data in order to improve the quality of the genome assembly sequences. Firstly, raw CLR data were aligned to the primary assembly contigs using **pbmm2**, a tool from the SMRTLink v12.0 workflow manager [13], and **GCpp**, a tool to compute a genomic consensus from these alignments. Secondly, two additional rounds of polishing were applied using high-quality Illumina SR data with the **Pilon** software (v1.24) [14], enabling the correction of small-scale sequence errors across the assemblies. Finally, contigs were scaffolded to chromosome-level assemblies using the **RagTag** software (v2.1.0) [15], with the ARS-UCD1.2 bovine reference genome as the backbone [2].

We assessed the quality of the genome assemblies using three main metrics: total assembly length, N50 score, and completeness assessment. Completeness of the assemblies was assessed using the **BUSCO** score (v5.4.7) [16] with the mammalian single-copy orthologous gene dataset (mammalia_odb10. 2024-01-08) as a reference.

### Construction of a pangenome graph

The pangenome graph was constructed using the latest ARS-UCD1.2 bovine reference assembly as backbone, with the 64 *de novo* genome assemblies being aligned iteratively based on their phylogenetic relationships. Firstly, we assessed the phylogenetic distance between samples using the **Mash** software (v2.3) [17]. Given the large size of cattle genomes, we established two key parameters within the Mash sketch function: a k-mer size of 32 using the “-k” option and a sketch size of 100,000,000 using the “-s” option. The resulting distance matrix was used to build a phylogenetic tree with the **ape** library [18] in **R** (4.3.1), and the tree was visualized with the **plot.phylo** function [19].

Subsequently, we constructed the pangenome graph using **Minigraph** (v0.21), with the “-cxggs” parameter to perform base-alignment. The constructed pangenome graph consisted of a series of bubbles, with the reference genome serving as the primary path. Finally, we visualized the pangenome graph through **BandageNG** (v2022.09) [20].

### Structural variations (SVs) calling and non-reference unique insertion sequences (NRUIs) extraction

Pangenome graph SVs are depicted as bubbles, where each bubble represents sequence variations across genome assemblies. We identified the allele present in each bubble for all samples by individually realigning the 64 assemblies to the pangenome graph using the “-cxasm --call” option of Minigraph. Node labelling was then generated along each genome’s alignment path, and the allele information for each sample was stored in a corresponding bed file. The 64 individual bed files were subsequently combined into a single VCF file following the **Minigraph-cookbook-v1** guidelines [21]. Briefly, we first used the **k8** tool alongside with the **mgutils.js** script to create a detailed bed file that encompassed all identified alleles across individuals. Subsequently, the bed file was converted into VCF format using the **mgutils-es6.js** script with the **merge2vcf -r0** option. Variants for which all individuals presented an unknown genotype (*e.g.*, “*./.*”) were filtered out from the final panel.

Each SV was subsequently classified based on two distinct criteria: number of alleles and SV type. Firstly, SVs were considered as biallelic if the bubble contained exactly two paths (reference and alternative), and as multi-allelic if more than two paths were present. Secondly, SVs were classified as insertions when the reference path contained no sequence (length = 0) and the alternative path contained an insertion sequence of at least 50 nucleotides, and as deletions when the alternative path contained no sequence while the reference path retained a sequence of at least 50 nucleotides. All remaining SVs, where both the reference and the alternative (non-reference) paths contained genomic sequences, have been considered as substitutions.

Finally, we extracted non-reference sequences (NRSs) by applying the following three filters : (1) all nodes that did not successfully occur in any of the individual paths were considered as nested nodes and where excluded from further analysis; (2) all nodes that occurred in the reference path were also excluded; and (3) all nodes that passed the first 2 filters and have a sequence length higher than 50 bases where selected and where considered as non-reference sequences. We also extracted non-reference unique insertion sequences (NRUIs) by identifying only true insertion-type SVs (representing sequences absent from the ARS-UCD1.2 reference genome). Nodes corresponding to these insertions were selected, and sequences exceeding 50 bp in length were retained and classified as NRUI sequences.

### SVs validation of SVs by high-throughput genotyping

To evaluate the efficiency and population-level relevance of our SV detection approach, we applied a previously developed high-throughput genotyping strategy based on the bovine Illumina EuroGMD SNP array [22]. In our study, we focused only on deletions and we applied our previously reported method to convert deletions into virtual SNPs and add these to the SNP chip [23]. Briefly, the predicted deletions were converted into “virtual SNPs” by testing the base change at the SV breakpoints as follows: if the first nucleotide of the deleted region was different from the first nucleotide which was located immediately after the SV 3’ breakpoint, then the reference allele of the “virtual SNP” corresponds to the first nucleotide of the deleted region and the alternative allele corresponded to the first nucleotide immediately after the SV 3’ breakpoint. This genotyping can be confirmed by performing a complementary test on the opposite strand of the DNA. This cost-effective method enables the genotyping of multiple SVs across large populations and, using this approach, we compiled the genotyping database used for genomic selection for 230 SVs across 2,838,235 animals from 21 French dairy and beef cattle breeds (Table 2).

**Table 2.** Number of animals genotyped for the 1900 structural variants.

| **Breed** | **Number of genotyped animals** |
| --- | --- |
| Abondance | 26,970 |
| Aubrac | 11,580 |
| Bazadaise | 133 |
| Blonde d'Aquitaine | 33,658 |
| Bretonne Pie Noire | 57 |
| Brown Swiss | 78,632 |
| Charolaise | 80,880 |
| Holstein | 1,593,213 |
| Inra 95 | 549 |
| Jersey | 12,179 |
| Limousine | 18,742 |
| Montbéliarde | 715,854 |
| Normande | 218,455 |
| Parthenaise | 5630 |
| Rouge des Prés | 560 |
| Rouge Flamande | 345 |
| Salers | 5472 |
| Simmental | 15,653 |
| Tarentaise | 13,891 |
| Vosgienne | 5747 |
| Other cattle breeds | 35 |
| **Total** | **2,838,235** |

Among the SVs identified in our pangenome-based approach, 230 deletions were already present in this existing database with available genotypes. We extracted the genotypes for these 230 SVs and estimated their allele frequencies both at the global population level and within each breed.

In addition, we evaluated two standard population genetic metrics to assess the informativeness of these SVs. Firstly, we calculated the heterozygosity ratio (He), defined as He=2pq, where *p* and *q* are the frequencies of the two alleles. Secondly, we computed the Polymorphic Information Content (PIC) score, which is given by the formula PIC=He-2p^2^q^2^ [24]. The PIC score measures the ability of a marker in detecting genetic polymorphisms of interest in linkage analysis.

### Analysis of population structure

To characterize the distribution of SVs and NRUIs across cattle breeds, we firstly conducted a population structure analysis based on the presence/absence variation (PAV) of SVs and NRUIs. For this purpose, we generated two binary PAV-matrices, each recording the presence or absence of either SV or NRUI for all individuals. Hierarchical clustering was then performed on the SV and NRUI PAV-matrices using the **HCLUST** function in **R** [25], enabling the visualization of breed clustering patterns based on SVs and NRUIs.

To further evaluate the reliability of population structure inferred from SVs, we used the genotyping data of the 230 SVs to carry out a Principal Components Analysis (PCA) using the “dudi.pca” function implemented in the R package **ade4** [26]. This PCA was based on validated SVs genotypes from a panel of 1500 bulls from the three main dairy breeds (Holstein (HOL), Montbéliarde (MON) and Normande (NMD)) used in our GWAS studies. These validated SVs correspond to the SVs shared between the pangenome and EuroGMD genotyping array. The population structure inferred from SVs was then compared to previously reported results based on 50k SNP data [27], allowing us to assess the consistency of SV-based clustering.

### Genome-wide association analyses

To assess the potential effects of the identified SVs on dairy cow performances, we conducted genome-wide association studies (GWAS) using the EuroGMD genotyping chip, which included the panel of 230 SVs detected through our pangenome-based approach.

The phenotypes used in GWAS were daughter yield deviations (DYD), defined as the average value of daughters’ performances, adjusted for systematic environmental effects and for the breeding value of their mates [28]. Bulls with records from at least 20 daughters with available phenotypes were included in the analysis.

#### Imputation of structural variants genotypes

SVs have been included since version 2 of the EuroGMD array. Specifically, 182 SVs are present in both v2 and v3, with an additional 48 included in v4, bringing the total to 230 SVs. These SVs are distributed across the entire genome but in a non-uniform manner, with only 2 SVs on BTA12 (*Bos taurus* autosome) and 28 SVs on BTA1 (Fig. 1). As the bulls had been genotyped using different versions of the EuroGMD chip, some of which did not initially contain the 230 SVs, we imputed the missing SV genotypes across the study population before conducting the association analyses. Following imputation, GWAS analyses were performed to investigate the associations between SV genotypes and key production and functional traits in dairy cattle.

**Figure 1.**
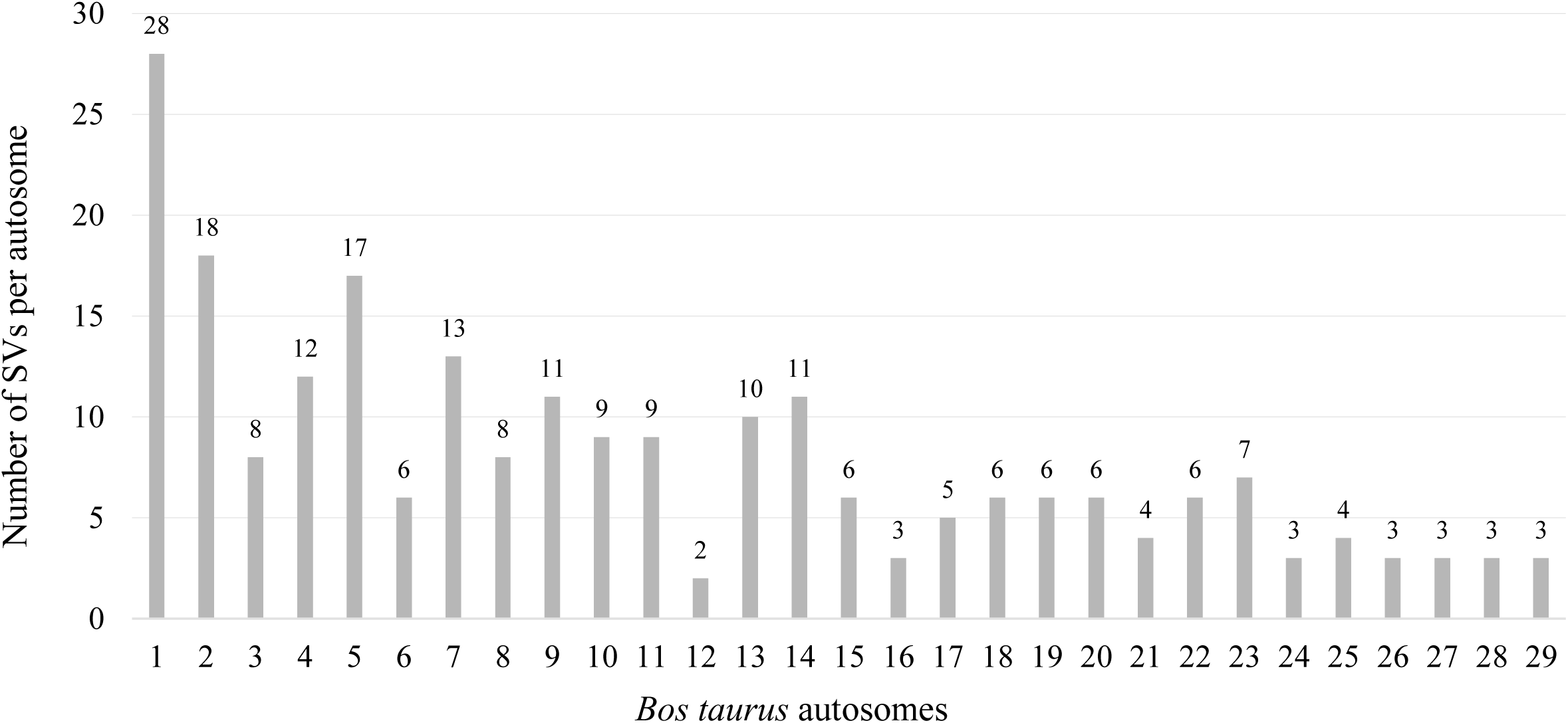
Distribution of the 230 structural variants present on the EuroGMD chip across all bovine autosomes *SVs* Structural variants

Genotypes for the 230 SVs were available for a subset of animals genotyped using different versions of the EuroGMD that included these SVs. These data were used to establish breed-specific reference populations for the three main French breeds under study (HOL, MON and NMD). To maximise the accuracy of the imputed genotypes, the selection of animals for the reference populations was guided by two main criteria: (i) the inclusion of widely used artificial insemination (AI) bulls in France, and (ii) the selection of animals closely related to the bulls targeted for imputation based on pedigree data for the HOL, MON and NMD breeds.

Imputation was performed using the **FImpute** software [29]. Each reference population was designed to include approximately 20,000 animals. To refine the selection, filters were applied based on the number of SVs genotyped per animal. Depending on breed-specific availability, a threshold was set at 180 out of 182 SVs and at 228 out of 230 SVs for EuroGMD v2, v3, and EuroGMD v4, respectively, in MON and HOL. For the NMD breed, the thresholds were set at 170 out of 182 SVs and 220 out of 230 SVs for EuroGMD v2, v3, and EuroGMD v4, respectively. After applying these filters, sires and dams were selected for all chip versions. Where possible, one descendant per bull from the GWAS population was progressively added for each chip version.

#### Genome wide association study

We evaluated the effect of the 230 SVs panel in the three main dairy breeds (HOL, MON and NMD) for the following 13 traits:

- Five fertility traits: heifer conception rate (HCR), cow conception rate (CCR), calving -first artificial insemination interval (ICAI1), heifer non-return rate (HNRR) and cow non-return rate (CNRR);
- Five milk production traits: milk yield (MY), fat content (FC), protein content (PC), fat yield (FY) and protein yield (PY);
- Two udder health traits: somatic cell score (SCS) and clinical mastitis (CM);
- One morphology trait: height at sacrum (HS)

Depending on the trait, between 3096 and 3672 bulls, 2417 and 2986 bulls, and 9822 and 11,428 bulls with DYDs were used for the association analyses (Table 3) for the MON, NMD, and HOL breeds, respectively.

**Table 3.** Number of MON, NMD, and HOL bulls with daughters’ performance for each trait.

| Type of trait |  | MON | NMD | HOL |
| --- | --- | --- | --- | --- |
| **Fertility** | Heifer conception rate (HCR) | 3394 | 2764 | 10,950 |
|  | Cow conception rate (CCR) | 3445 | 2882 | 10,978 |
|  | Calving-First insemination Interval (ICAI1) | 3508 | 2889 | 11,075 |
|  | Heifer non-return rate (HNRR) | 3424 | 2771 | 10,924 |
|  | Cow non-return rate (CNRR) | 3489 | 2892 | 10,953 |
| **Milk production** | Milk yield (kg) (MY) | 3672 | 2979 | 11,423 |
|  | Fat content (%) (FC) | 3672 | 2979 | 11,422 |
|  | Protein content (%) (PC) | 3672 | 2979 | 11,422 |
|  | Fat yield (kg) (FY) | 3672 | 2979 | 11,420 |
|  | Protein yield (kg) (PY) | 3672 | 2979 | 11,420 |
| **Udder health** | Somatic cell score (SCS) | 3620 | 2986 | 11,448 |
|  | Clinical mastitis (CM) | 3096 | 2417 | 9822 |
| **Morphology** | Height at sacrum (HS) | 3599 | 2720 | 10,609 |

*MON* Montbéliarde; *NMD* Normande; *HOL* Holstein

We performed GWAS with 58,191 genomic variants, including 230 SVs, for each breed separately, analysing one trait at a time. We used the *mlma* (Mixed Linear Model Analysis) approach implemented in the GCTA software. This method applies a mixed linear model including the variant to be tested:

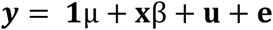

Where **y** represents the vector of DYD; μ is the overall mean; β is the fixed additive effect of the variant being tested for association; **x** is the vector of imputed genotypes, coded as the number of copies of the tested allele 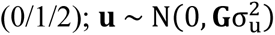 is the vector of random polygenic effects, where **G** is the genomic relationship matrix (GRM) derived from 50k SNP genotypes, and 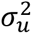 is the polygenic variance, estimated from the null model (**y** = **1**μ + **u** + **e**), and then held fixed while testing the association between each variant and the trait of interest; finally, 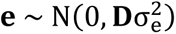 is the vector of random residual effects, with 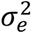 the residual variance. **D** was a diagonal matrix with inverse weights for DYD to account for heterogeneous accuracy.

A genome-wide significance threshold was applied using the Bonferroni correction, calculated as P-value = 0.05 / 58,191, where 0.05 is the nominal significance level and 58,191 is the number of variants analysed. This resulted in a significance threshold of P-value = 8.59 × 10^−7^, corresponding to -log_10_(P-value) = 6.07, rounded to 6.1.

#### QTL identification

To determine the number of QTL regions associated with each phenotype, we applied an iterative procedure as described in Sanchez *et al.* [30], for each breed separately. Briefly, this procedure aims at (i) defining confidence intervals (CIs) for QTL based on LD between SNPs and (ii) identifying putative multiple QTLs within a given region. The procedure was applied using the following parameters: a window size of 10 Mb and an LD threshold of 0.7.

In the identified QTL regions, most significant SVs were selected to precisely determine their genomic localization. Therefore, breakpoint coordinates of each SV were extracted and mapped its position onto the reference genome. To functionally annotate these SVs, whether they directly affected a gene or were located near gene regulatory regions was checked, using the **Ensembl** [31] and **UCSC** [32] databases. This approach follows the methodology described by Boussaha *et al.* [23] and Letaief *et al.* [33].

### Pangenome validation of the SV within *MATN3* gene

In QTL regions where an SV showed a significant effect on the phenotype, we validated the functional impact of the SV by constructing a local pangenome over a 2 Mb region surrounding the locus. To achieve this, the sequences of each animal from the panel of 64 CLR assemblies corresponding to the ARS-UCD1.2 reference genome coordinates were extracted following three main steps: (i) aligning genomes against the reference using **minimap2** (v2-2.28) [34], (ii) extracting the region of interest using **IMplicite Pangenome Graph (IMPG)** (v0.2.1) [35], and (iii) obtaining the corresponding FASTA sequences using **samtools faidx** (v1.20) [36]. Finally, a pangenome was build using Minigraph as described previously and visualized the region on the graph using **BandageNG** [20].

### Validation by PCR and Sanger sequencing of the deletion within *MATN3* gene

Targeted SV breakpoints were validated by PCR amplification and Sanger sequencing. Six animals were used in the validation step: two heterozygotes, two homozygotes for the deletion, and two non-carrier animals. Primers were produced by Eurofins Genomics (Ebersberg, Germany). We designed these primers to amplify two distinct amplicons (Table 4), allowing the detection of presence or absence of the deletion. The first amplicon, which was 422 bp in size, was amplified using the A1-A3 primer pair and identified homozygous individuals for the reference allele (*i.e.*, non-carriers of the deletion). The second amplicon, obtained with the A1-A2 primers flanking the deletion breakpoints, measured 580 bp and identified individuals carrying the deletion, corresponding to the alternative allele (Fig. 2). The size difference between these two fragments allowed the distinction of heterozygous individuals, which displayed both amplicons at the same time.

**Figure 2.**
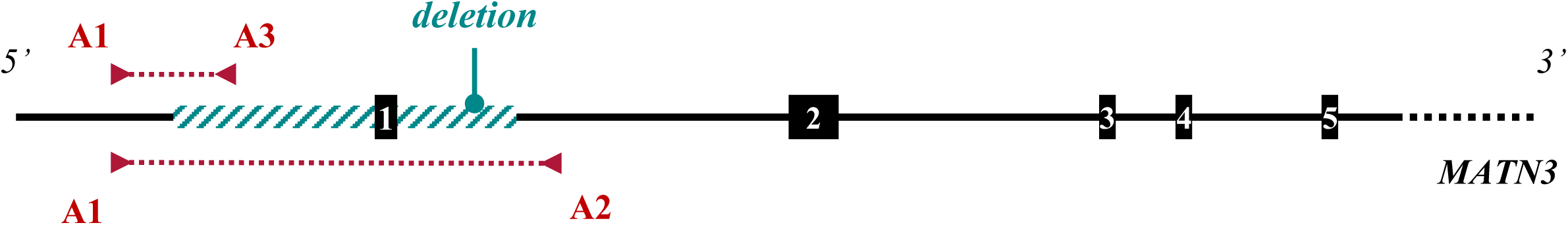
PCR-based validation of a deletion Structure of the *MATN3* gene. Black boxes and lines represent first 5 exons and introns, respectively. Primers pairs A1-A2, A1-A3 are indicated in red. The blue hatched box represents the deletion.

**Table 4.**
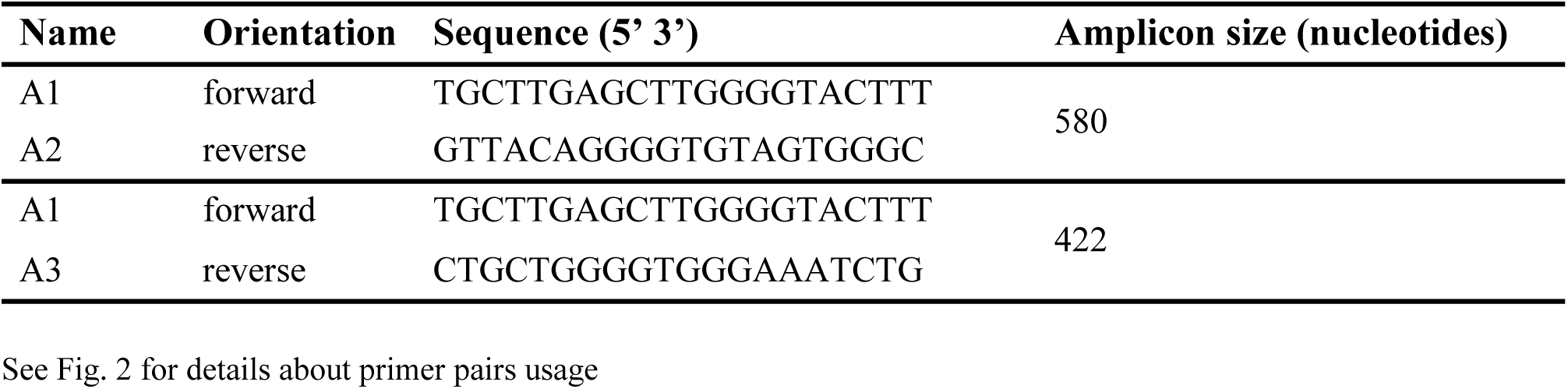
Features of PCR primers.

PCR was performed using 100 ng of DNA in a 35 µL reaction mixture consisting of 1X GoTaq Flexi Buffer [37], 2.5 mM MgCl_2_, 800 µM dNTPs, 0.875 UI GoTaq DNA polymerase [37], and 0.5 µM of each primer (A1, A2, and A3). The PCR program consisted of an initial denaturation step at 94°C for 3 min, followed by 35 three-step cycles: (i) denaturation at 94°C for 30 s, (ii) annealing at 67°C for 30 s, and (iii) extension at 72°C for 1 min, and a final extension step at 72°C for 5 min before cooling to 15°C. PCR products were visualized by gel electrophoresis on a 2% agarose gel. Additionally, PCR products were sequenced by Eurofins Genomics (Ebersberg, Germany) to determine the precise breakpoint sequences of the studied SVs.

## Results

### Quality of the 64 *de novo* genome assemblies

The current ARS-UCD1.2 reference assembly metrics are: 2,759,153,975 bases total size including unmapped contigs, 103.3 Mb N50 and 95.8% BUSCO scores. To assess the quality of our final panel of 64 polished *de novo* assemblies, we computed the genome length, N50 metrics, and BUSCO scores (Additional file 1: Table S1). The 64 *de novo* genome assemblies presented an average genome length of 2,636,580,836 bases, ranging from 2,612,227,180 to 2,688,017,017 bases. The average N50 scaffold length was approximately 100 Mb, with values ranging from 88 to 106.5 Mb and an average L50 scaffold count of around 12. These high N50 and low L50 scores suggested that the genome assemblies were contiguous and exhibit minimal fragmentation. The average BUSCO score was 95.1%, with individual scores ranging from 93.9% to 95.7%. These high BUSCO values reflected high genome completeness. Furthermore, alignment of these *de novo* assemblies with D-Genies [38] (Additional file 2: Figures S1-S14) showed a high degree of similarity and concordance with the chromosomes of the ARS-UCD1.2 reference genome assembly.

Collectively, these quality metrics confirm that all 64 genome assemblies were of sufficient quality and suitable for constructing a cattle pangenome graph.

### Characterization of the cattle pangenome graph

We constructed a cattle pangenome graph using the 64 *de novo* assemblies, with the ARS-UCD1.2 reference genome sequence used as backbone. Firstly, we estimated phylogenetic distances between assemblies using Mash and clustered samples into 14 groups according to their breed of origin (Fig. 3). These distances were subsequently used to construct a phylogenetic tree. The resulting pangenome graph consisted of 521,756 nodes connected by 735,034 edges, representing a total sequence length of 2,933,608,906 bases. Notably, 5.95% of the graph (174,454,931 bases) corresponded to sequences missing from the ARS-UCD1.2 bovine reference genome assembly. To ensure the accuracy of node labelling, we realigned each individual genome assembly to the graph and traced the corresponding path for each sample. A total of 507,822 out of the 521,756 nodes were successfully linked into the paths, covering 2,858,048,002 bases. The remaining 13,935 nodes, representing 75,560,904 bases, did not occur in any genome. They were considered as nested nodes and were therefore excluded from further analysis.

**Figure 3.**
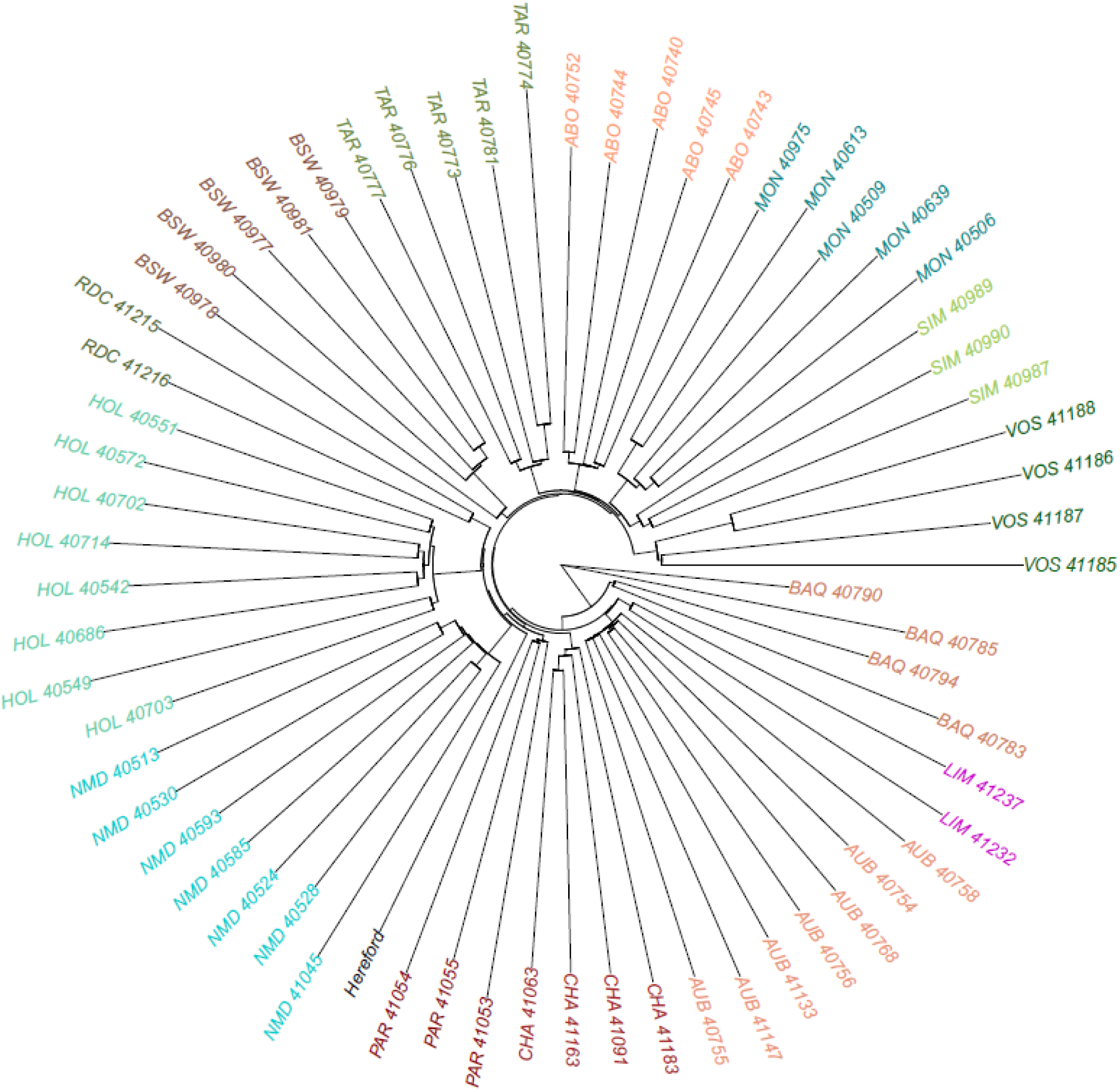
Phylogenetic distance between the 64 genome assemblies and ARS-UCD1.2 Phylogenetic tree derived from 64 bovine assemblies and the current Hereford reference genome assembly

The core pangenome contained 112,525 nodes, corresponding to 2,562,959,040 bases (90% of the graph). On the other hand, flexible pangenome regions contained 395,298 nodes, with a cumulative sequence length of 295,088,962 bases (10%). Within the flexible regions, 83,050 nodes containing 99,187,939 bases were identified as breed-specific (Table 5). Additionally, we identified 151,771 nodes with a cumulated sequence length of 158,701,735 bases that passed our filtering criteria and were therefore classified as non-reference nodes. We further analysed the 158.7 Mb of novel sequences (referred as NRSs) by focusing only on true insertions that originated from bubbles with no reference nodes. This analysis revealed 27,750 non-reference nodes, representing a total sequence length of 25,470,897 bases of true and good quality NRUIs. Notably, 9915 of these nodes, containing 13,344,651 bases, were identified as breed-specific, appearing exclusively in single breeds (Table 5).

**Table 5.** Distribution per breed of flexible sequences and non-reference unique insertions.

| **Breed** | **Number of assemblies** | **Number of flexible nodes** | **Total length of flexible sequences (nucleotides)** | **Number of breed-specific NRUIs nodes** | **Total length of breed-specific NRUIs (nucleotides)** |
| --- | --- | --- | --- | --- | --- |
| Hereford | 1 | 1719 | 8,747,369 | 0 | 0 |
| Abondance | 5 | 7388 | 8,930,960 | 717 | 790,073 |
| Aubrac | 7 | 7143 | 8,249,769 | 1064 | 1,559,693 |
| Blonde d'Aquitaine | 4 | 3847 | 4,228,801 | 528 | 866,336 |
| Brown Swiss | 5 | 6819 | 7,693,269 | 839 | 1,158,611 |
| Charolaise | 4 | 4349 | 4,947,248 | 640 | 948,814 |
| Holstein | 8 | 9137 | 10,685,671 | 1168 | 1,732,578 |
| Limousine | 2 | 2232 | 2,255,173 | 334 | 419,269 |
| Montbéliarde | 5 | 6353 | 7,332,347 | 739 | 921,838 |
| Normande | 7 | 8088 | 8,851,454 | 1060 | 1,240,542 |
| Parthenaise | 3 | 4638 | 4,765,273 | 557 | 648,038 |
| Rouge Flamande | 2 | 1838 | 1,780,683 | 4260 | 615,828 |
| Simmental | 3 | 5560 | 6,472,286 | 434 | 659,700 |
| Tarentaise | 5 | 5495 | 6,854,827 | 724 | 1,084,256 |
| Vosgienne | 4 | 8444 | 7,392,809 | 691 | 699,075 |
| **Total** | **65** | **83,050** | **99,187,939** | **9,915** | **13,344,651** |

Hierarchical clustering based on presence/absence variation (PAV) matrix of NRUIs (Fig. 4) successfully grouped samples according to their breed of origin, highlighting the strong association between these unique insertions and breed structure.

**Figure 4.**
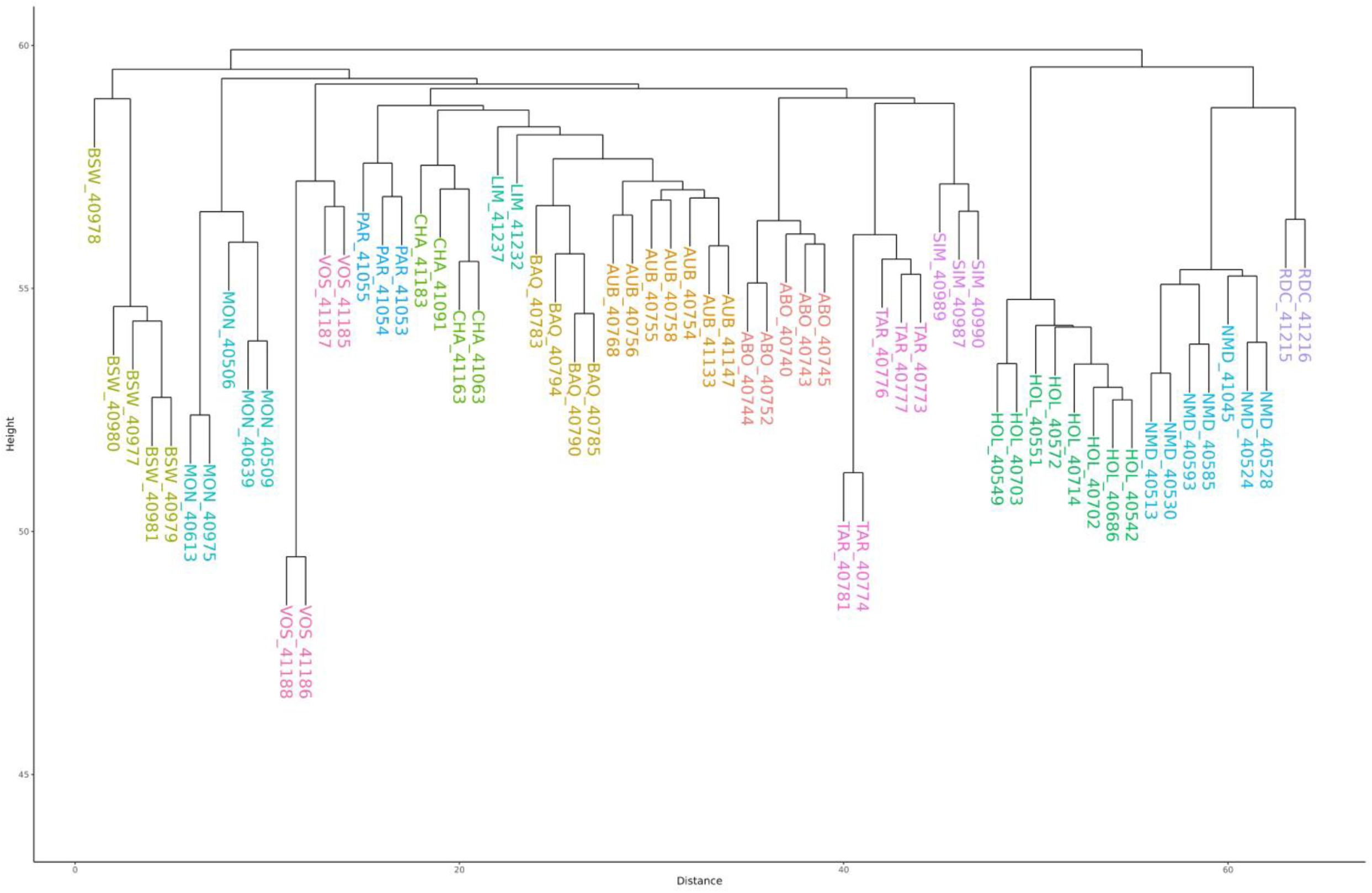
Hierarchical clustering of the 64 assemblies based on the NRUI PAV-matrix of *PAV* presence/absence variation; *NRUIs* non-reference unique insertions; Diagram showing the clustering of the 64 assemblies according to the 14 breeds

### Display SVs identified from the graph

In total, we identified 109,275 SVs (Table 6). Out of these, 77.4% (84,612 SVs) were classified as biallelic. Among the biallelic SVs, 55.4% were further categorized into 21,840 insertions (25.8%) and 21,340 deletions (25.2%). The remaining SVs corresponded to bubble that contained sequences in both the reference and non-reference paths, and were therefore classified as sequence substitutions. We analysed the length distribution of biallelic insertions and deletions (Additional file 3: Figures S15-S28) and observed a symmetric distribution between the sizes of SVs classified as insertions and those classified as deletions. Moreover, we identified several notable peaks at 150 bp, 250 bp, 5.5 kb and 8.6 kb, which likely correspond to structural variations associated with different families of transposable elements (SINE, LTR, LINE).

**Table 6.** Distribution of SVs by allele count and SV type.

| **Mutations** | **Bi-allelic count** | **Multi-allelic count** | **Total** |
| --- | --- | --- | --- |
| Insertions | 21,840 | 1997 | 23,837 |
| Deletions | 21,340 | 3437 | 24,777 |
| Sequence substitutions | 41,432 | 19,229 | 60,661 |
| **Total** | **84,612** | **24,663** | **109,275** |

Similar to results observed with NRUIs, hierarchical clustering based on PAV-matrix of SVs (Fig. 5) correctly assigned all samples to their corresponding breed of origin.

**Figure 5.**
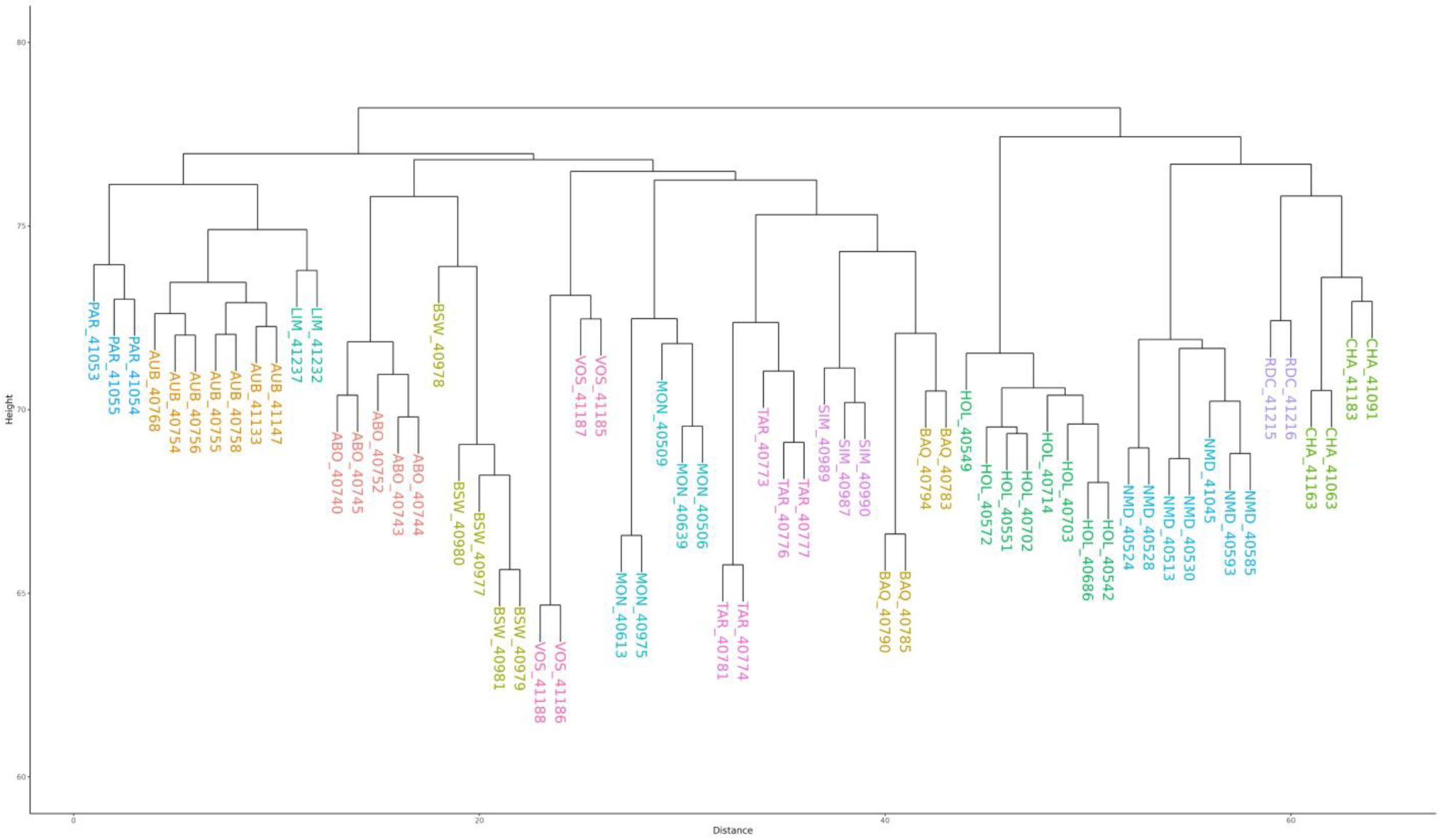
Hierarchical clustering of the 64 assemblies based on the SV PAV-matrix PAV presence/absence variation; SVs structural variants; Diagram showing the clustering of the 64 assemblies according to the 14 breeds

### Validation of SVs by SNP chip genotyping

Relevance of pangenome-derived SVs was assessed using the EuroGMD SNP chip genotyping data of 230 selected SVs obtained for a large population of animals representing 21 cattle breeds (Additional file 4: Table S2). All SVs were successfully genotyped, of which 221 were polymorphic. The 9 monomorphic SVs were likely to be either false positives or SVs specific to the Hereford reference genome assembly.

Mean observed minor allele frequency (MAF) across polymorphic SVs was 0.17 at the population level, ranging from 0.14 in Salers up to 0.21 in Bretonne Pie Noire (Additional file 4: Table S2). The mean observed heterozygosity across loci was 0.24, ranging from 0.20 in Salers to 0.29 in Bretonne Pie Noire. The mean PIC (Polymorphic Information Content) was 0.19 and varied from 0.17 in Salers to 0.23 in Bretonne Pie Noire (Additional file 4: Table S2). Heterozygosity and PIC are key parameters to assess the informativeness of genetic markers. Based on the values observed in our SV panel across the 21 breeds, these markers can be considered informative and are particularly suitable for population structure analysis and association studies.

To further assess the quality and informativeness of the validated SV panel, we investigated population structure using genotyping data from the three main French dairy breeds (MON, NMD and HOL). PCA accurately assigned all individuals to their breeds of origin (Fig. 6). These results provide additional statistical validation, complementing the validation of the SV panel.

**Figure 6.**
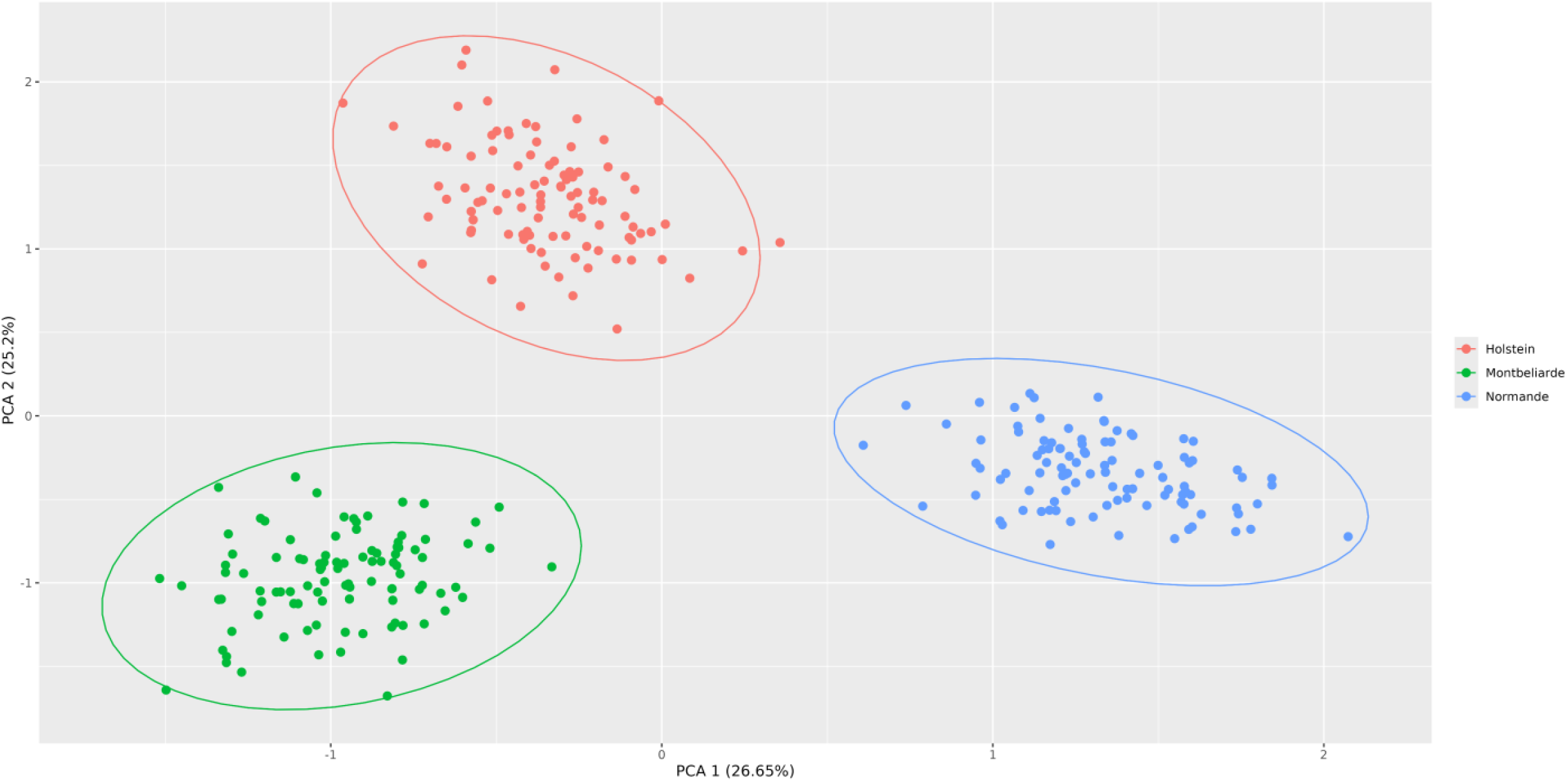
Results of PCA PCA results were shown for the 3 mains dairy breeds, i.e., Holstein, Montbéliarde and Normande

### Genome wide association analyses

GWAS were performed for 13 traits related to milk production and composition, udder health, fertility, and stature, with 58,191 genomic variants, including 230 SVs. We detected a total of 49, 61, and 158 QTL with significant effects on various phenotypes (-log_10_(P) ≥ 6.1) in MON, NMD and HOL bulls’ panel, respectively (Table 7). Specifically, for MON, between 2 and 18 QTL were identified for milk production traits, 4 for stature, 2 for fertility traits and 1 for udder health traits. In the NMD breed, 3 to 19 QTL were found for milk production traits, 1 QTL for HCR, and 13 QTL for SH. Finally, in the HOL panel, from 1 to 48 QTL were found for each trait, except for CM. Overall, milk production traits exhibited the highest number of QTL across the three breeds, with 41, 47, and 130 QTL detected in MON, NMD, and HOL, respectively. In contrast, fewer QTL were found for traits related to fertility (3, 1 and 15, respectively) and udder health (1, 0 and 3, respectively) (Additional file 5: Figures S29-S41).

**Table 7.** Number of QTL identified per breed and phenotype, along with the corresponding number of significant SNPs and SVs.

|  |  | MON |  |  | NMD |  |  | HOL |  |  |
| --- | --- | --- | --- | --- | --- | --- | --- | --- | --- | --- |
| Type of trait |  | #QTL | #SNP | #SV | #QTL | #SNP | #SV | #QTL | #SNP | #SV |
| **Fertility** | Heifer conception rate | 0 | - | - | 1 | 5 | - | 3 | 4 | - |
|  | Cow conception rate | 1 | 1 | - | 0 | - | - | 2 | 5 | - |
|  | Calving-1<sup>st</sup> AI Interval | 2 | 8 | - | 0 | - | - | 8 | 29 | - |
|  | Heifer non-return rate | 0 | - | - | 0 | - | - | 1 | 6 | - |
|  | Cow non-return rate | 0 | - | - | 0 | - | - | 1 | 1 | - |
| **Milk production** | Milk yield (kg) | 3 | 17 | - | 4 | 9 | - | 14 | 94 | - |
|  | Fat content (%) | 2 | 14 | - | 3 | 5 | - | 21 | 133 | - |
|  | Protein content (%) | 2 | 2 | - | 4 | 8 | - | 5 | 8 | - |
|  | Fat yield (kg) | 18 | 93 | - | 17 | 144 | - | 48 | 270 | - |
|  | Protein yield (kg) | 16 | 70 | - | 19 | 120 | - | 42 | 213 | - |
| **Udder health** | Somatic cell score | 0 | - | - | 0 | - | - | 3 | 5 | - |
|  | Clinical mastitis | 1 | 14 | - | 0 | - | - | 0 | - | - |
| **Morphology** | Height at sacrum | 4 | 31 | - | 13 | 68 | - | 10 | 28 | 1 |

SNP Single Nucleotide Polymorphism; SV Structural Variant; MON Montbéliarde; NMD Normande; HOL Holstein

QTL associated with milk production traits were identified on chromosomes 6 and 14 in all three breeds. Specifically, a peak was detected on BTA6 at approximately 85.5 Mb for PY (Additional file 5: Figure S38). On BTA14, a peak was observed around 600 kb for MY and FC, and it was associated with the most significant effect for FY across the three breeds (Additional file 5: Figures S34, S35, and S37). In contrast, QTL associated with fertility, udder health, and morphology traits were located on different chromosomes depending on the breed. For instance, for HCR, no QTL was detected in MON, but peaks were found on BTA19 in NMD, and on BTA6, 7, and 15 in HOL (Additional file 5: Figure S29). Similarly, another fertility trait (*i.e.,* CCR), showed no significant association in NMD but displayed distinct peaks on BTA29 in MON and on BTA14 and 18 in HOL (Additional file 5: Figure S30).

Among genomic variants with significant effects on traits, we identified one SV presenting the second most significant association with height at sacrum in the HOL breed (-log_10_(P) = 19.22) (Fig. 7). This SV corresponds to a 6.2 kb deletion spanning the BTA11:78,819,207-78,825,389 region of the pangenome.

**Figure 7.**
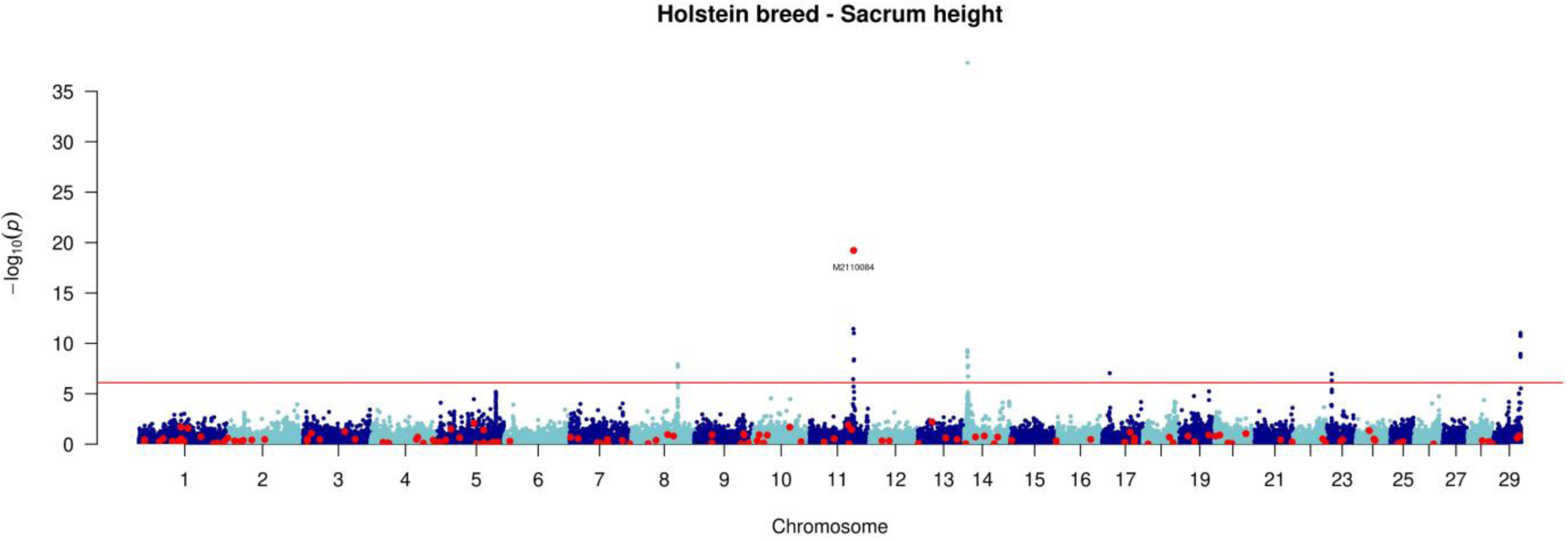
Manhattan plot of GWAS analysis: -log_10_(P) values plotted against the positions of *Bos taurus* autosomes for variants associated with height at sacrum in Holstein bulls Red dots correspond to SVs alongside the genome, blue dots correspond to SNPs.

Analysis of a local pangenome graph within the 2 Mb region surrounding this structural variant revealed the presence of two major paths. The first corresponded to the reference allele, spanning nodes s20, s21, s22, s23, and s24 (Fig. 8a). The second path represented the alternative allele and included a 6.2 kb deletion, spanning nodes s20, s23, and s24 (Fig. 8b). This deletion overlaps with the *MATN3* gene, potentially disrupting its structure. Alignment of *MATN3* gene sequences revealed that the major part of the gene is located within the core pangenome (Fig. 8c). However, the first exon and part of the first intron of the gene are located within the nodes of the flexible genome, suggesting that the 6.2 kb deletion may affect the 5’ UTR regulatory region, as well as the first exon of the *MATN3* gene.

**Figure 8.**
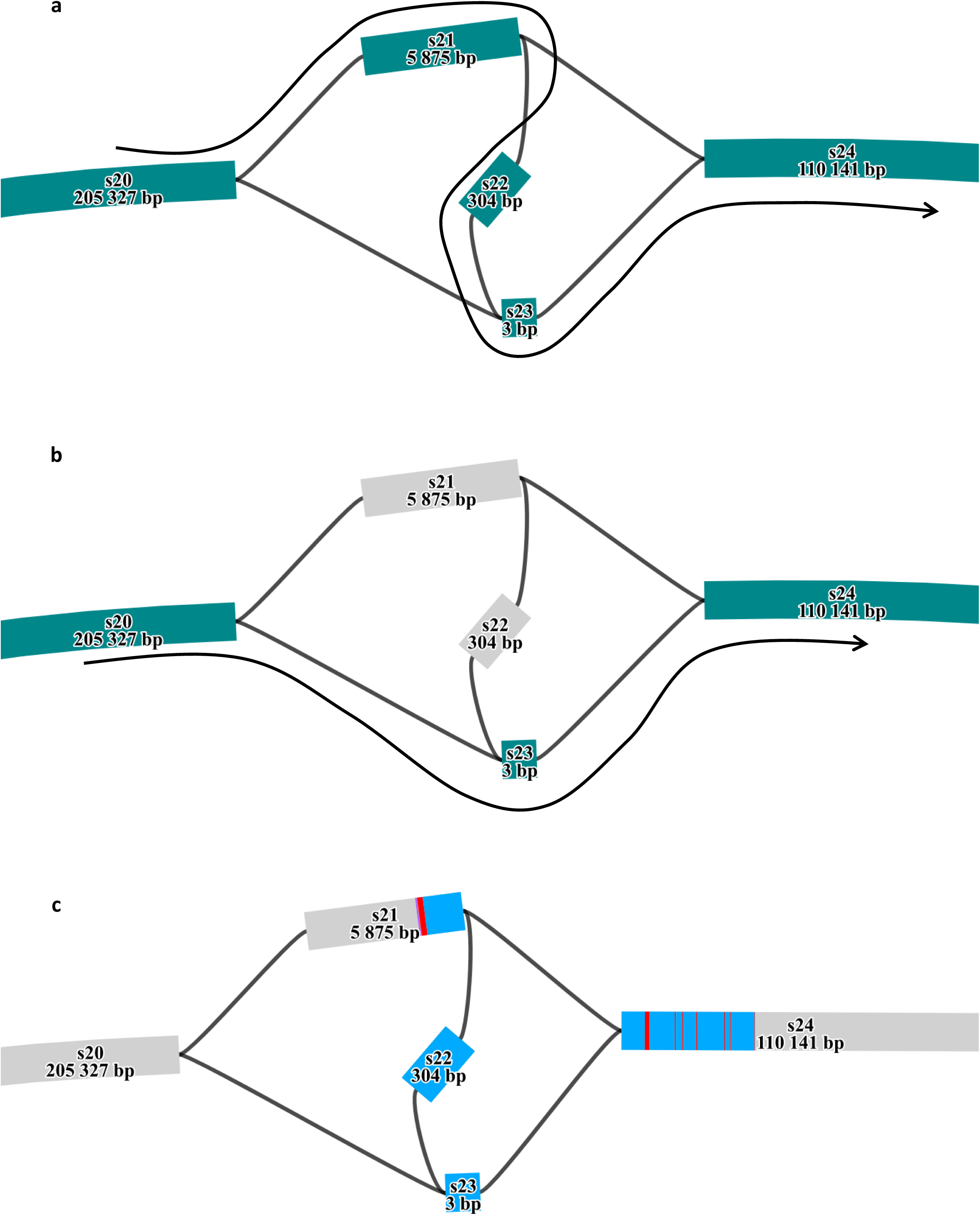
Local pangenome of the *MATN3* region (a) Reference allele path represented by colored nodes, *i.e.* s20, s21, s22, s23 and s24. (b) Alternative allele path represented by colored nodes, and grey nodes were absent from the path, corresponding to the 6.2 kb deletion, *i.e.* s20, s23 and s24. (c) Alignment of MATN3 exons (red bands) and introns (blue bands) on the local pangenome.

### *MATN3* breakpoint validation

The *MATN3* deletion region, identified through GWAS and pangenome structural variant analysis, was amplified from genomic DNA in six individuals, *i.e.* two of each genotype. Gel electrophoresis confirmed the expected amplicon patterns: individuals 1 and 2 carried the reference allele (422 bp), individuals 5 and 6 carried the alternative allele (580 bp), and heterozygous individuals (samples 3 and 4) displayed both fragments (Fig. 9a). PCR products from one reference homozygote (sample 2) and the two alternative homozygotes were sequenced. Sanger sequencing validated the deletion breakpoints at positions 78,819,206 bp and 78,825,386 bp on chromosome 11, occurring between nucleotides G and C (Fig. 9b).

**Figure 9.**
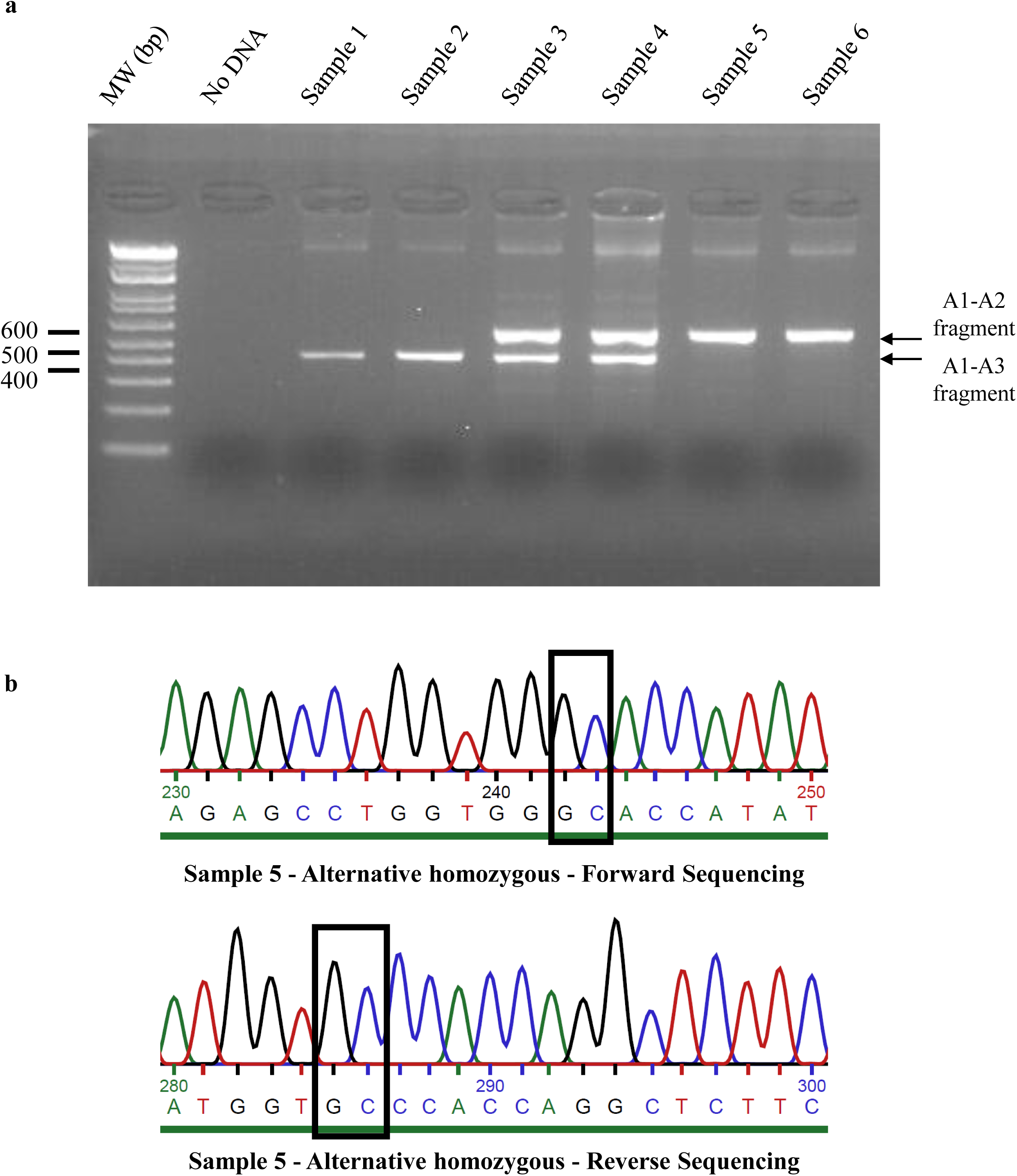
PCR validation of the *MATN3* deletion (a) Gel electrophoresis of A1-A2-A3 PCR products obtained from six different samples. Samples 1 and 2 showed a product around 400 bp, indicating homozygosity for the reference allele. Samples 3 and 4 displayed two bands around 400 and 600 bp, indicating heterozygosity. Samples 5 and 6 presented a product around 600 bp, indicating homozygosity for the alternative allele. (b) Sanger sequencing results for sample 5 showing the breakpoints of the deletion (indicated by black boxes).

## Discussion

The current ARS-UCD1.2 reference genome is a consensus assembly sequence from a single Hereford cow. It thus presents limitations to study the whole spectrum of cattle genetic diversity. In this study, we used 64 *de novo* genome assemblies for 14 dairy and beef cattle breeds to construct a more globally representative cattle pangenome graph. To ensure the construction of a high-quality pangenome graph, we applied three extensive sequence polishing steps on both CLR long reads and high-quality Illumina short reads to correct for a large proportion of small assembly sequence errors. Although slightly shorter than the bovine reference genome, the sizes of these *de novo* genome assemblies are highly comparable to other previously published bovine genome assemblies [39]. Analyses using D-Genies [38] between assemblies and the reference genome assembly also confirm assemblies quality. Phylogenetic tree reconstruction further shows a clear clustering of individuals to their breed of origin.

Our pangenome graph provides new insights into the genetic diversity of the 14 French dairy and beef breeds used in this study and can be considered as a valuable resource for future genomic studies. To our knowledge, this is the first study of this scale in terms of diversity and number of *Bos taurus taurus* assemblies used, allowing the identification of several megabases of novel sequences not present in the ARS-UCD1.2 reference genome. Compared to the findings of Crysnanto *et al.* [7], our pangenome is characterized by a significantly higher number of nodes (507,822 vs. 182,940) and a larger total size (2,858,048,002 bases vs. 2,558,596,439 bases). These differences can be attributed to the inclusion of a larger number of assemblies (64 vs. 6 individuals), as well as a more diverse set (14 vs. 6 breeds), which enabled us to capture a larger part of genomic variation while increasing the structural complexity of the pangenome graph. Similar conclusions were reported by Miao *et al.* [40], who observed an increase in pangenome diversity when utilizing 21 pig assemblies compared to 11 in a previous study [41]. The increased heterogeneity of our assemblies also influences the distribution of sequences between the core and the flexible genomes. Specifically, 90% of the pangenome (2,562,959,040 bases) are conserved across all individuals, while the remaining 10% (295,088,962 bases) correspond to variable regions. This proportion of the flexible genome is higher than previously reported, where the core genome accounted for 93.9% (2,402,561,410 bases) and 95.8% (2,598,811,581 bases) of the total pangenome size, compared to 6.1% (156,035,029 bases) and 4.2% (109 Mb) of variable sequences, in Crysnanto *et al.* [7] and Leonard *et al.* [42] studies, respectively.

In total, we identified 151,771 nodes representing 158.7 Mb absent from the cattle reference genome. This exceeds the amount of NRSs reported in previous studies, which identified 70.3 Mb in *Bos taurus* [7], 18.6 Mb [43] and 148.5 Mb [44] in *Bos indicus*, 38.3 Mb in goats [45], and 72.5 Mb [46] and 105.16 Mb [40] in pigs, based on pangenome graphs constructed from 6, 5, 20, 8, 11, and 21 assemblies, respectively. However, our dataset includes a substantially larger number of individuals, which likely accounts for the higher amount of identified NRSs. These findings are consistent with the patterns observed in our pangenome graph, particularly regarding breed-specific NRUIs. Indeed, we observed a positive correlation between number of individuals per breed and total NRUI length.

For instance, Holstein cattle (8 individuals) exhibited a total of 1.73 Mb of NRUIs, whereas Limousine and Rouge Flamande breeds (2 individuals each) showed lower values of 0.42 Mb and 0.62 Mb, respectively.

From the pangenome graph, we identified 109,275 SVs, which is higher than SV catalogs previously reported. For example, a catalog based on 16 HiFi cattle haplotype-resolved assemblies detected 53,297 SVs [47], and another study identified 68,328 SVs from six cattle assemblies [7]. In our SV catalog, 84,612 SVs (77%) were characterized as bi-allelic, with 21,840 insertions and 21,340 deletions. The distribution of insertion and deletion sizes revealed a symmetrical pattern between these two categories, with a majority of small size (between 50 and 200 bp) SVs, and a decreasing number of SVs as size increased. Additionally, four peaks (150 bp, 250 bp, 5.5 kb and 8.6 kb) were observed and corresponded to the SVs caused by transposable elements. These results are consistent with previous studies, such as those reported by Eche *et al.* [39]. Moreover, the proportion of bi-allelic SVs in our study (77%) is lower than the 94% (64,224 SVs) reported by Crysnanto *et al.* [7]. This difference can be attributed to the inclusion of a larger number of assemblies from diverse cattle breeds in our pangenome, which increases the likelihood that some SVs display more than two alleles, thus classifying them as multi-allelic. Further analysis of the insertions revealed 25.4 Mb of NRUIs, that are of particular interest as they may code for potential functional elements.

Inspection of this SV panel revealed the presence of several known SVs related to phenotypic traits. For example, we identified the 8.4 kb LINE1 insertion at the *ASIP* locus on BTA13 which encodes the AGOUTI signalling protein involved in mammalian pigmentation. This SV was found exclusively in all the Normande animals and is known to be associated with coat colour variation [48].

As nine out the 230 SVs were found monomorphic in our study, we used the genotyping data to investigate the effect of the 221 remaining polymorphic SVs by conducting GWAS analyses in the three main French dairy breeds for 13 phenotypes related to fertility, milk production, udder health, and morphology. We identified numerous QTL associated with the analysed phenotypes, many of them have previously been reported in other studies [49–51]. In most cases, candidate genes were highlighted, particularly for milk production and composition. Among them, the well-known *DGAT1* gene (*diacylglycerol O-acyltransferase 1*) that encodes an enzyme catalysing the synthesis of triglycerides in milk, was identified on BTA14 (∼600 kb) [52,53]. Similarly, the cluster of genes encoding αs1-casein (*CSN1S1*), αs2-casein (*CSN1S2*), β-casein (*CSN2*), and κ-casein (*CSN3*) was detected on BTA6 (∼85.5 Mb) and is strongly associated with milk protein content [54,55]. Additionally, *MGST1* (*microsomal glutathione S-transferase 1*), located on BTA5 (∼93.5 Mb), has been found associated with milk fat content [56,57]. However, identifying the causal mutation underlying a candidate gene remains a major challenge and is not systematically determined. To date, most GWAS have focused on SNPs or InDels, while SVs, which represent a substantial portion of genetic and phenotypic variability [47,58], remain largely unexplored, particularly in cattle.

In this context, our study aimed to better characterize the impact of a SV panel on phenotypes of interest, providing new insights into their role in the genetic architecture of complex traits. Our results highlight a major QTL located on BTA11, associated with stature, where the most significant variant is an SV, *i.e.*, a 6.2 kb deletion located in the upstream region and the first exon of the *MATN3* gene. This finding represents a promising step toward integrating SVs into GWAS for phenotype analysis. Its significance is further underscored by the fact that our study used only a subset of the SVs detected from the pangenome (230 SVs out of 84,612 bi-allelic SVs). This observation suggests that leveraging a more extensive SV panel could enhance the power to detect loci and potentially identify causal mutations underlying agronomically relevant phenotypes in GWAS.

Several studies have investigated the genetic determinism of stature in cattle, including a large-scale meta-analysis encompassing 17 cattle populations from nine countries [59]. This study identified 163 genomic regions of 1 Mb with significant effects on this phenotype, eight of which were located on chromosome 11. Among these, a SNP at position 78,870,305 bp (reference genome: UMD 3.1) exhibited the strongest association with stature (-log_10_(P) = 42.89) but no candidate gene was identified at this position. Our results confirm the involvement of this region in the genetic determinism of height at sacrum in cattle. The SV detected in our study is located at 78,825,400 bp, in close proximity to this region. Moreover, we identify the *MATN3* candidate gene that may be affected by the presence of this SV. Several other studies have also reported significant QTLs in this genomic region (BTA11 ∼78 Mb), two mentioning the *MATN3* gene as a positional candidate gene [60,61]. However, no functional validation has yet been conducted to confirm its biological role. Nonetheless, these findings further support our GWAS results and suggest a functional impact of this region, warranting further investigations. This also highlights that adding SVs into GWAS analyses can provide a better understanding of the genetic determinism of complex traits.

*MATN3* is part of the matrilin genes family and has been widely studied in recent years. This gene encodes the protein matrilin 3, which is primarily expressed in cartilage tissue and plays a key role in extracellular matrix assembly [62]. Due to its role in the collagen development, studies have shown that mutations in *MATN3* are associated with a predisposition to osteoarthritis and the premature development of growth plate chondrocytes in mice [62]. Additionally, a *MATN3* mutation has been associated to spondylo-epi-metaphyseal dysplasia and dwarfism in human [64].

In our study, the identified SV corresponds to a 6.2 kb deletion that affects the first exon and half of the first intron of *MATN3*. Analysis of the *MATN3* coding protein indicates the presence of three transcript isoforms in Ensembl [31], all of which consistently include exon 1 in the translated protein.

Concordance between Ensembl and Uniprot [65] data further confirms that the functional *MATN3* protein contains exon 1. Given the structural variant’s location, it is likely that this deletion also affects the 5’ UTR region of the gene and its associated enhancer. Moreover, the cattle Genotype-Tissue Expression atlas (CattleGTEx) [66] database does not report any isoform for the *MATN3* protein, providing no evidence of an impact of the identified deletion in our study. However, this absence may be due to sample limitations, as the *MATN3* gene is primarily expressed in cartilage tissue during development stages, as previously described.

Thus, in addition to characterizing the pangenome and identifying SVs and NRUIs in French cattle breeds, our study identifies a positional and functional candidate SV associated with stature in the Holstein breed. However, additional functional validation analyses are required to confirm the involvement of this variant in the genetic determinism of this trait.

## Conclusions

Numerous studies have underscored the value of pangenome graphs in identifying large structural variations often missed when relying on a single linear reference genome. Our findings corroborate these observations by using 64 *de novo* assemblies from 14 French dairy and beef cattle breeds to construct a comprehensive cattle pangenome. This approach enabled the identification of an extensive catalog of structural variations and non-reference sequences. By integrating some of these SV into GWAS analyses, we detected a 6.2 kb deletion in the *MATN3* gene, strongly associated with stature in Holstein cattle. This study emphasizes the importance of incorporating pangenome-based approaches in genetic studies to better capture variants that may contribute to key phenotypic traits in cattle.

## Supporting information

Access_to_supplemental_files

## Declarations

### Ethics approval and consent to participate

Not applicable

### Consent for publication

Not applicable

### Availability of data and materials

The fasta files of the 64 assemblies used to build the pangenome are available in the European Nucleotide Archive (ENA) at EMBL-EBI under accession number PRJEB68295 (https://www.ebi.ac.uk/ena/data/view/PRJEB68295). Individual assembly accession numbers are available in the Additional file 1 Table S1. Additionally, paired-end Illumina SR data (2 × 150 bp) are available for the same 64 animals and publicly accessible under ENA project accession number PRJEB64023.

## Authors’ contributions

V.S., M.B., M-P.S., G.T-K., D.B. and LD conceived the study; V.S. conducted the analysis and wrote the original draft; M.B., M-P.S., G.TK. and L.D. contributed to writing the manuscript; Ce.G., S.F. and C.E. collected samples and realized extraction; Ce. G supervised PCR validation of breakpoints; M-M.N. performed population structure analysis; C.E., C.M., A.S., C.D., Ch.G., C.I. and D.M. conceived the experimental design and supervised the technical aspects of the project; C.B. and C.K. generated assemblies; All authors reviewed the manuscript.

## Acknowledgements

We are grateful to the Genotoul bioinformatics platform Toulouse Occitanie (Bioinfo Genotoul, https://doi.org/10.15454/1.5572369328961167E12) for providing computing and storage resources.

We thank Claire Kuchly and Caroline Vernette for their help to submit the de novo genome assembly sequences in the ENA database.

