## Supplementary material for "Application of a French cattle pangenome, from structural variant discovery to association studies on key phenotypes": Access_to_supplemental_files: Additional file 2 Figures S1-S14.pdf

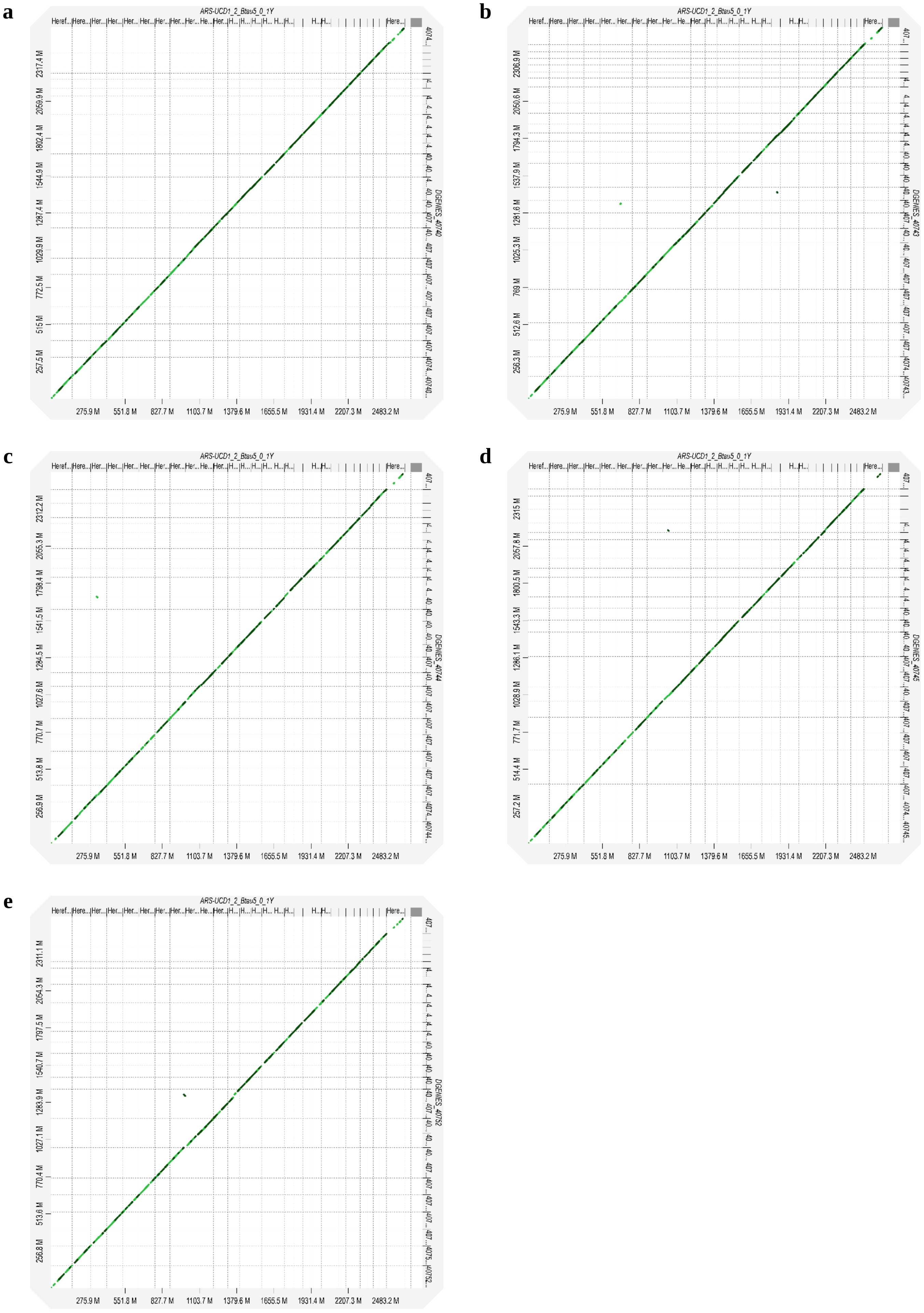


**Figure S1** Chromosomal alignment concordance between ARS-UCD1.2 on *x*-axis and Abondance assemblies on *y*-axis: **a** 40740, **b** 40743, **c** 40744, **d** 40745, and **e** 40752

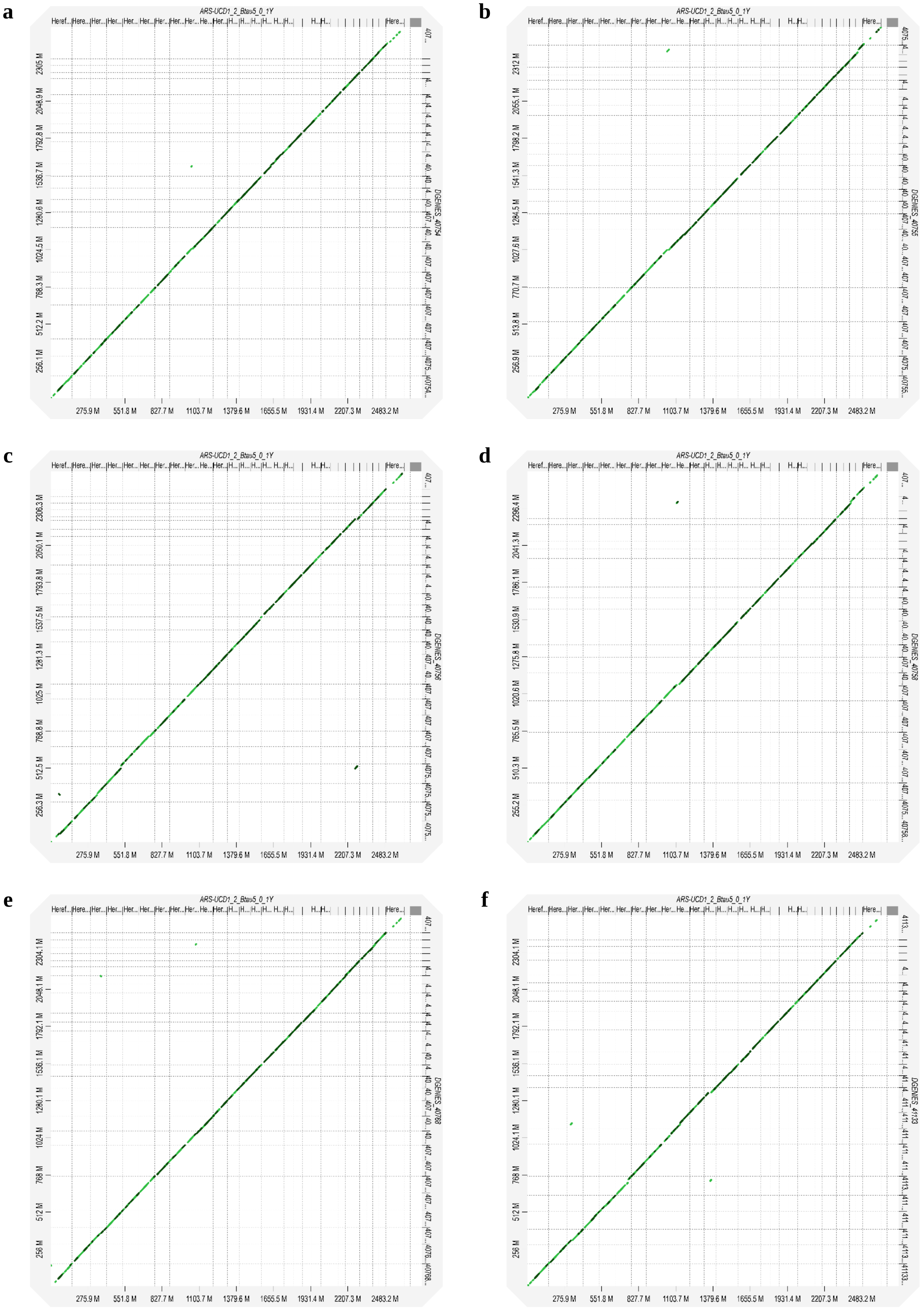

a
b
c
d
e
f
ARS-UCD1_2_Btau5_0_1Y
DGENES_40754
DGENES_40755
DGENES_40756
DGENES_40758
DGENES_40768
DGENES_41133
275.9 M
551.8 M
827.7 M
1103.7 M
1379.6 M
1655.5 M
1931.4 M
2207.3 M
2483.2 M

**g**

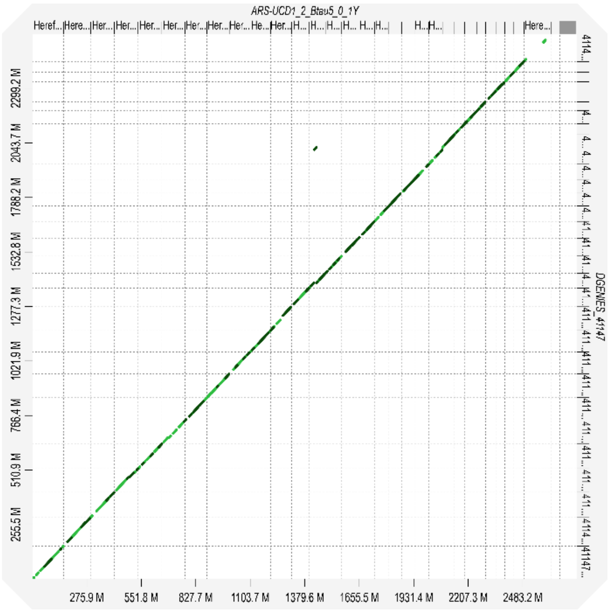


**Figure S2** Chromosomal alignment concordance between ARS-UCD1.2 on *x*-axis and Aubrac assemblies on *y*-axis: **a** 40754, **b** 40755, **c** 40756, **d** 40758, **e** 40768, **f** 41133, and **g** 41147

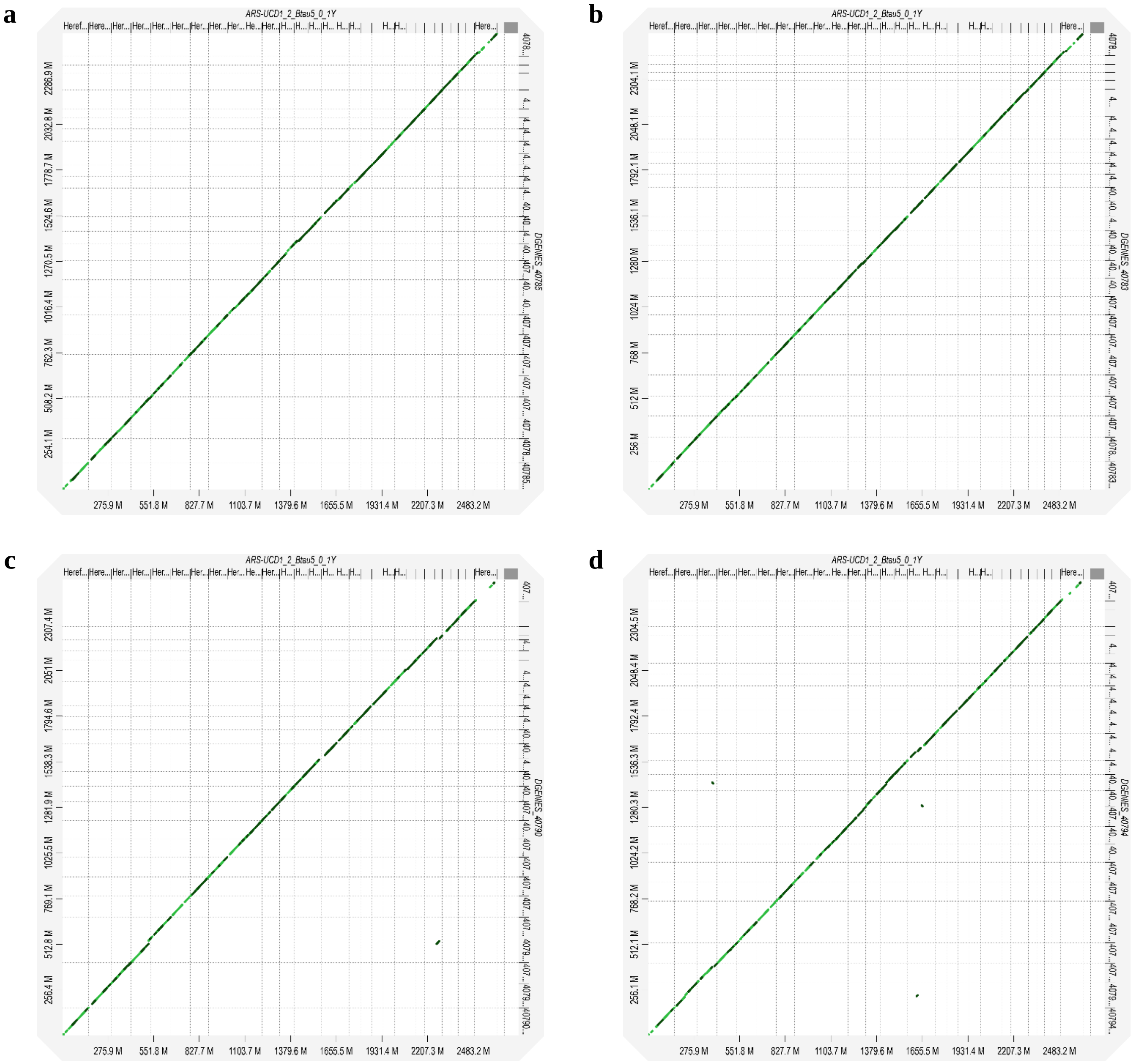


**Figure S3** Chromosomal alignment concordance between ARS-UCD1.2 on *x*-axis and Blonde d'Aquitaine assemblies on *y*-axis: **a** 40783, **b** 40785, **c** 40790, and **d** 40794

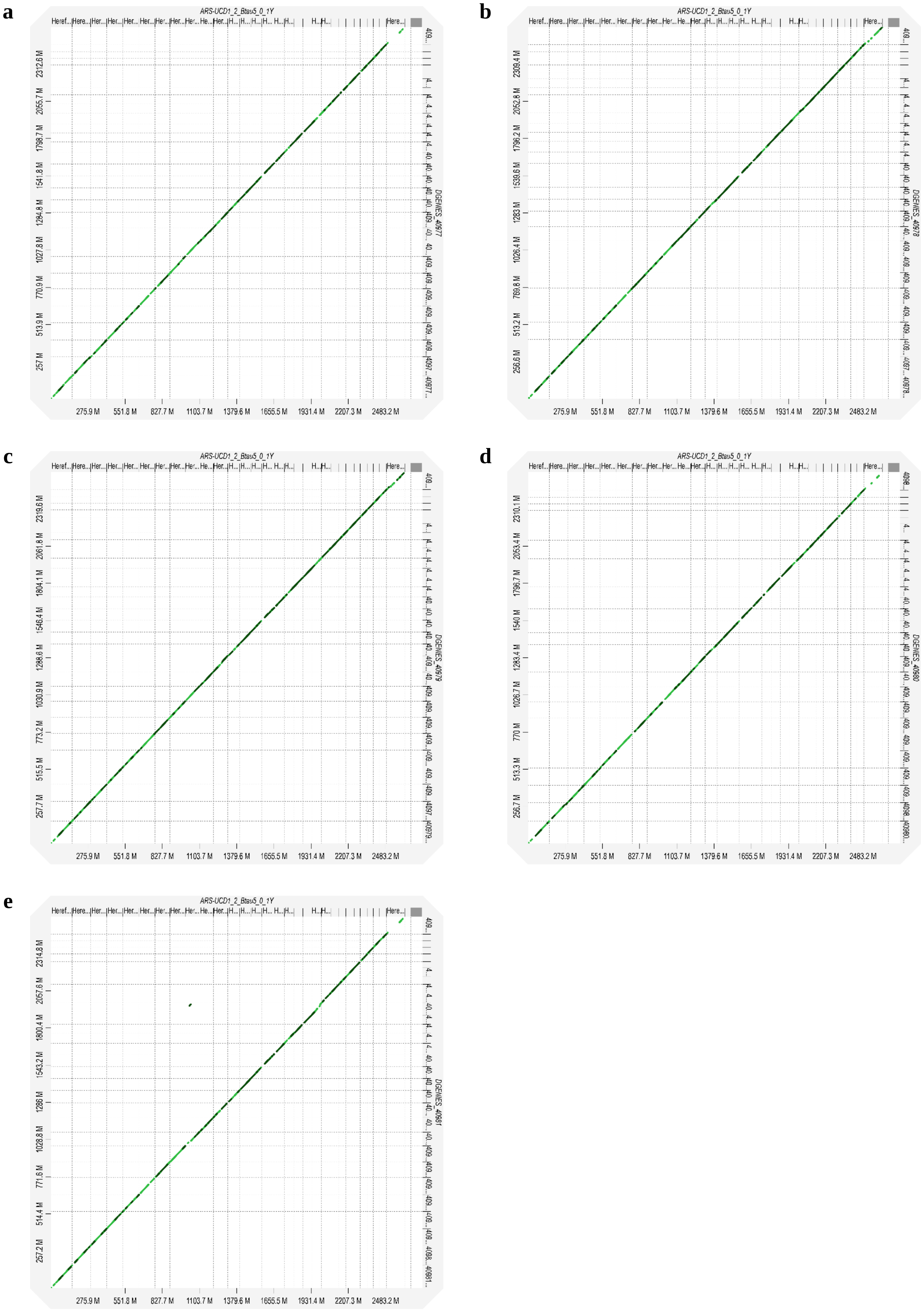


**Figure S4** Chromosomal alignment concordance between ARS-UCD1.2 on *x*-axis and Brown Swiss assemblies on *y*-axis: **a** 40977, **b** 40978, **c** 40979, **d** 40980, and **e** 40981

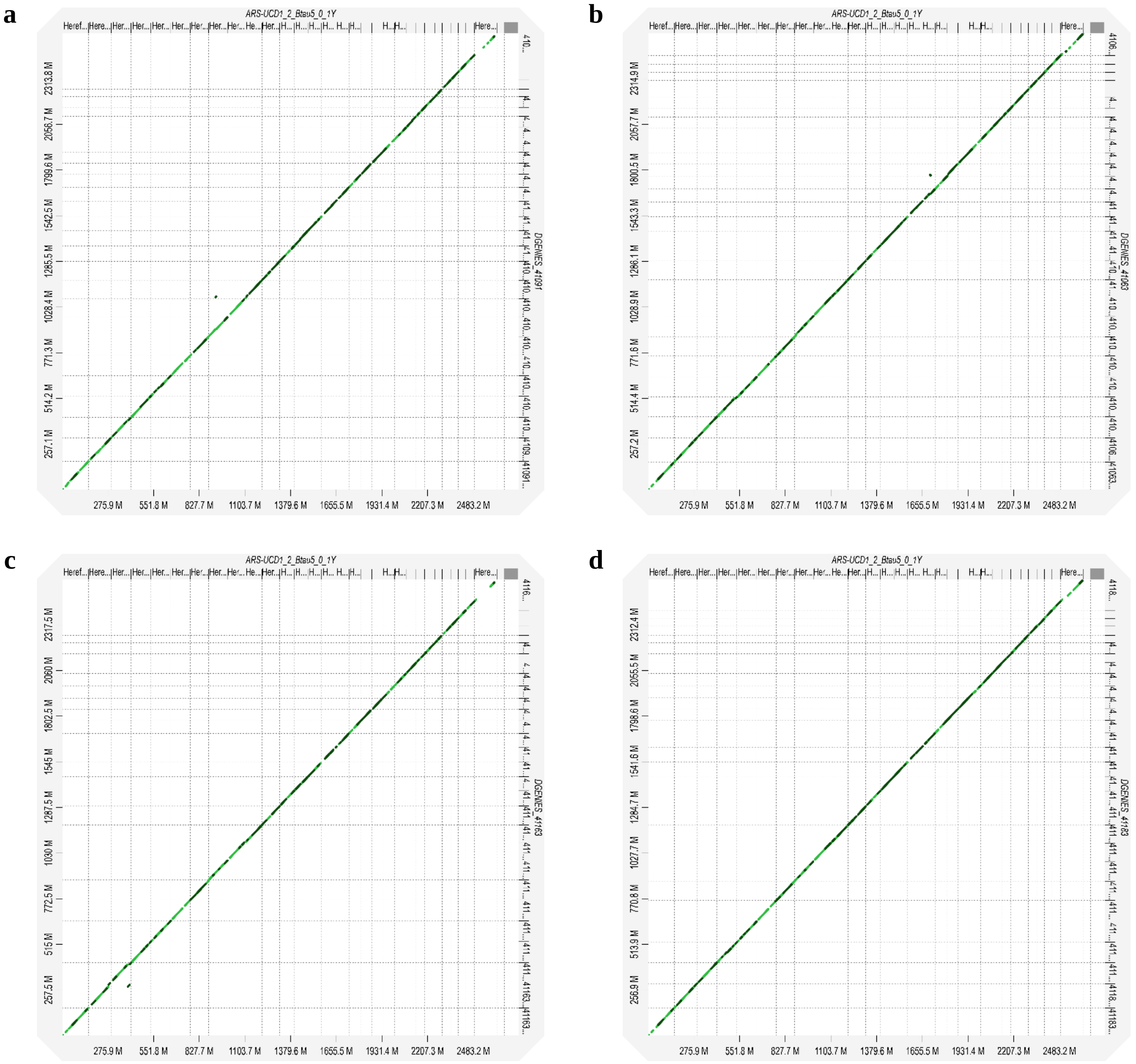


**Figure S5** Chromosomal alignment concordance between ARS-UCD1.2 on *x*-axis and Charolaise assemblies on *y*-axis: **a** 41063, **b** 41091, **c** 41163, and **d** 41183

**a**

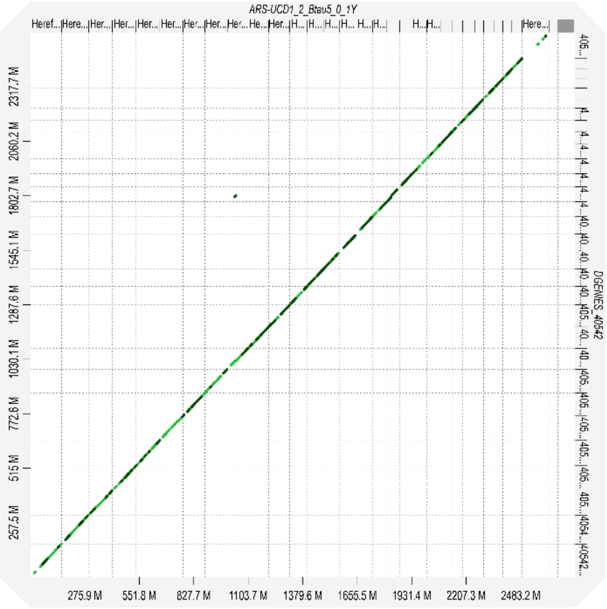
ARS-UCD1_2_Btau5_0_1Y
DGENES_40542
2317.7 M 2060.2 M 1802.7 M 1545.1 M 1287.6 M 1030.1 M 772.6 M 515 M 257.5 M
275.9 M 551.8 M 827.7 M 1103.7 M 1379.6 M 1655.5 M 1931.4 M 2207.3 M 2483.2 M

**b**

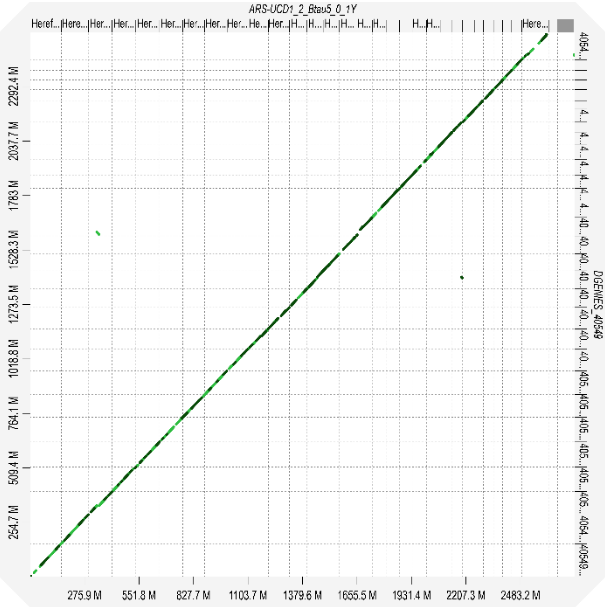
ARS-UCD1_2_Btau5_0_1Y
DGENES_40549
2292.4 M 2037.7 M 1783 M 1528.3 M 1273.5 M 1018.8 M 764.1 M 509.4 M 254.7 M
275.9 M 551.8 M 827.7 M 1103.7 M 1379.6 M 1655.5 M 1931.4 M 2207.3 M 2483.2 M

**c**

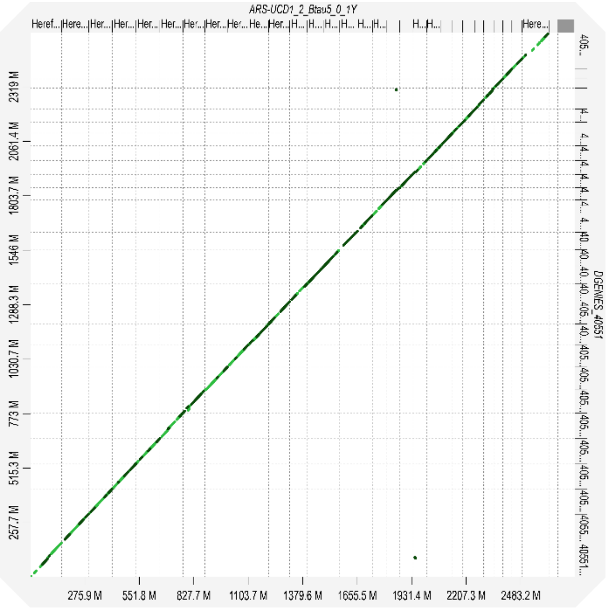
ARS-UCD1_2_Btau5_0_1Y
DGENES_40551
2319 M 2061.4 M 1803.7 M 1546 M 1288.3 M 1030.7 M 773 M 515.3 M 257.7 M
275.9 M 551.8 M 827.7 M 1103.7 M 1379.6 M 1655.5 M 1931.4 M 2207.3 M 2483.2 M

**d**

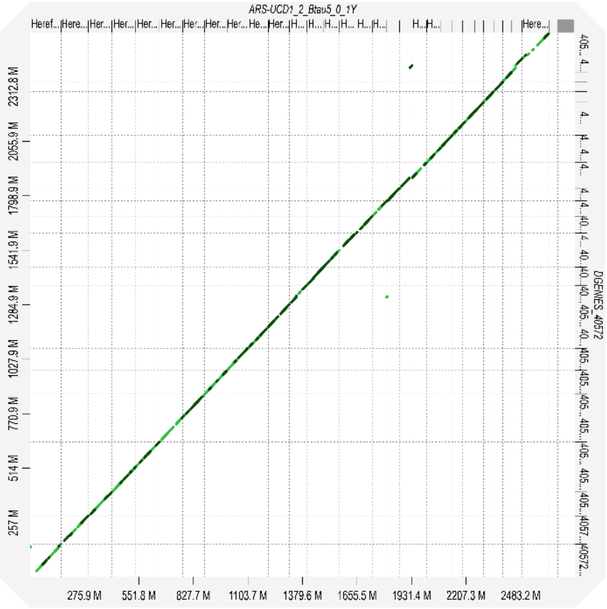
ARS-UCD1_2_Btau5_0_1Y
DGENES_40572
2312.8 M 2055.9 M 1798.9 M 1541.9 M 1284.9 M 1027.9 M 770.9 M 514 M 257 M
275.9 M 551.8 M 827.7 M 1103.7 M 1379.6 M 1655.5 M 1931.4 M 2207.3 M 2483.2 M

**e**

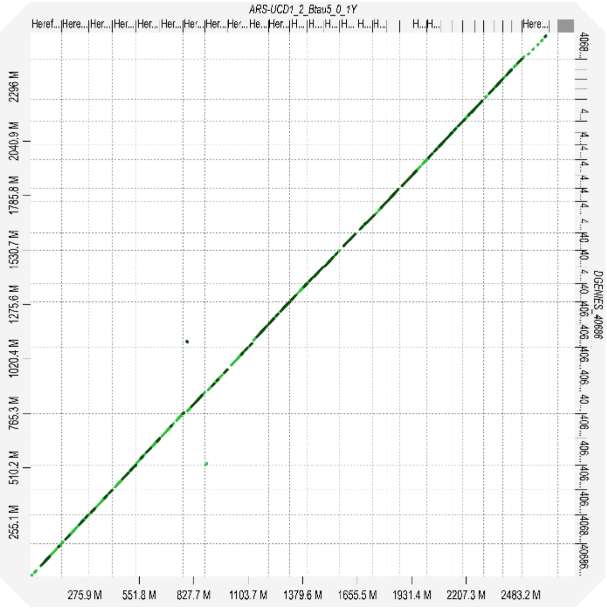
ARS-UCD1_2_Btau5_0_1Y
DGENES_40686
2296 M 2040.9 M 1785.8 M 1530.7 M 1275.6 M 1020.4 M 765.3 M 510.2 M 255.1 M
275.9 M 551.8 M 827.7 M 1103.7 M 1379.6 M 1655.5 M 1931.4 M 2207.3 M 2483.2 M

**f**

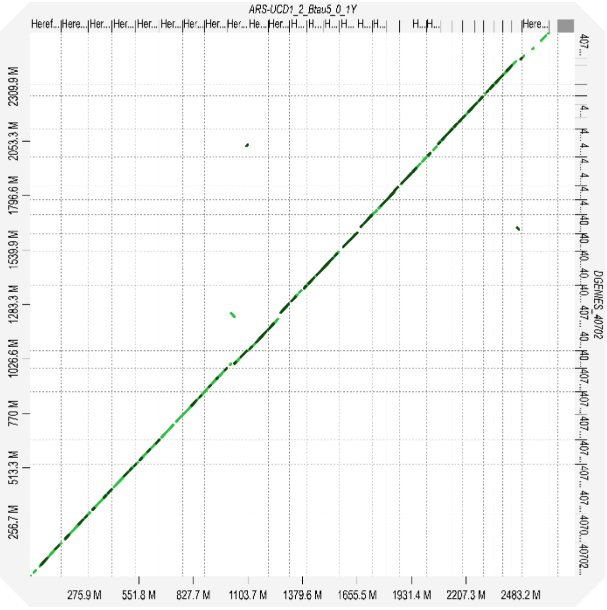
ARS-UCD1_2_Btau5_0_1Y
DGENES_40702
2309.9 M 2053.3 M 1796.6 M 1539.9 M 1283.3 M 1026.6 M 770 M 513.3 M 256.7 M
275.9 M 551.8 M 827.7 M 1103.7 M 1379.6 M 1655.5 M 1931.4 M 2207.3 M 2483.2 M

**g** **h**

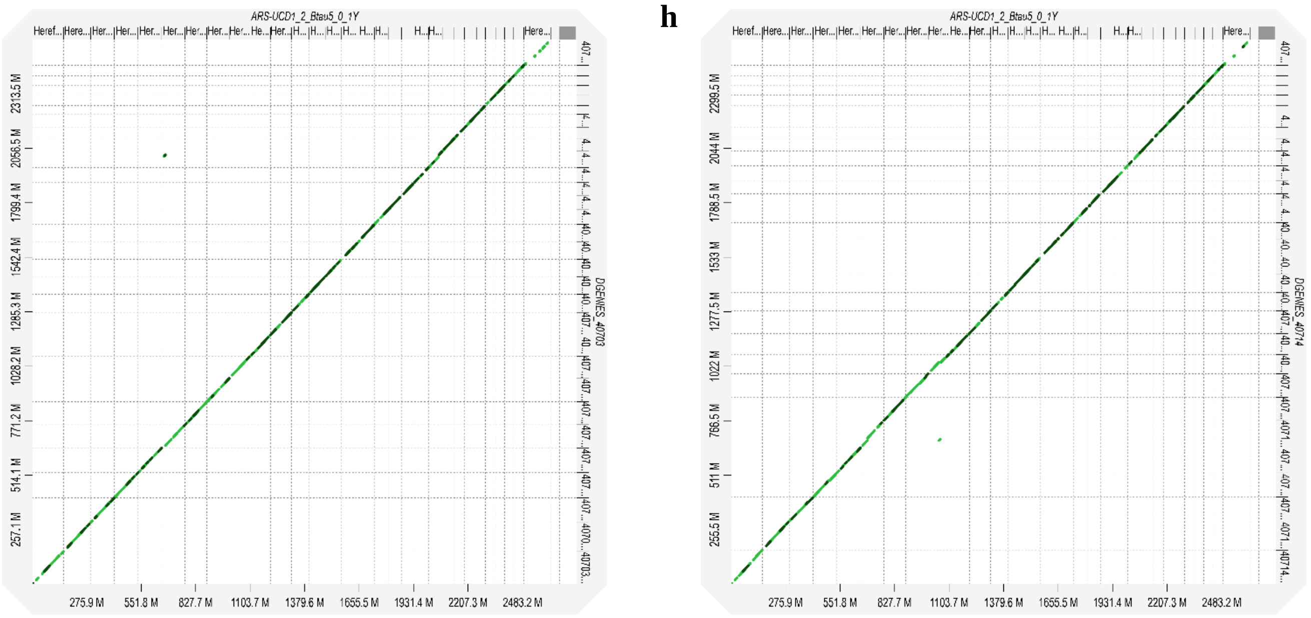


**Figure S6** Chromosomal alignment concordance between ARS-UCD1.2 on *x*-axis and Holstein assemblies on *y*-axis: **a** 40542, **b** 40549, **c** 40551, **d** 40572, **e** 40686, **f** 40702, **g** 40703, and **h** 40714

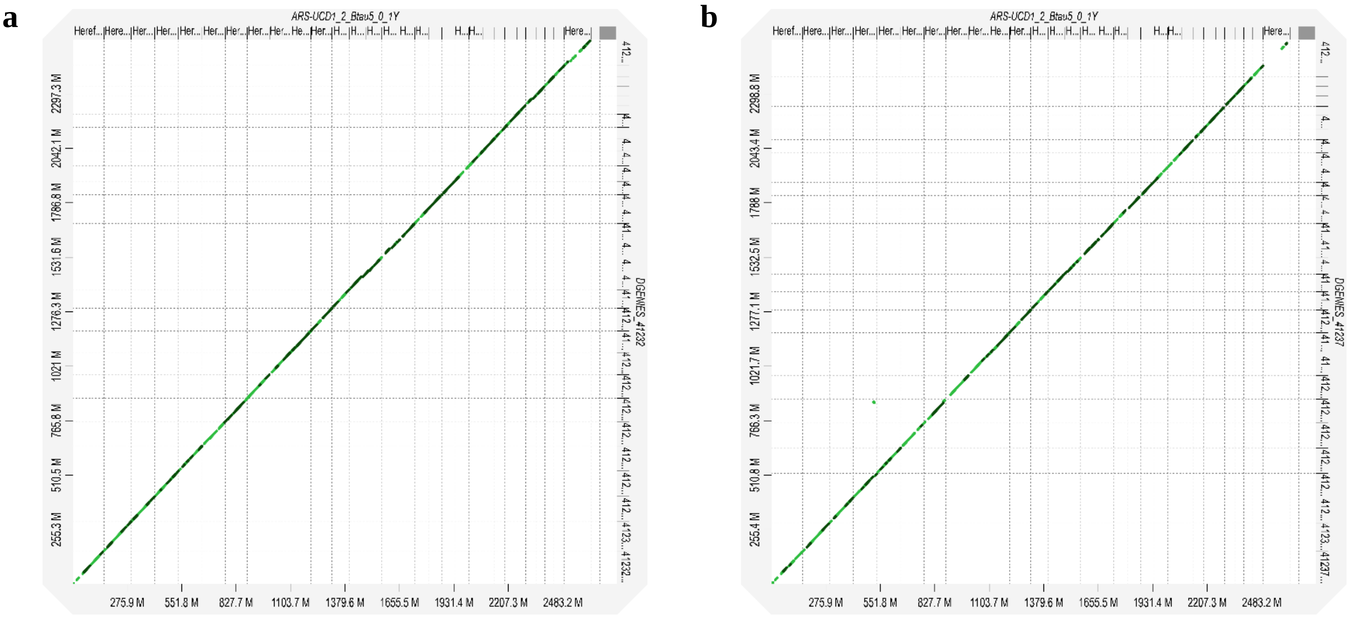


**Figure S7** Chromosomal alignment concordance between ARS-UCD1.2 on *x*-axis and Limousine assemblies on *y*-axis: **a** 41232, and **b** 41237

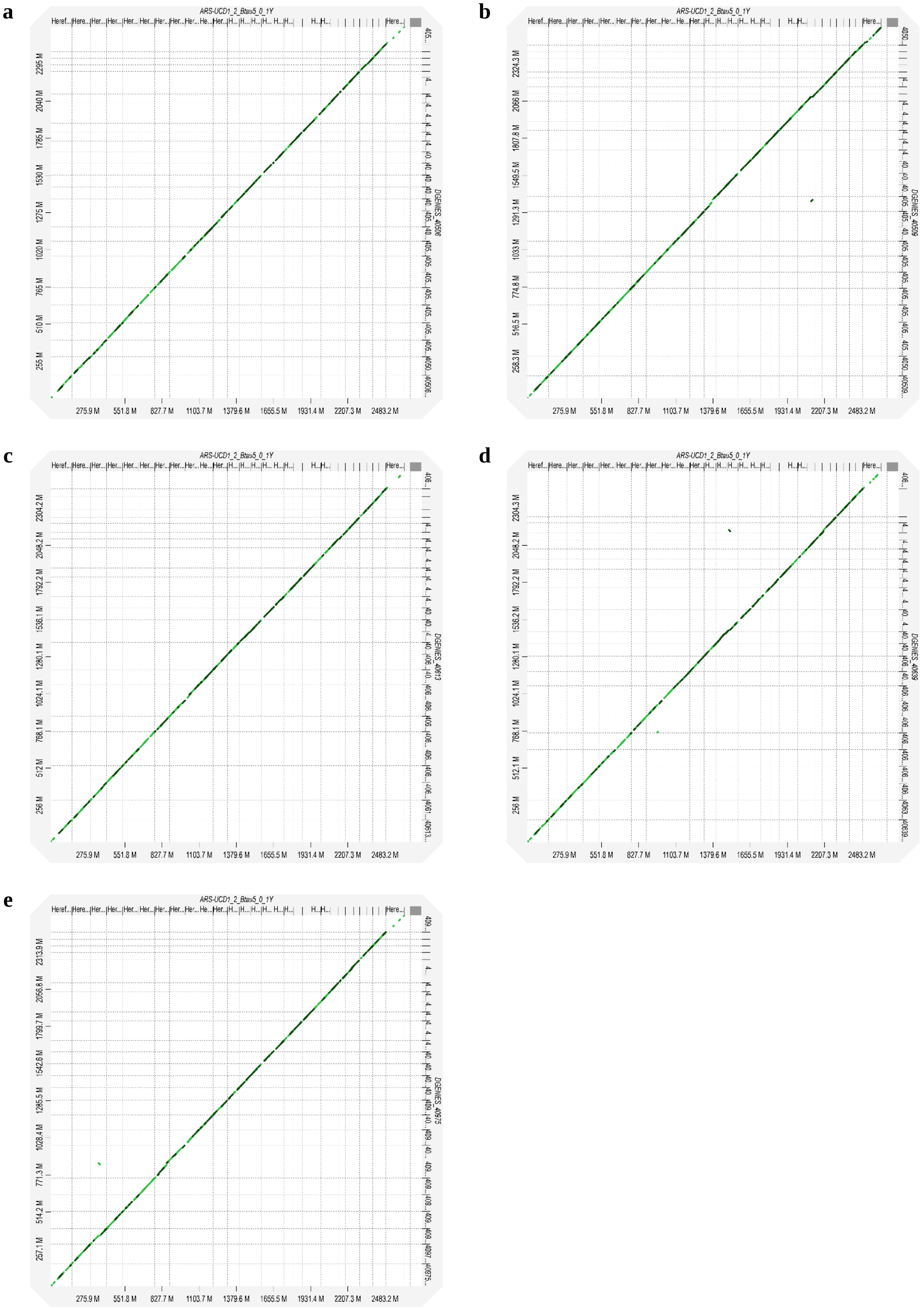


**Figure S8** Chromosomal alignment concordance between ARS-UCD1.2 on *x*-axis and Montbéliarde assemblies on *y*-axis: **a** 40506, **b** 40509, **c** 40613, **d** 40639, and **e** 40975

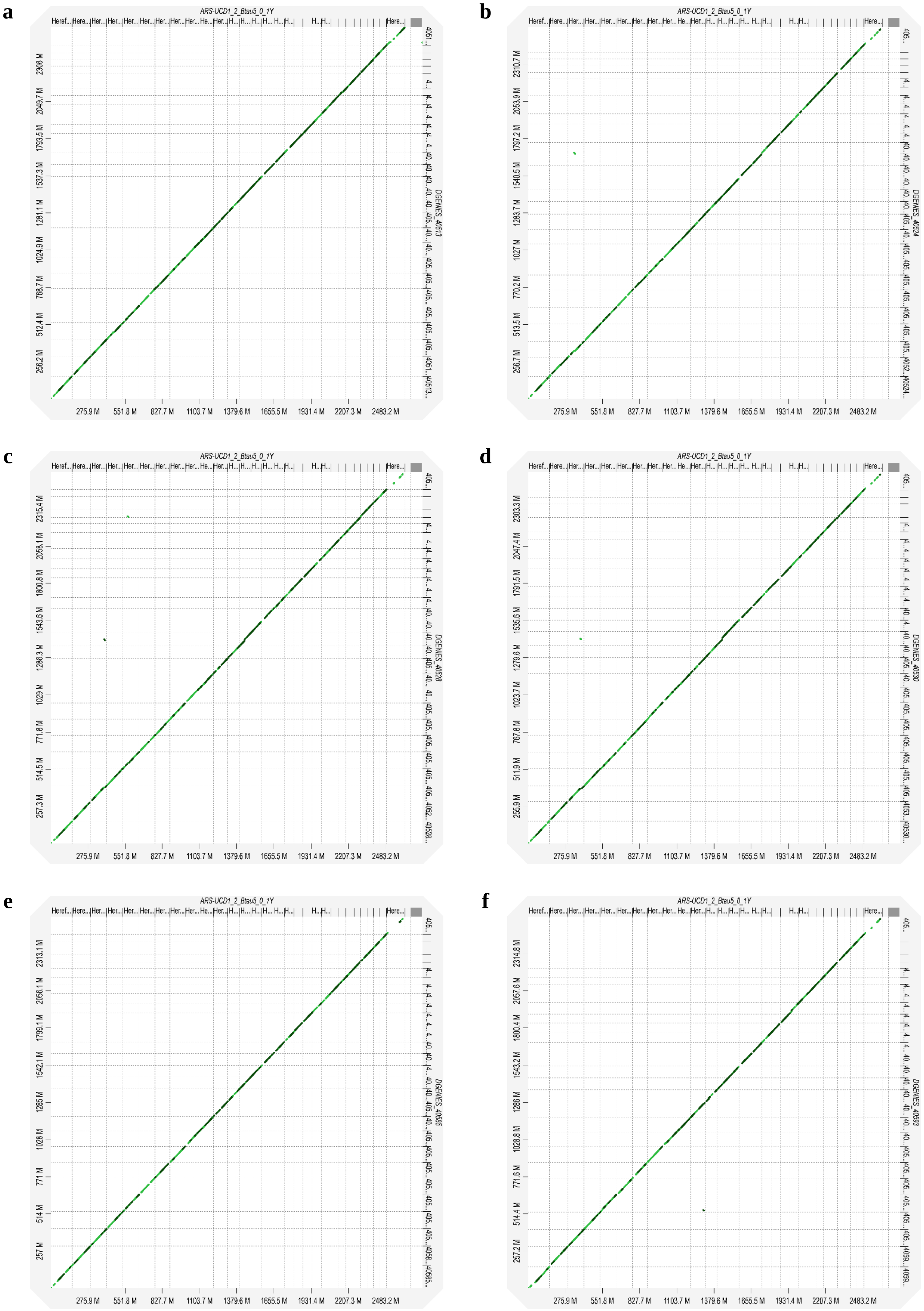
a
ARS-UCD1_2_Btau5_0_1Y
DGENES_40513
2306 M
2049.7 M
1793.5 M
1537.3 M
1281.1 M
1024.9 M
768.7 M
512.4 M
256.2 M
275.9 M
551.8 M
827.7 M
1103.7 M
1379.6 M
1655.5 M
1931.4 M
2207.3 M
2483.2 M
b
ARS-UCD1_2_Btau5_0_1Y
DGENES_40524
2310.7 M
2053.9 M
1797.2 M
1540.5 M
1283.7 M
1027 M
770.2 M
513.5 M
256.7 M
275.9 M
551.8 M
827.7 M
1103.7 M
1379.6 M
1655.5 M
1931.4 M
2207.3 M
2483.2 M
c
ARS-UCD1_2_Btau5_0_1Y
DGENES_40528
2315.4 M
2058.1 M
1800.8 M
1543.6 M
1286.3 M
1029 M
771.8 M
514.5 M
257.3 M
275.9 M
551.8 M
827.7 M
1103.7 M
1379.6 M
1655.5 M
1931.4 M
2207.3 M
2483.2 M
d
ARS-UCD1_2_Btau5_0_1Y
DGENES_40530
2303.3 M
2047.4 M
1791.5 M
1535.6 M
1279.6 M
1023.7 M
767.8 M
511.9 M
255.9 M
275.9 M
551.8 M
827.7 M
1103.7 M
1379.6 M
1655.5 M
1931.4 M
2207.3 M
2483.2 M
e
ARS-UCD1_2_Btau5_0_1Y
DGENES_40585
2313.1 M
2056.1 M
1799.1 M
1542.1 M
1285 M
1028 M
771 M
514 M
257 M
275.9 M
551.8 M
827.7 M
1103.7 M
1379.6 M
1655.5 M
1931.4 M
2207.3 M
2483.2 M
f
ARS-UCD1_2_Btau5_0_1Y
DGENES_40593
2314.8 M
2057.6 M
1800.4 M
1543.2 M
1286 M
1028.8 M
771.6 M
514.4 M
257.2 M
275.9 M
551.8 M
827.7 M
1103.7 M
1379.6 M
1655.5 M
1931.4 M
2207.3 M
2483.2 M

g

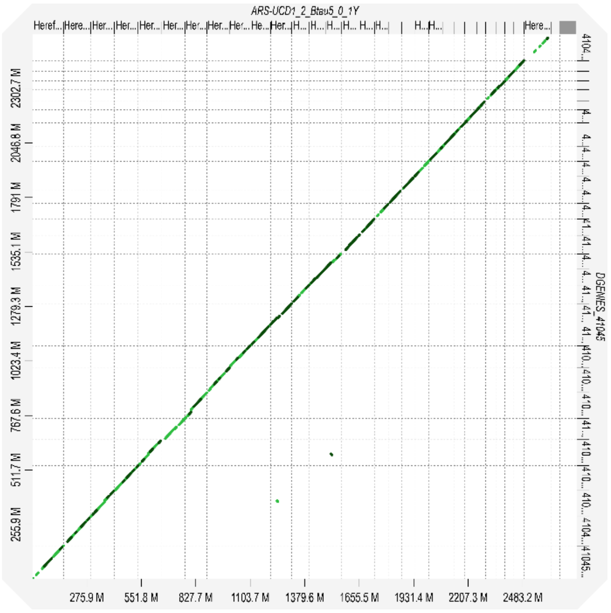


**Figure S9** Chromosomal alignment concordance between ARS-UCD1.2 on *x*-axis and Normande assemblies on *y*-axis: **a** 40513, **b** 40524, **c** 40528, **d** 40530, **e** 40585, **f** 40593, and **g** 41045

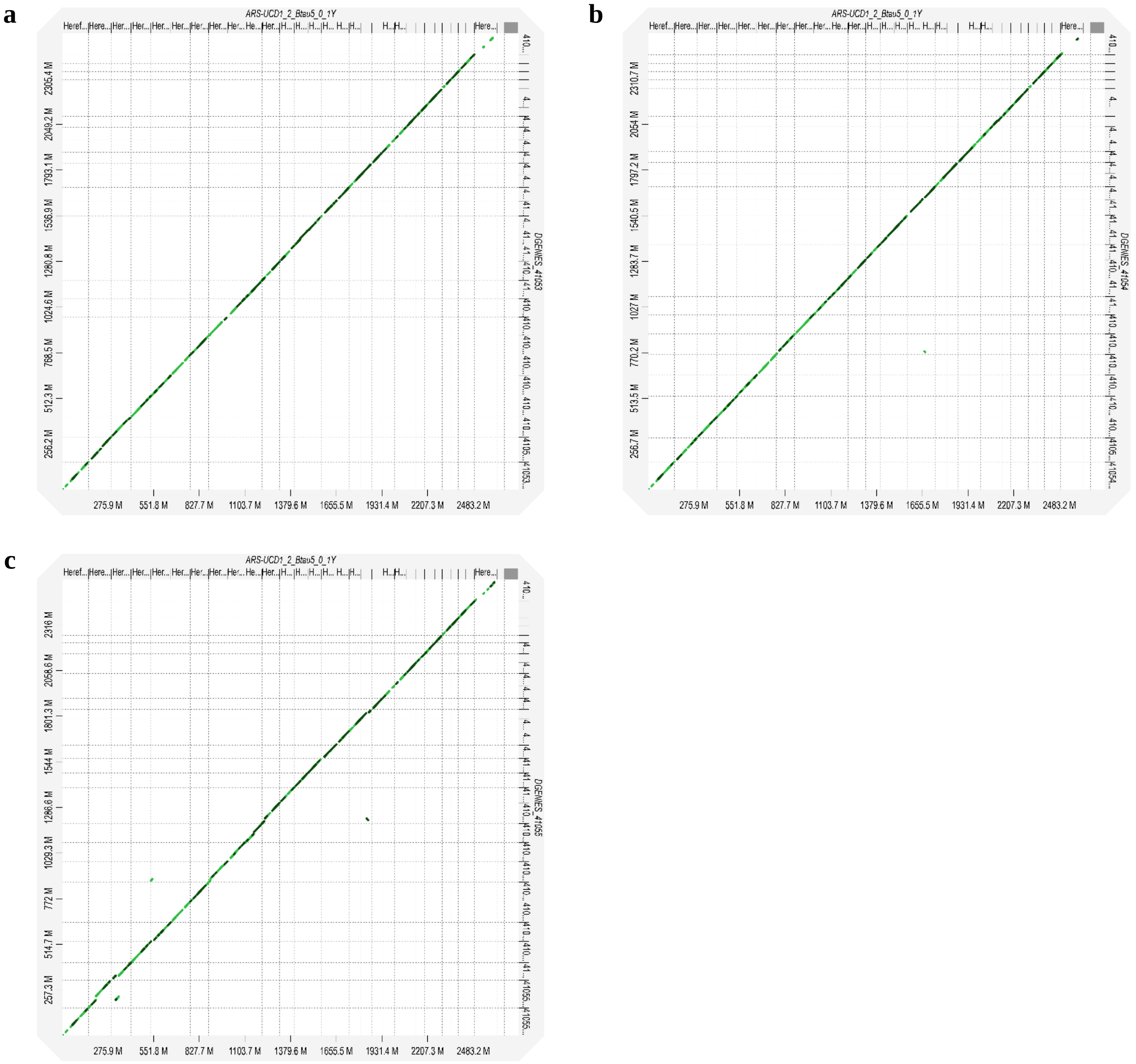


**Figure S10** Chromosomal alignment concordance between ARS-UCD1.2 on *x*-axis and Parthenaise assemblies on *y*-axis: **a** 41053, **b** 41054, and **c** 41055

a

b

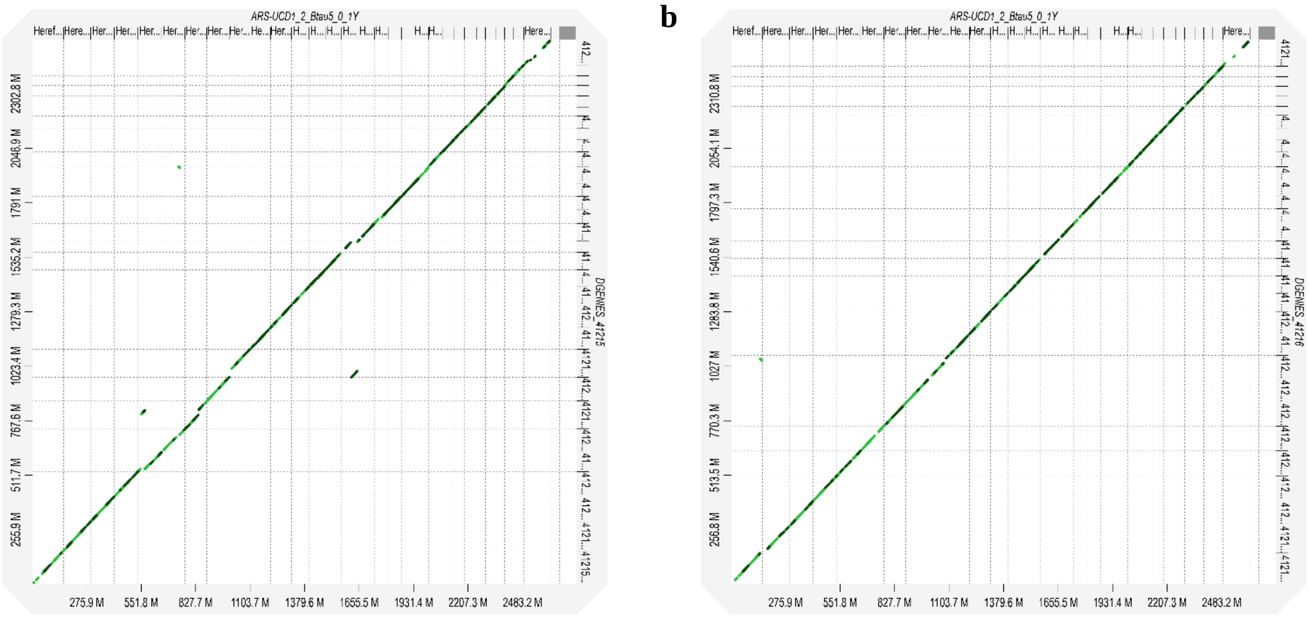

**Figure S11** Chromosomal alignment concordance between ARS-UCD1.2 on *x*-axis and Rouge Flamande assemblies on *y*-axis: **a** 41215, and **b** 41216

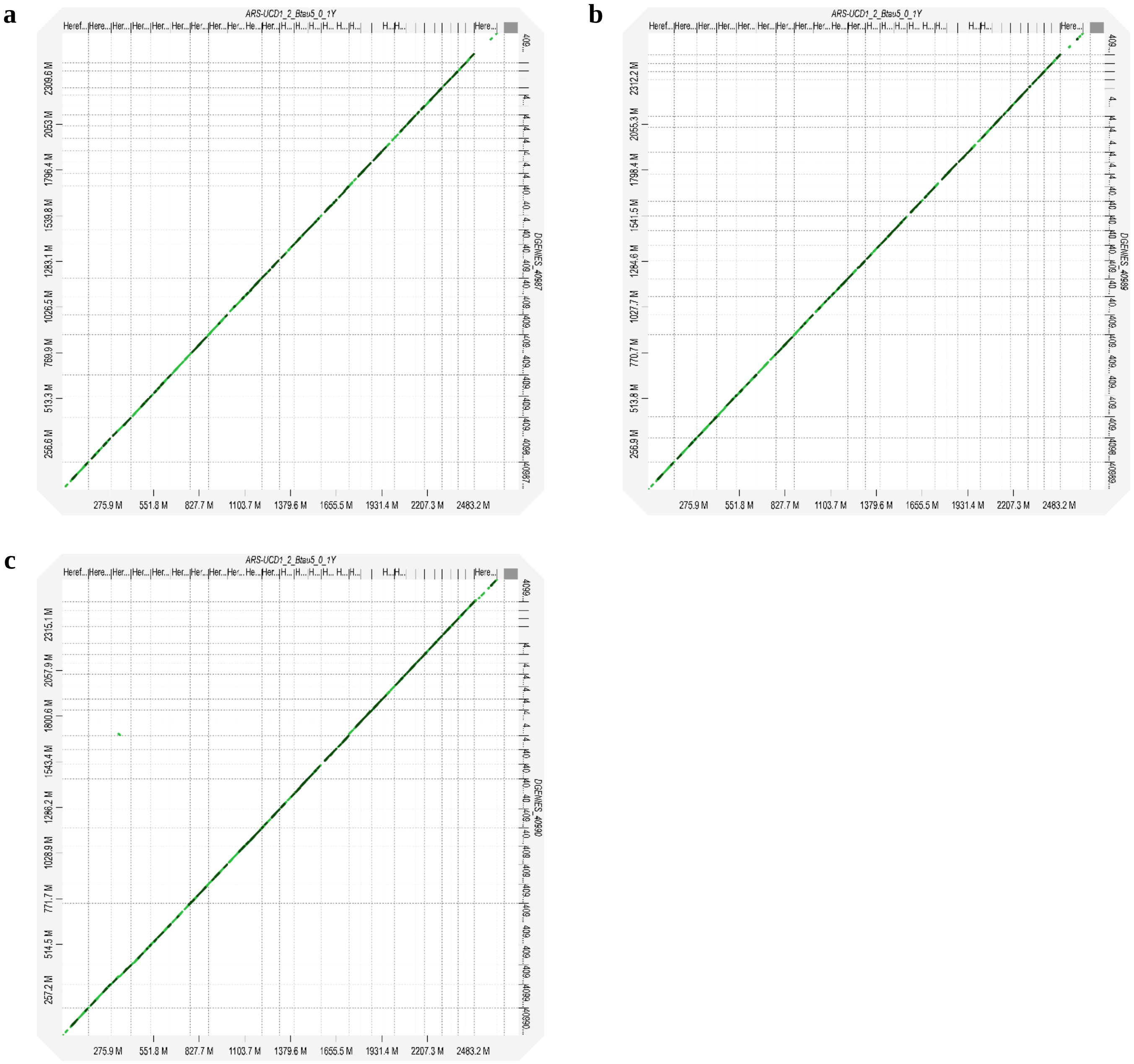


**Figure S12** Chromosomal alignment concordance between ARS-UCD1.2 on *x*-axis and Simmentale assemblies on *y*-axis: **a** 40987, **b** 40989, and **c** 40990

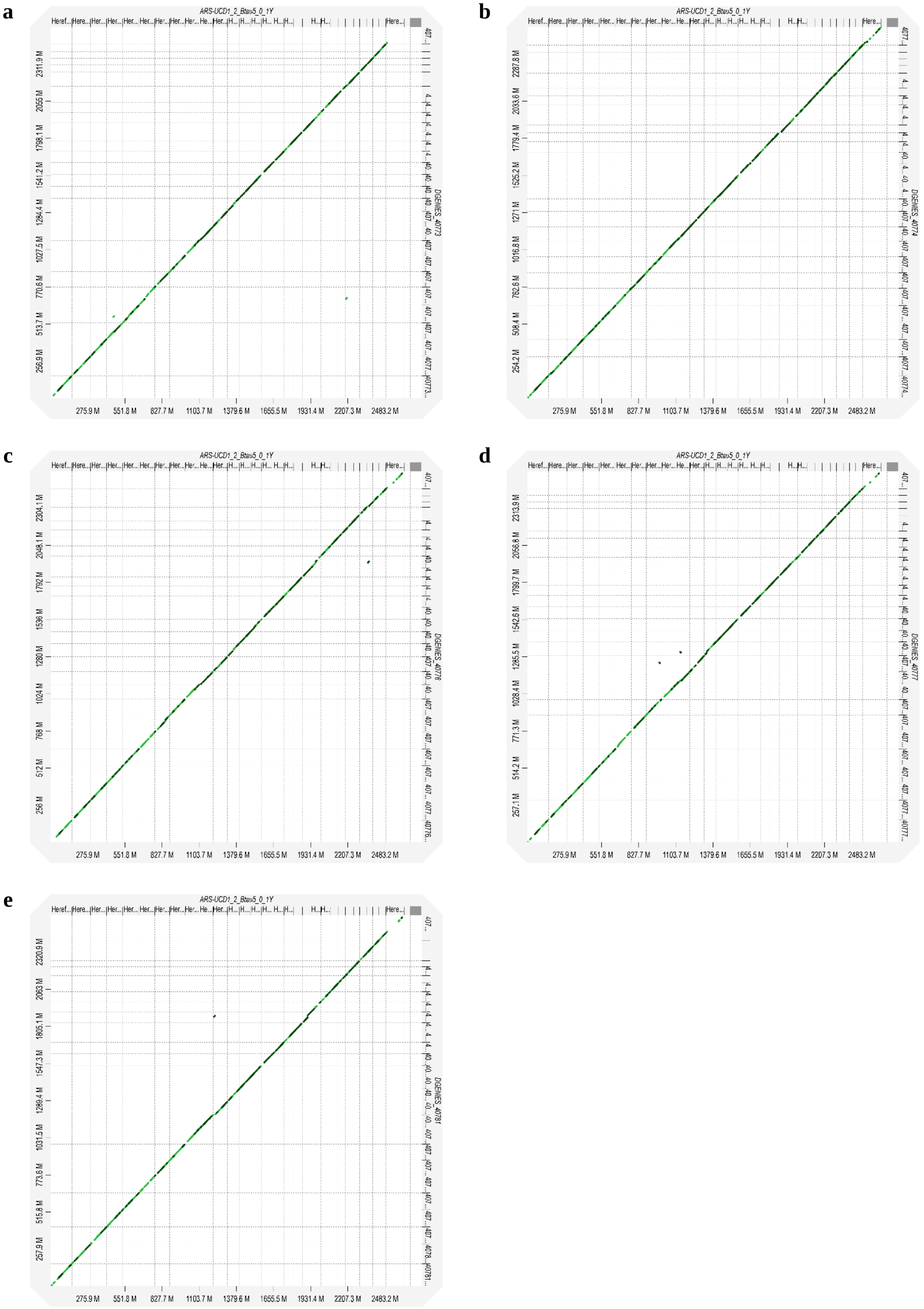


**Figure S13** Chromosomal alignment concordance between ARS-UCD1.2 on *x*-axis and Tarentaise assemblies on *y*-axis: **a** 40773, **b** 40774, **c** 40776, **d** 40777, and **e** 40781

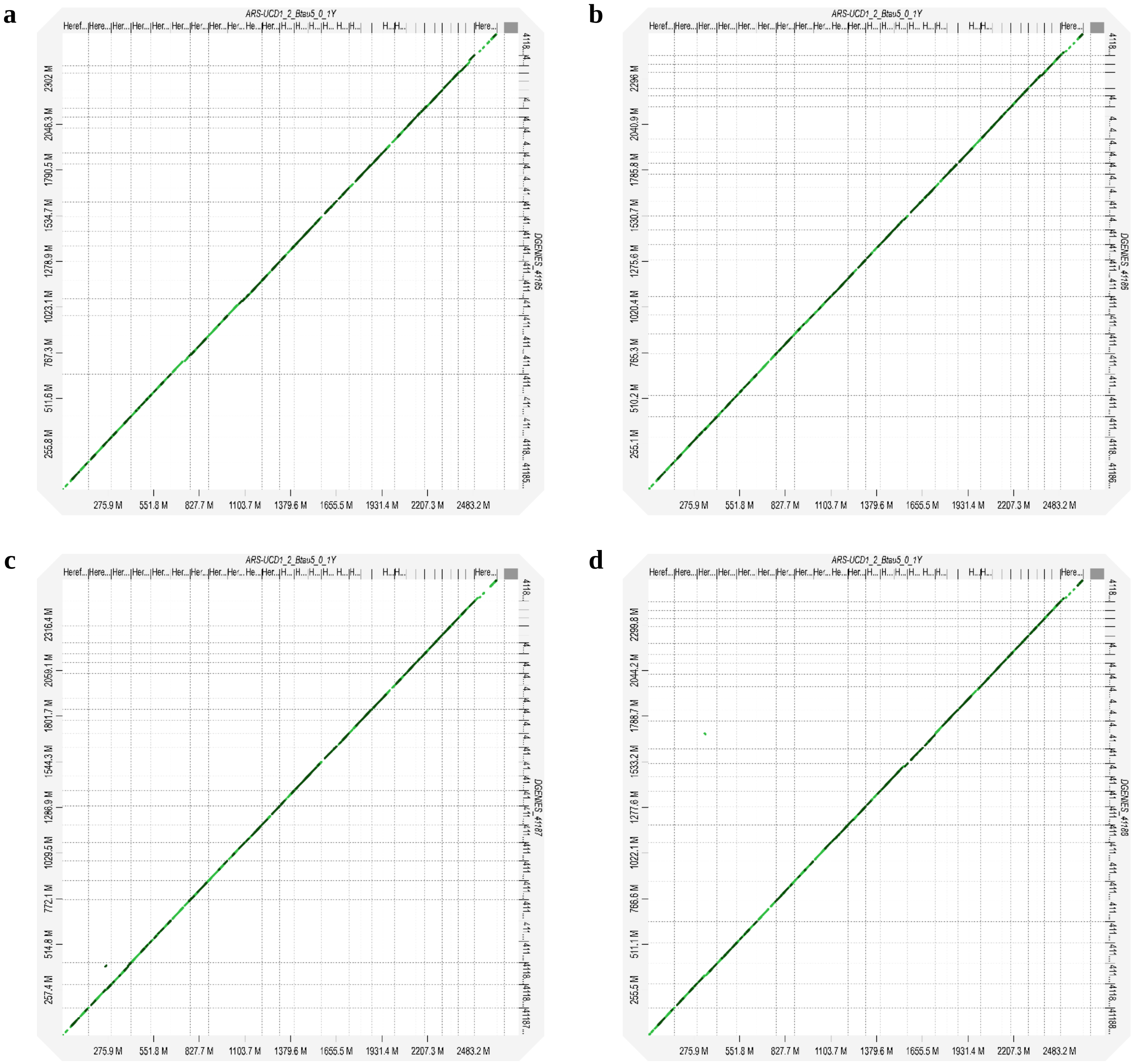


**Figure S14** Chromosomal alignment concordance between ARS-UCD1.2 on *x*-axis and Vosgienne assemblies on *y*-axis: **a** 41185, **b** 41186, **c** 41187, and **d** 41188
