## Supplementary material for "Application of a French cattle pangenome, from structural variant discovery to association studies on key phenotypes": Access_to_supplemental_files: Additional file 3 Figures S15-S28.pdf

**a**

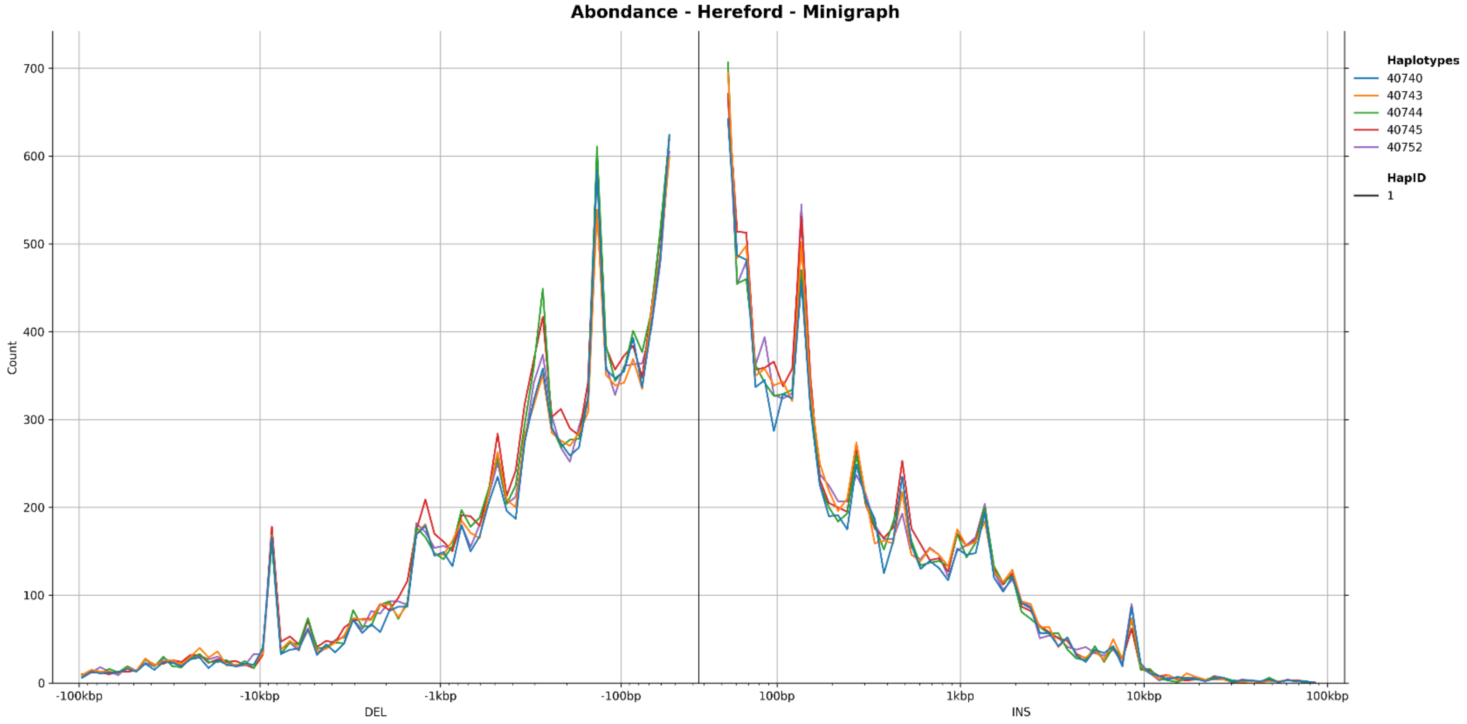


**b**

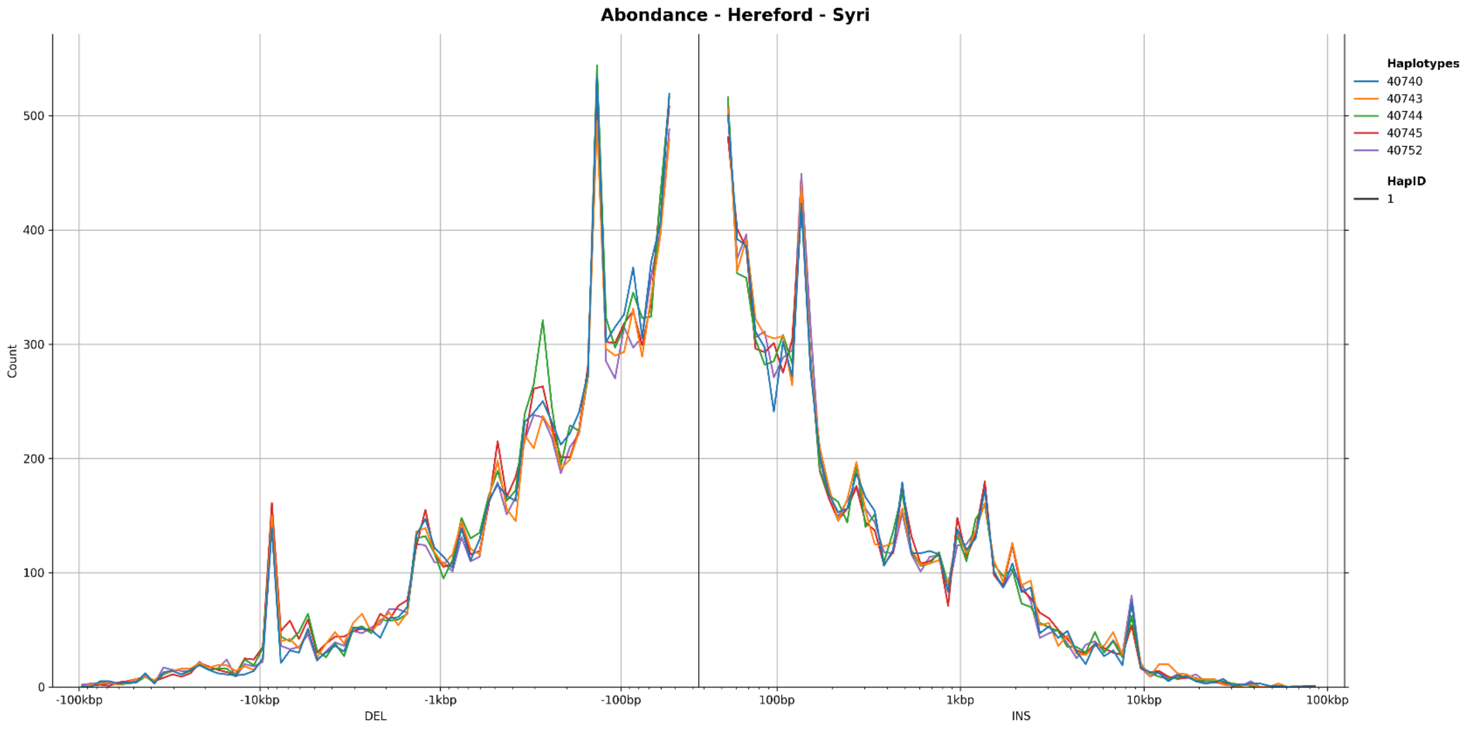


**Figure S15** Size distribution of SVs classified as deletions (left) and insertions (right), identified using **a** Minigraph, and **b** SyRI for the Abondance breed

**a**

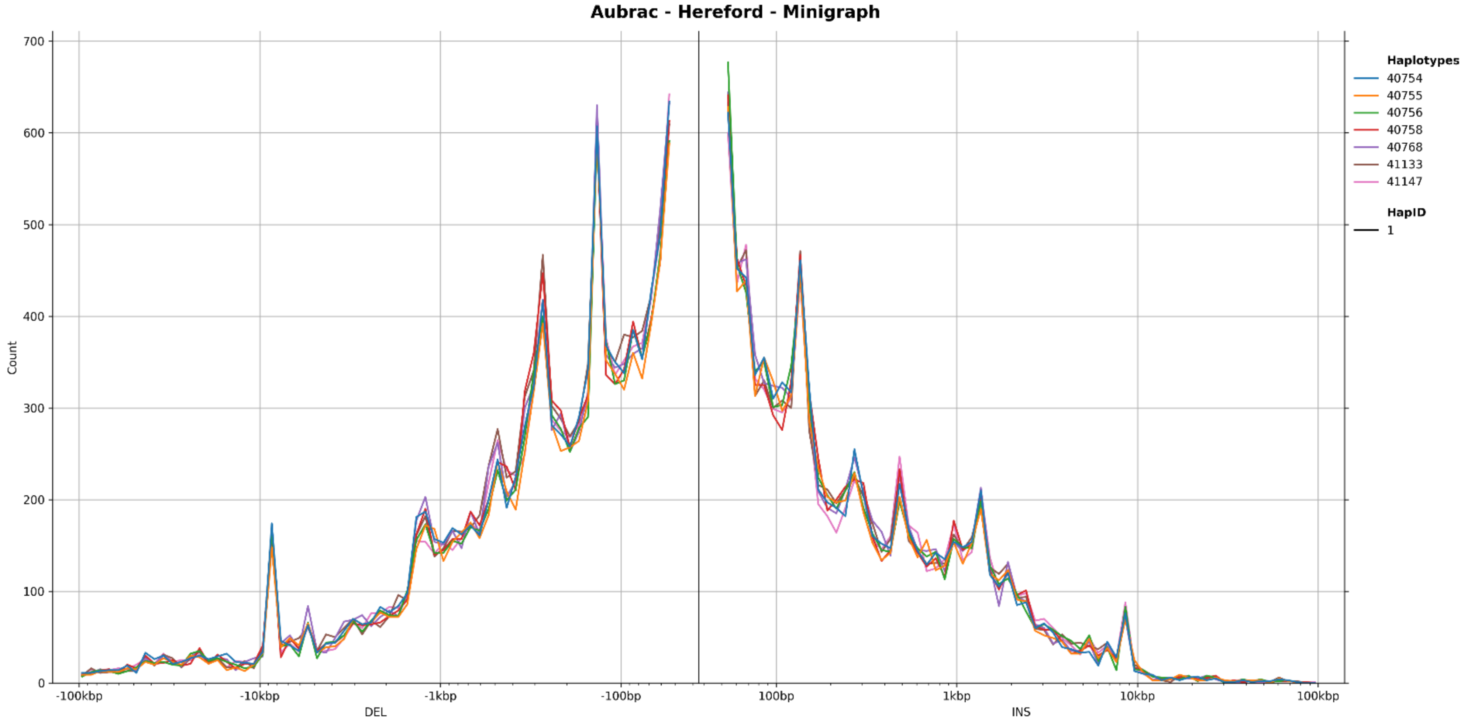


**b**

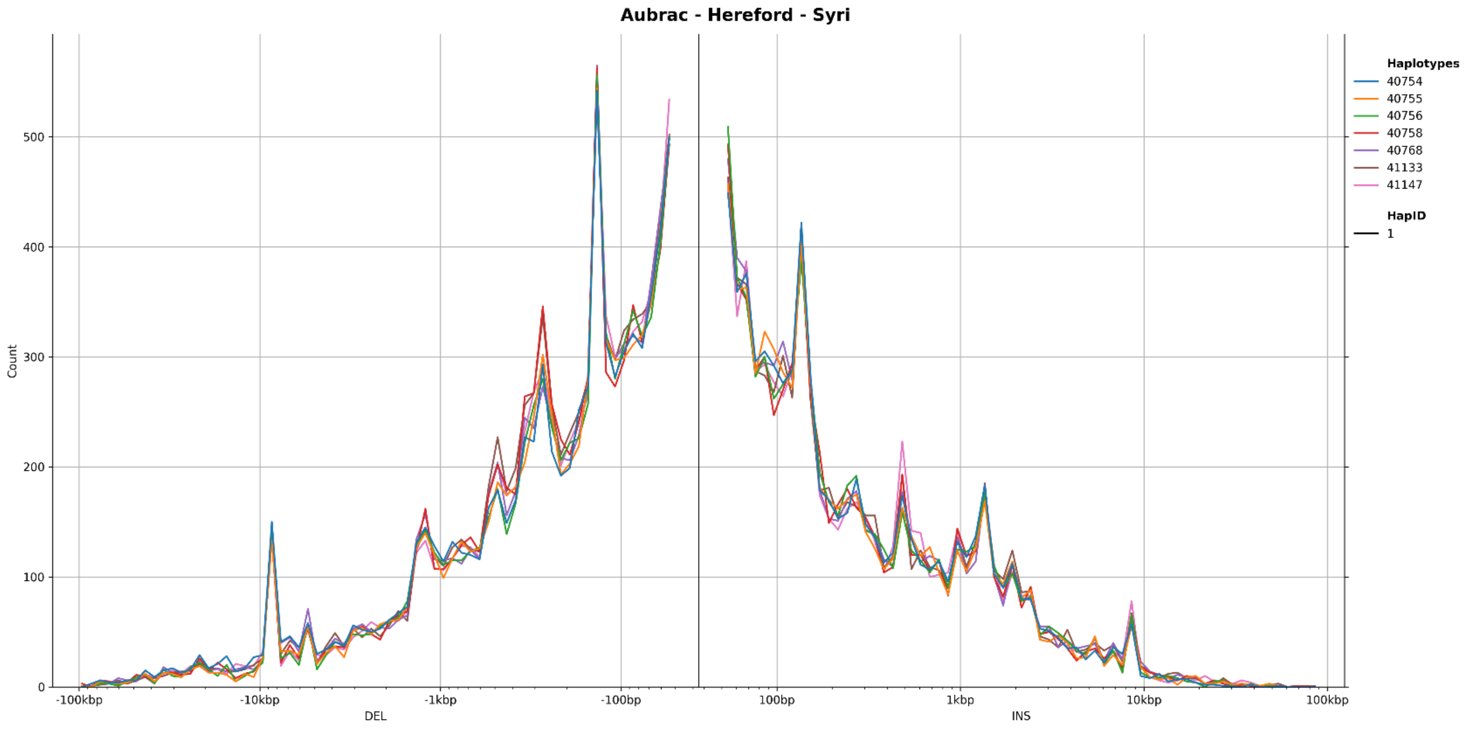


**Figure S16** Size distribution of SVs classified as deletions (left) and insertions (right), identified using **a** Minigraph, and **b** SyRI for the Aubrac breed

**a**

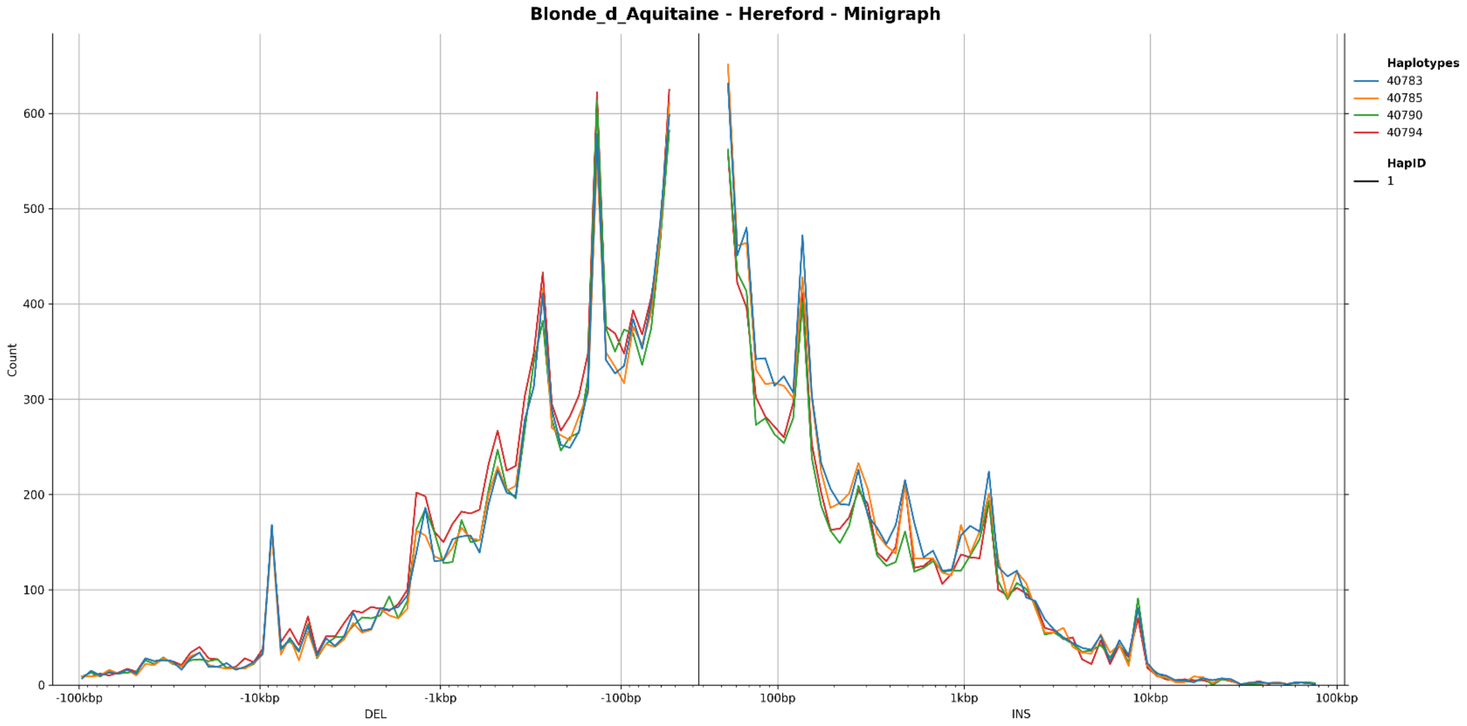


**b**

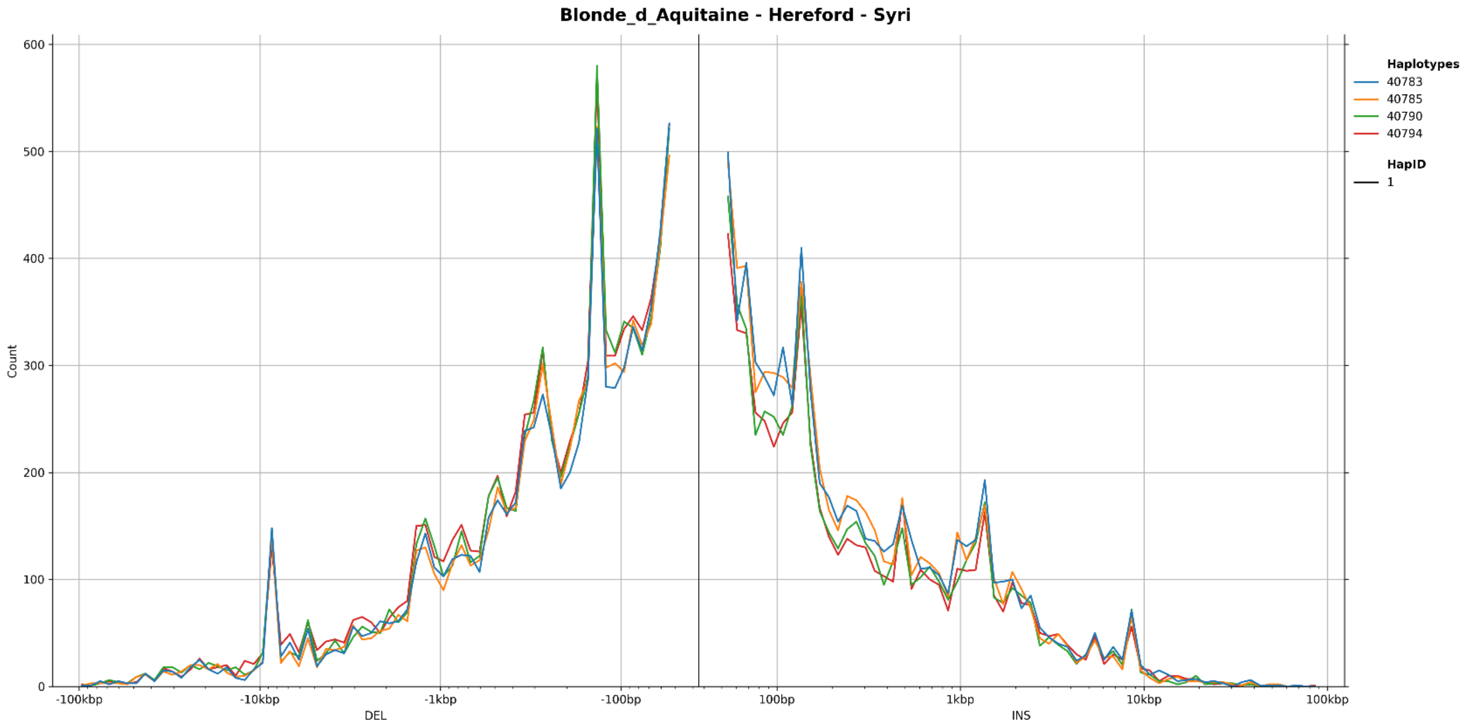


**Figure S17** Size distribution of SVs classified as deletions (left) and insertions (right), identified using **a** Minigraph, and **b** SyRI for the Blonde d'Aquitaine breed

**a**

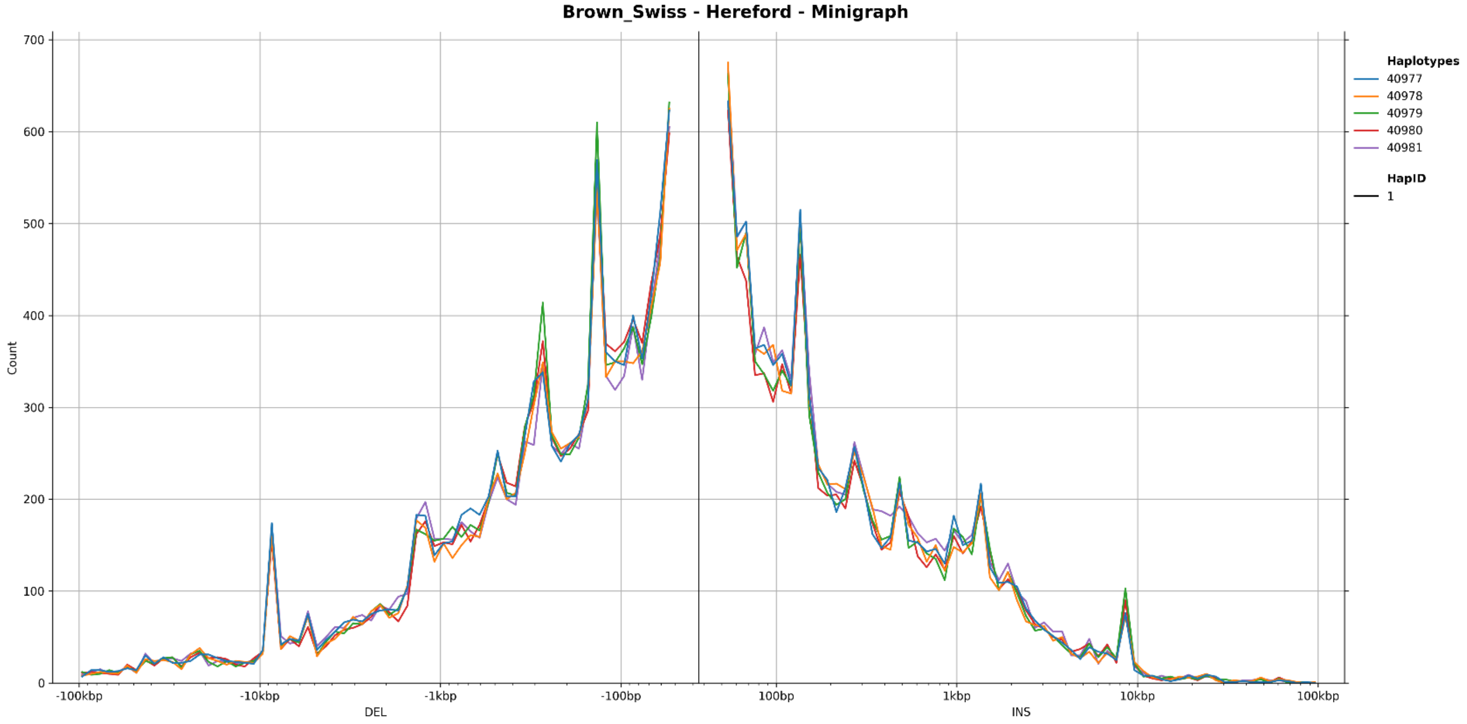


**b**

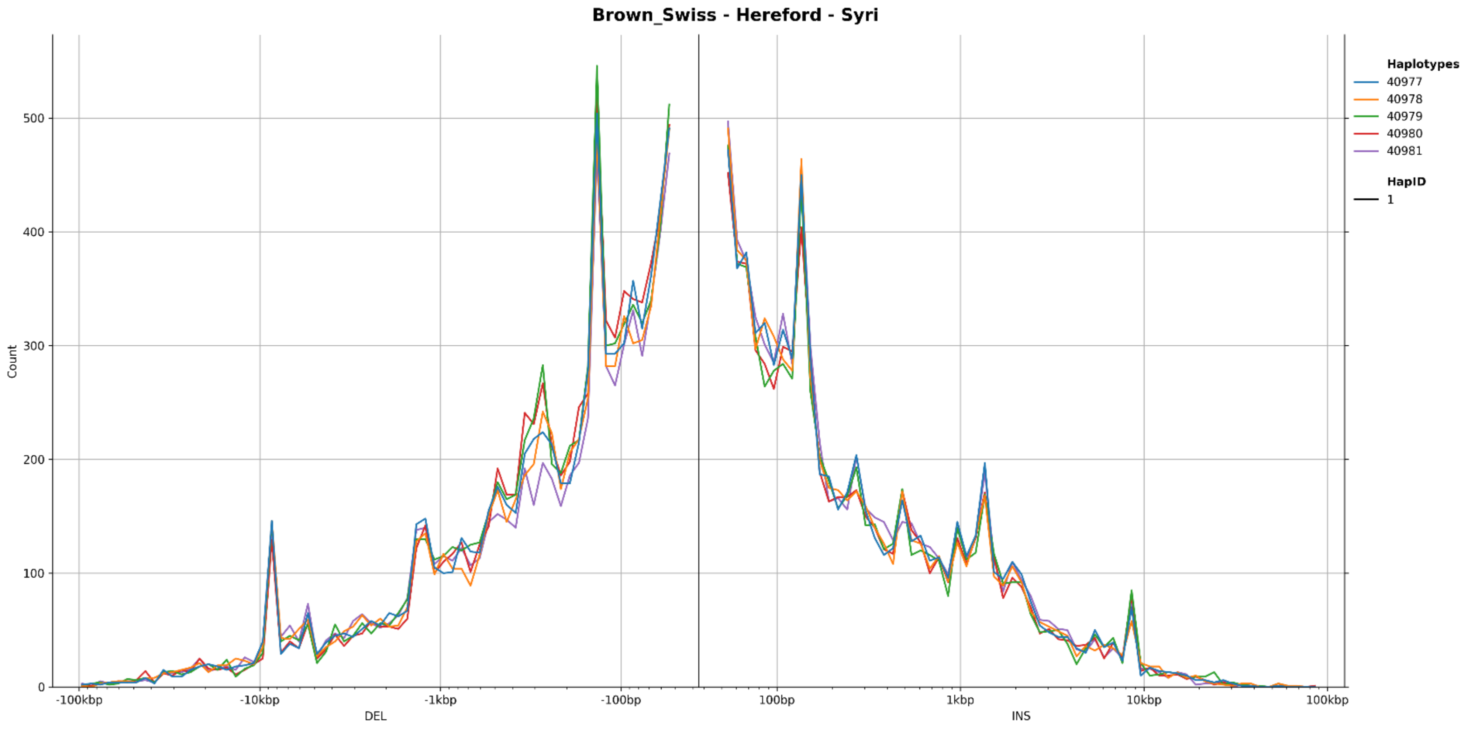


**Figure S18** Size distribution of SVs classified as deletions (left) and insertions (right), identified using **a** Minigraph, and **b** SyRI for the Brown Swiss breed

**a**

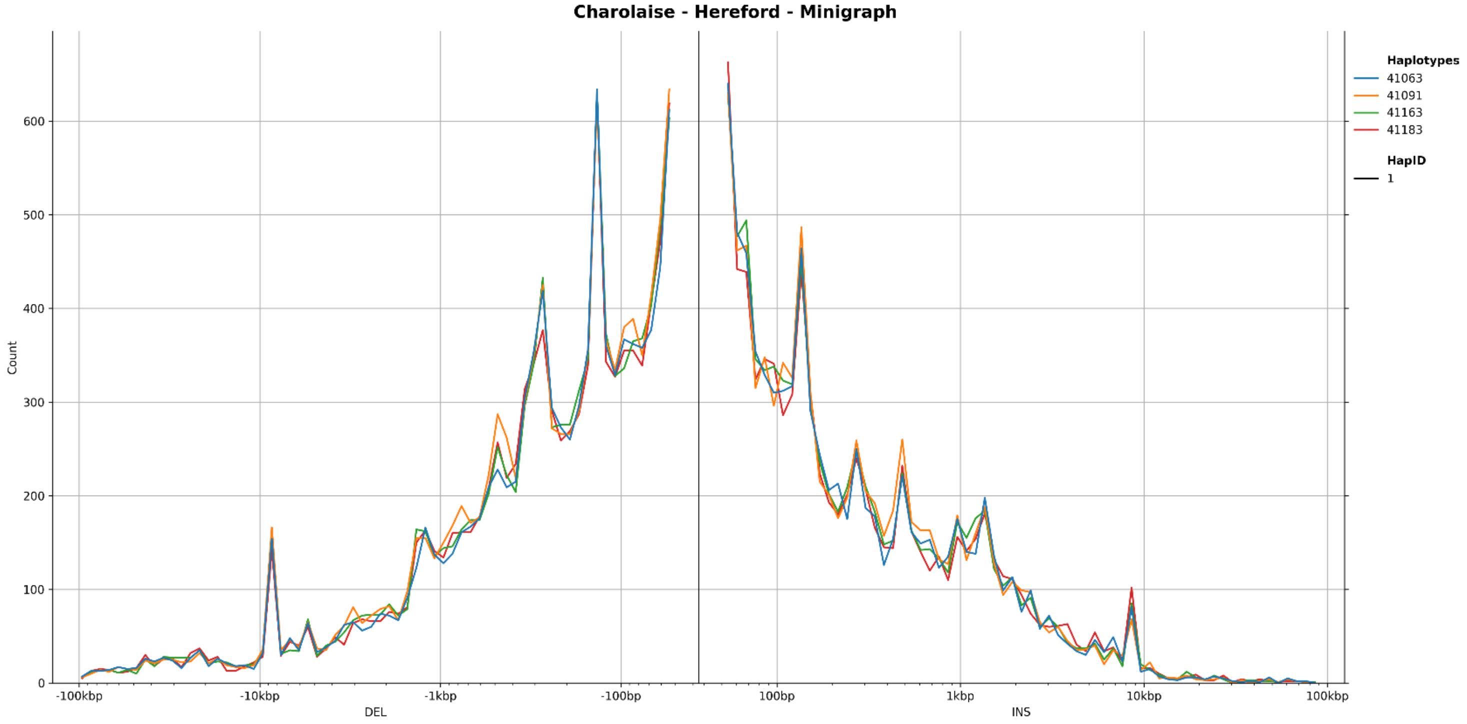


**b**

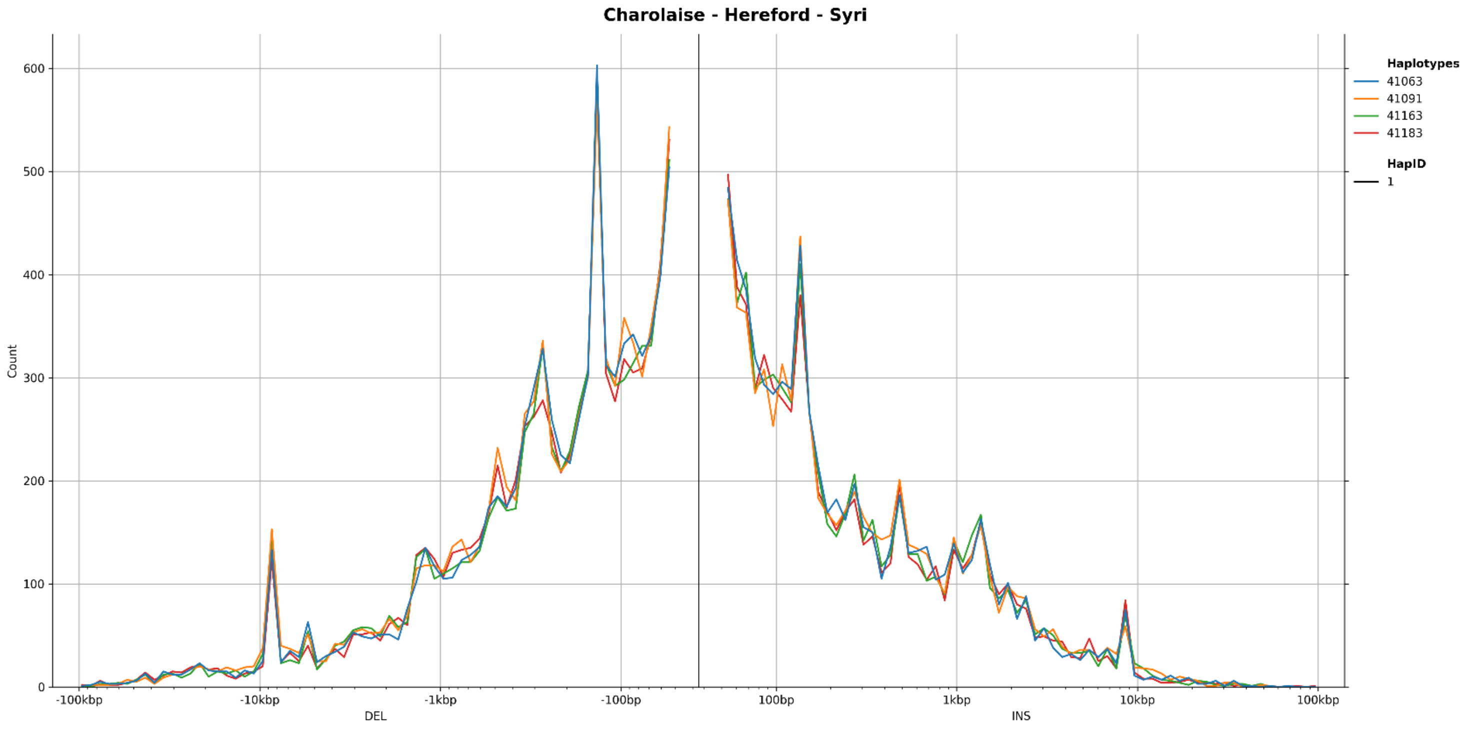


**Figure S19** Size distribution of SVs classified as deletions (left) and insertions (right), identified using **a** Minigraph, and **b** SyRI for the Charolaise breed

a

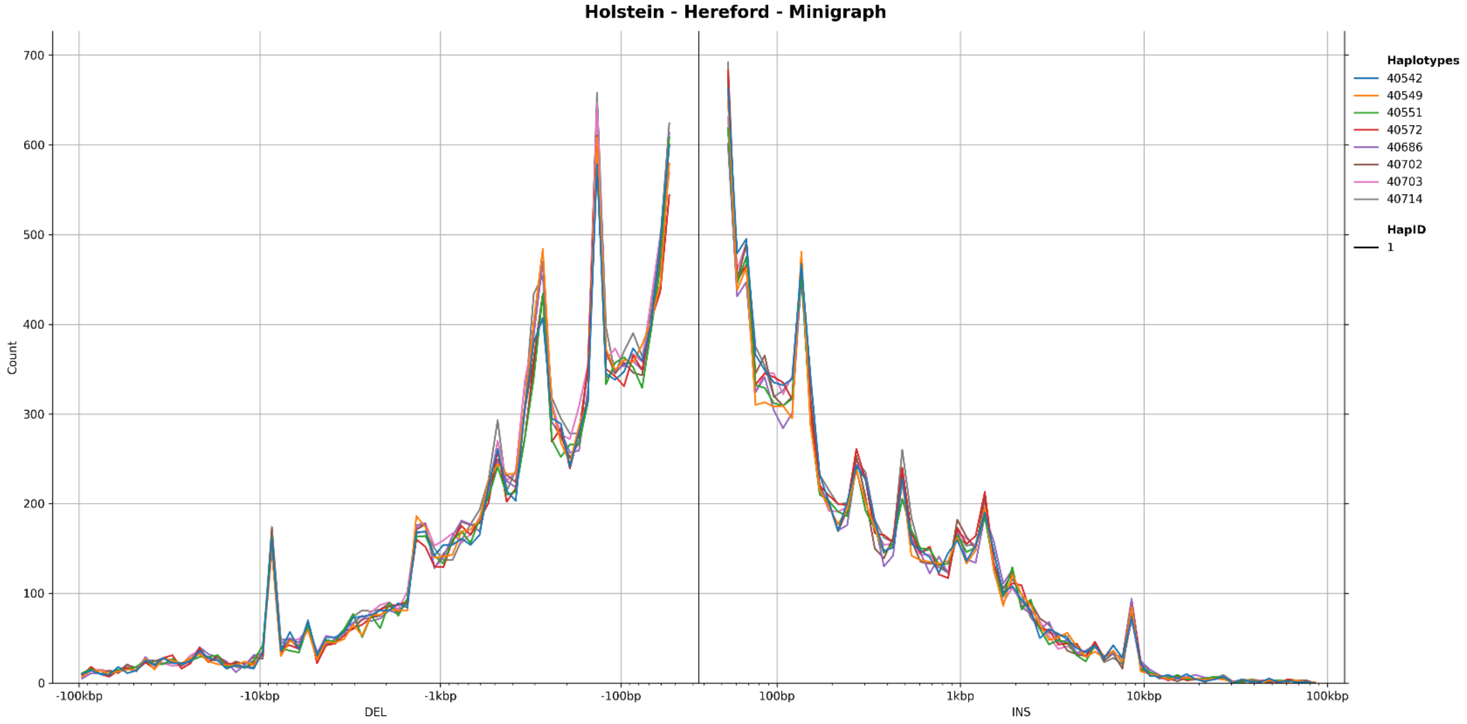


b

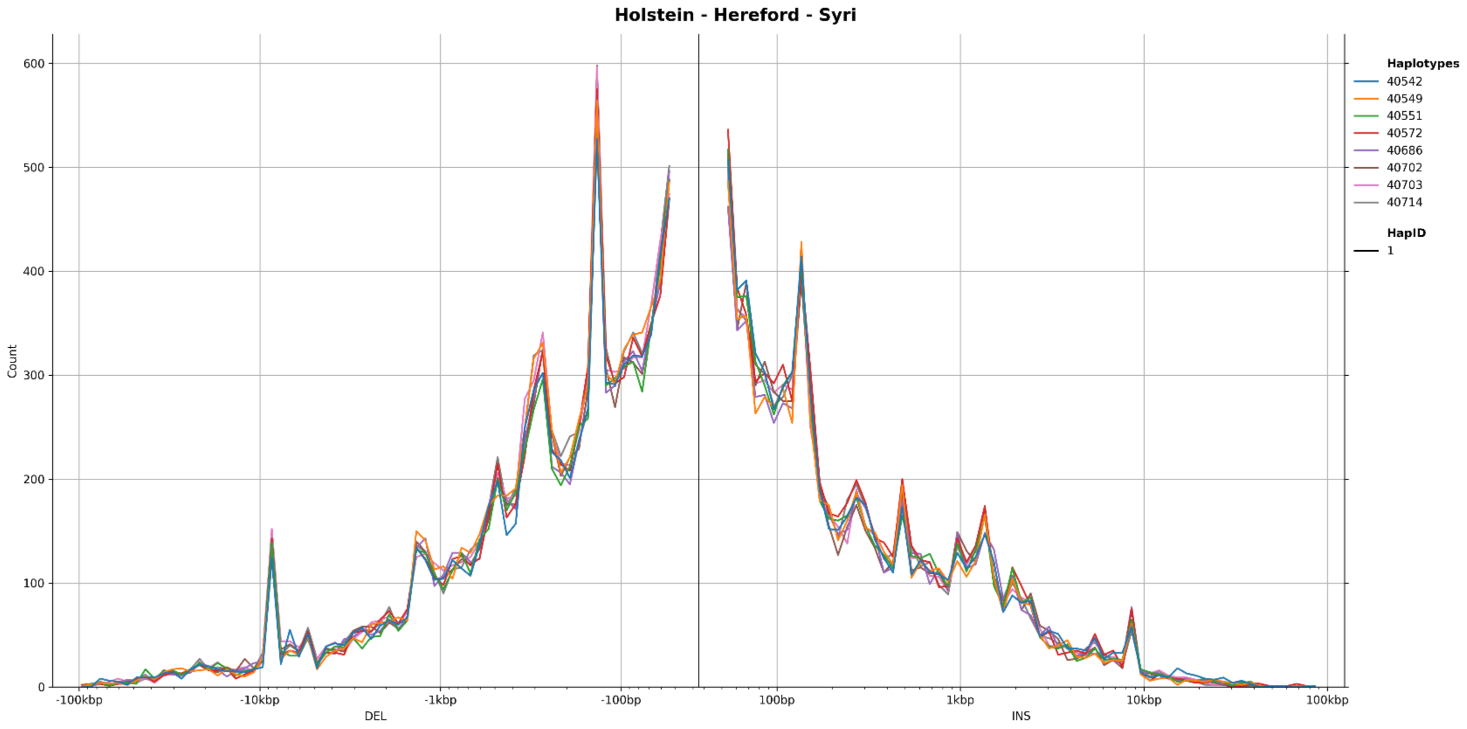


**Figure S20** Size distribution of SVs classified as deletions (left) and insertions (right), identified using **a** Minigraph, and **b** SyRI for the Holstein breed

**a**

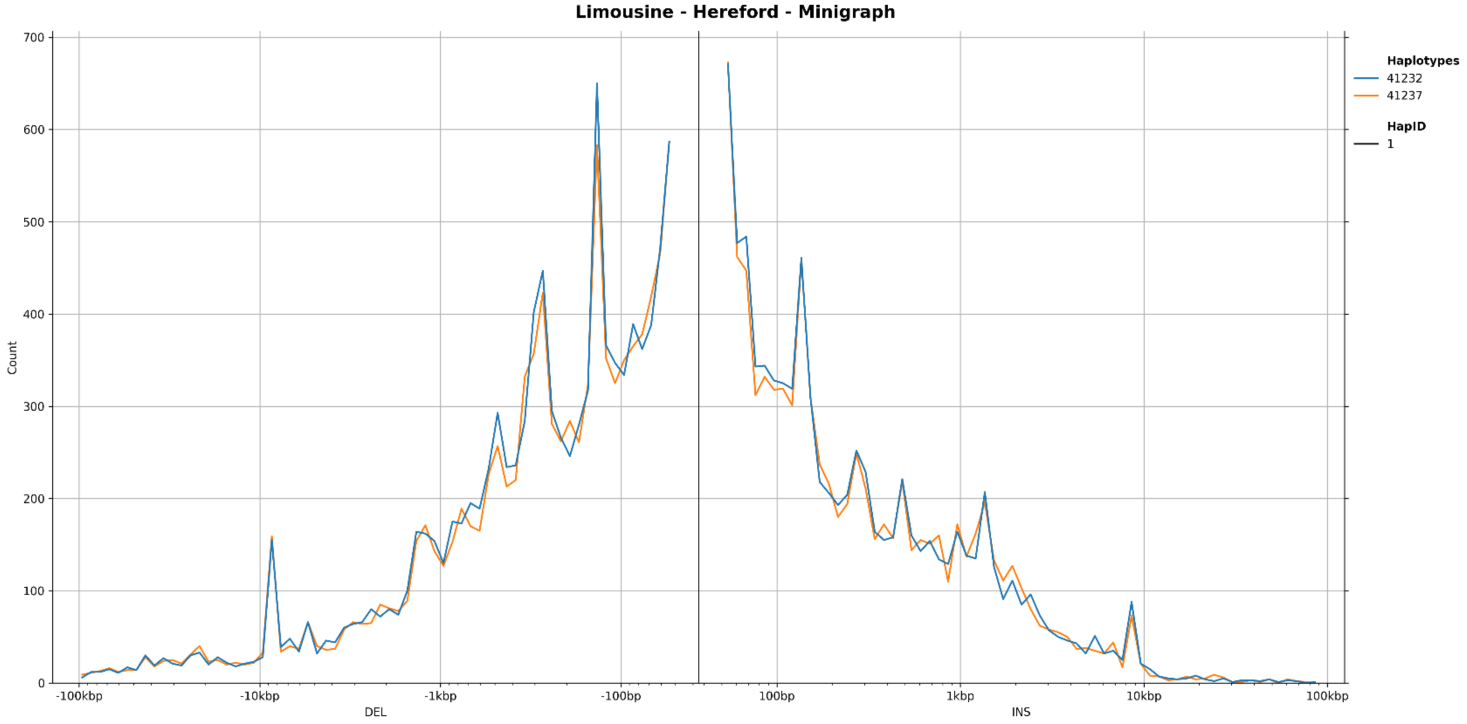


**b**

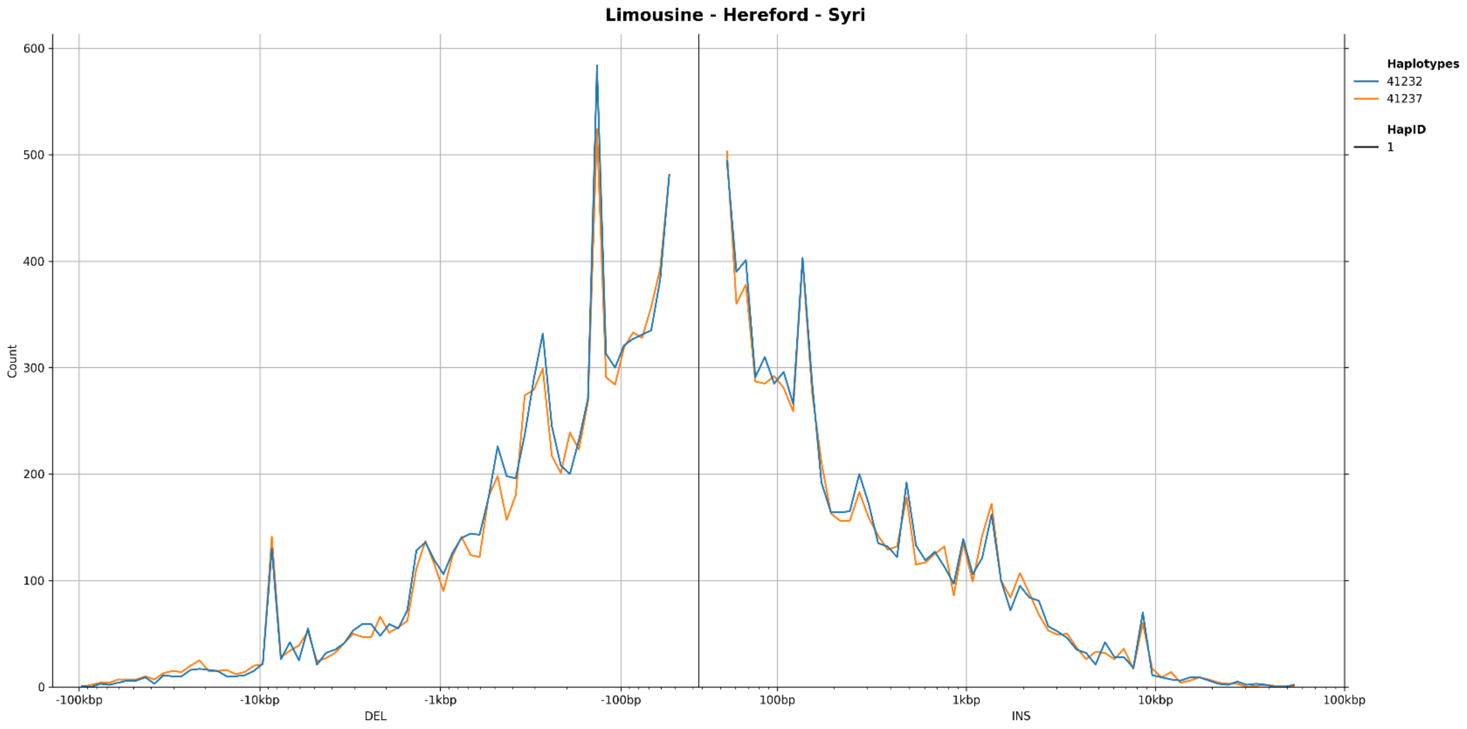


**Figure S21** Size distribution of SVs classified as deletions (left) and insertions (right), identified using **a** Minigraph, and **b** SyRI for the Limousine breed

**a**

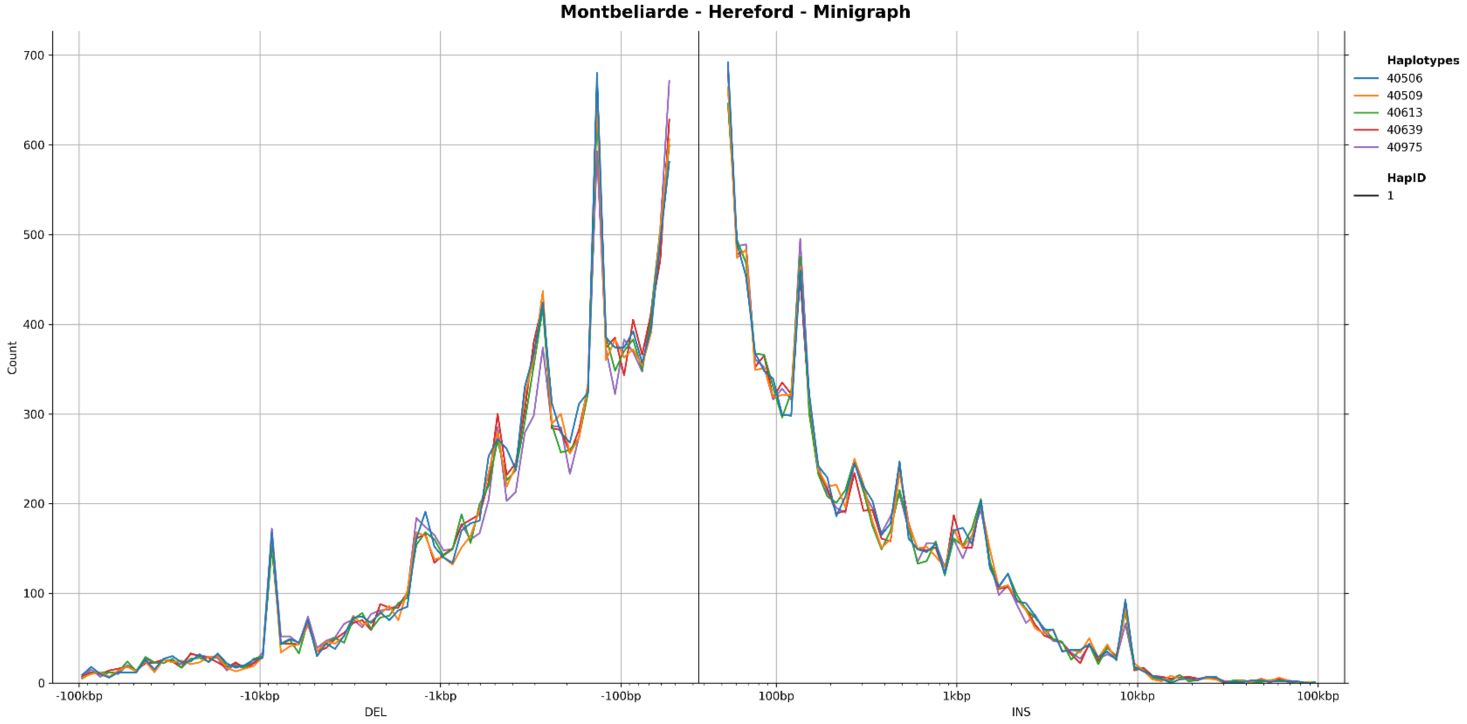


**b**

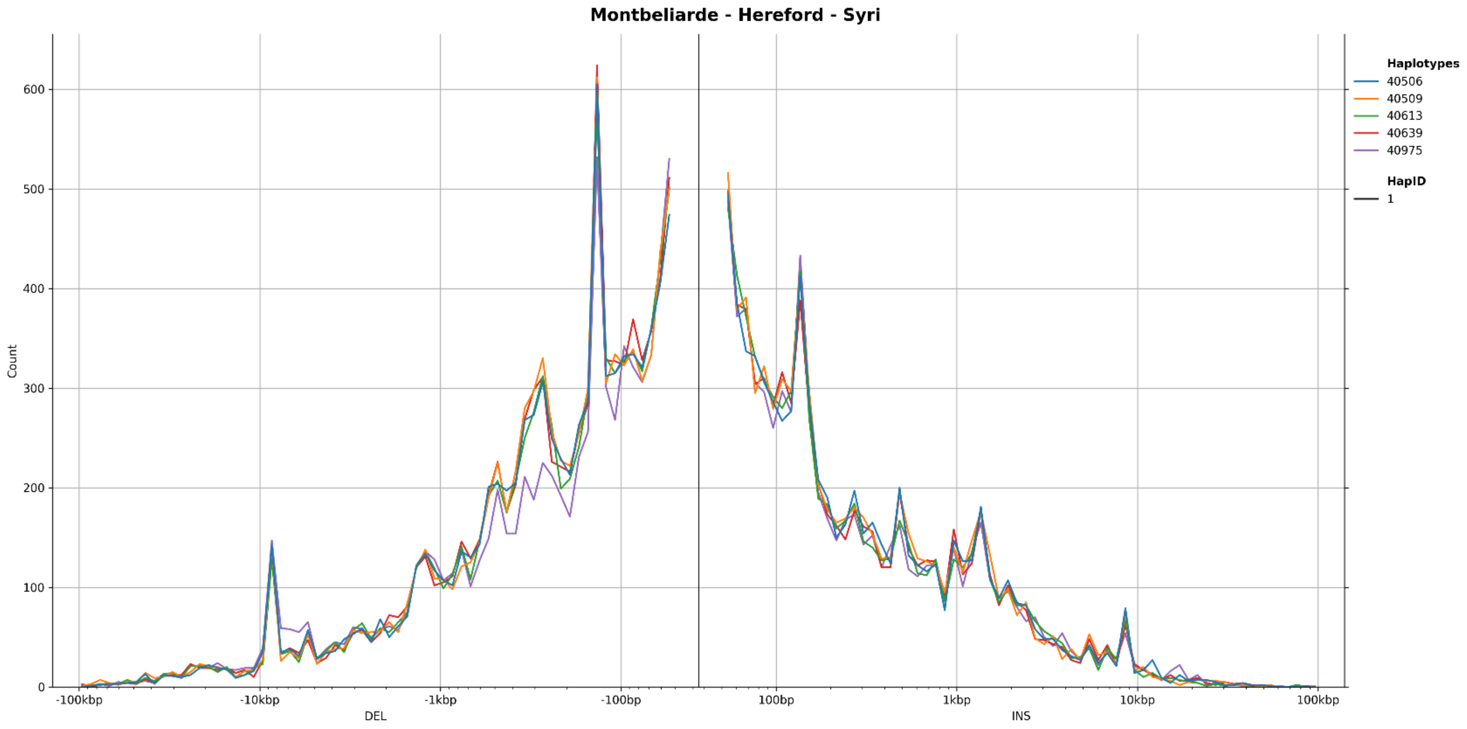


**Figure S22** Size distribution of SVs classified as deletions (left) and insertions (right), identified using **a** Minigraph, and **b** SyRI for the Montbéliarde breed

**a**

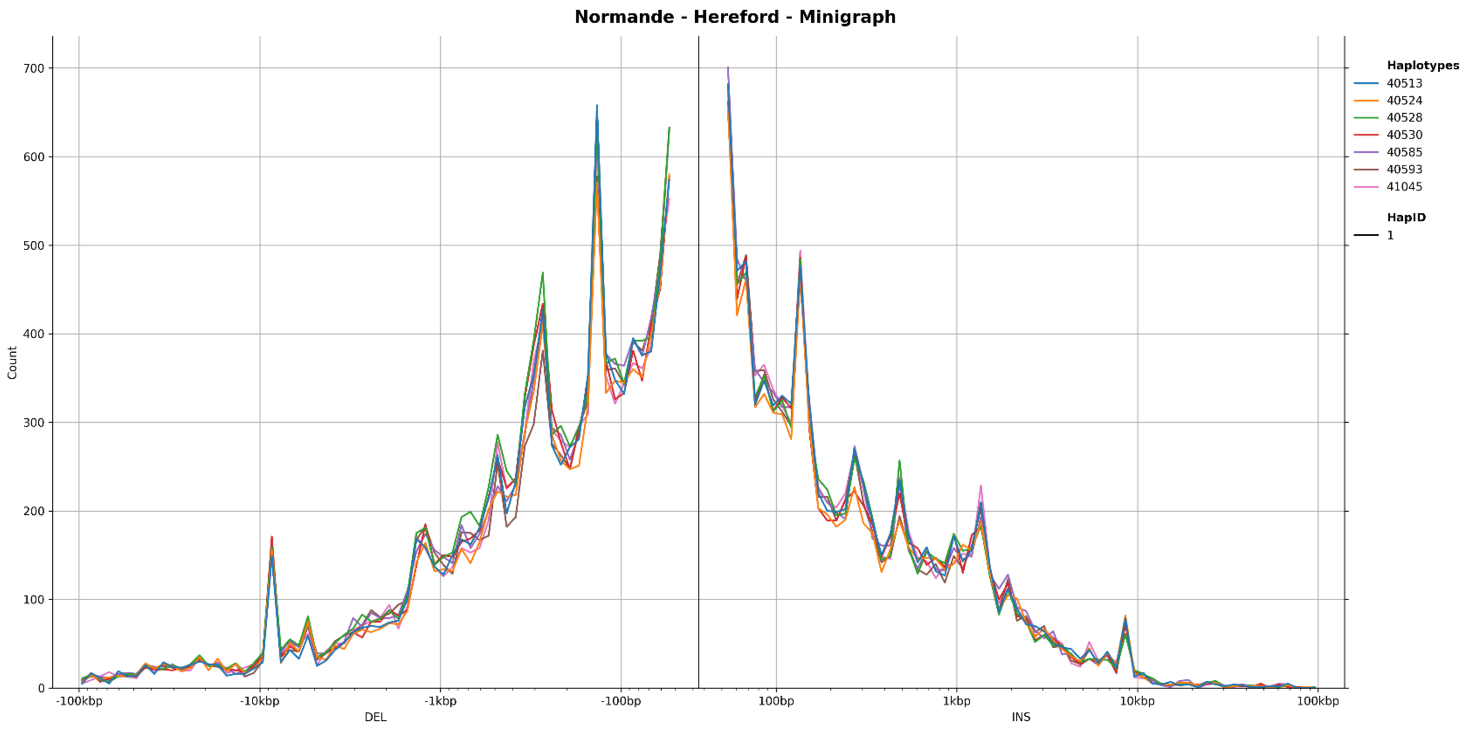


**b**

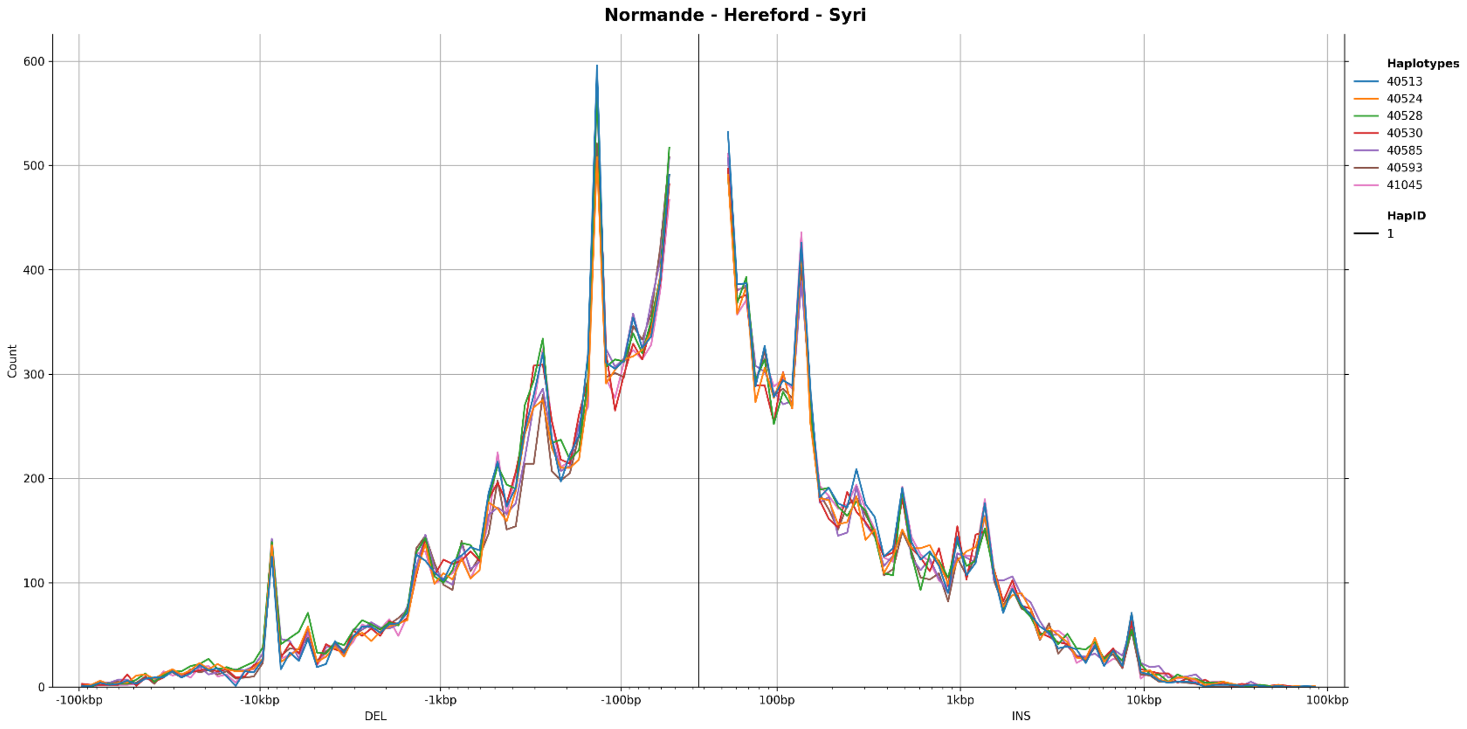


**Figure S23** Size distribution of SVs classified as deletions (left) and insertions (right), identified using **a** Minigraph, and **b** SyRI for the Normande breed

**a**

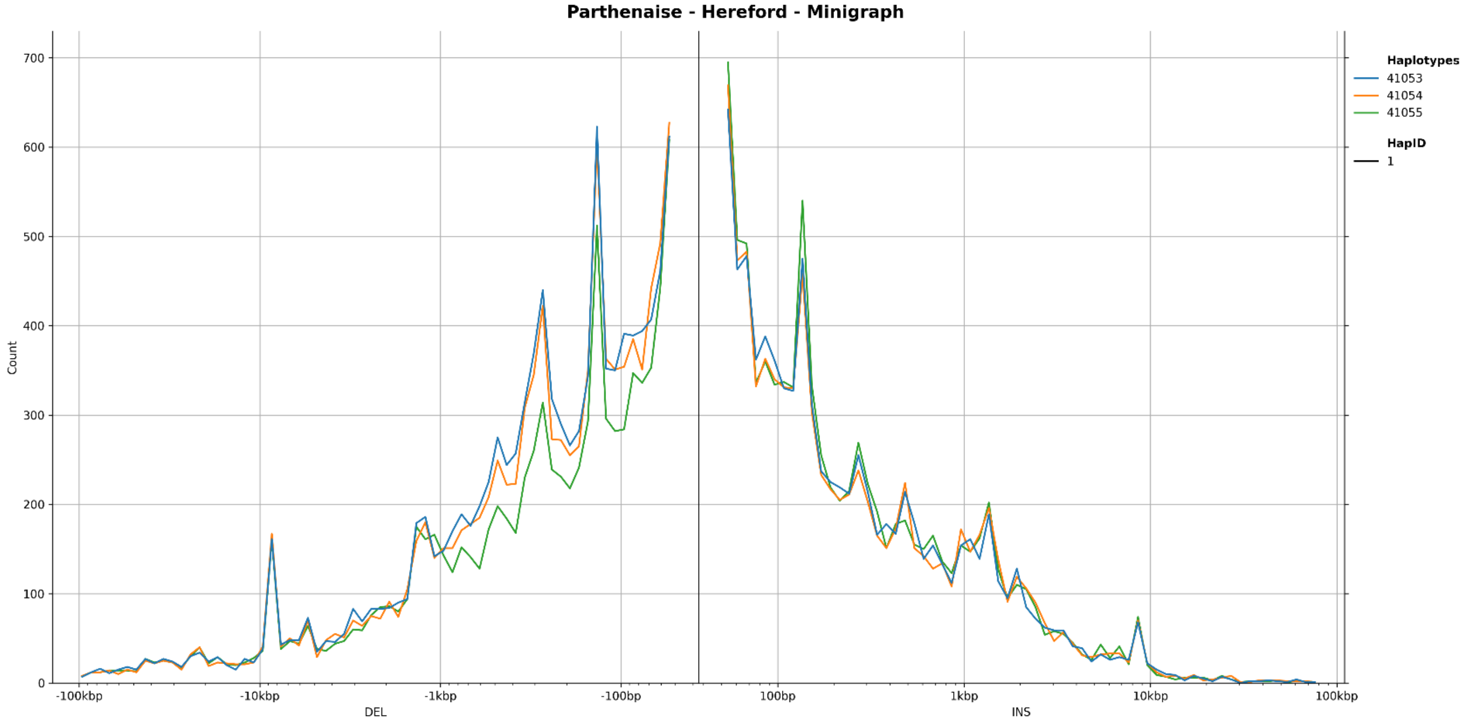


**b**

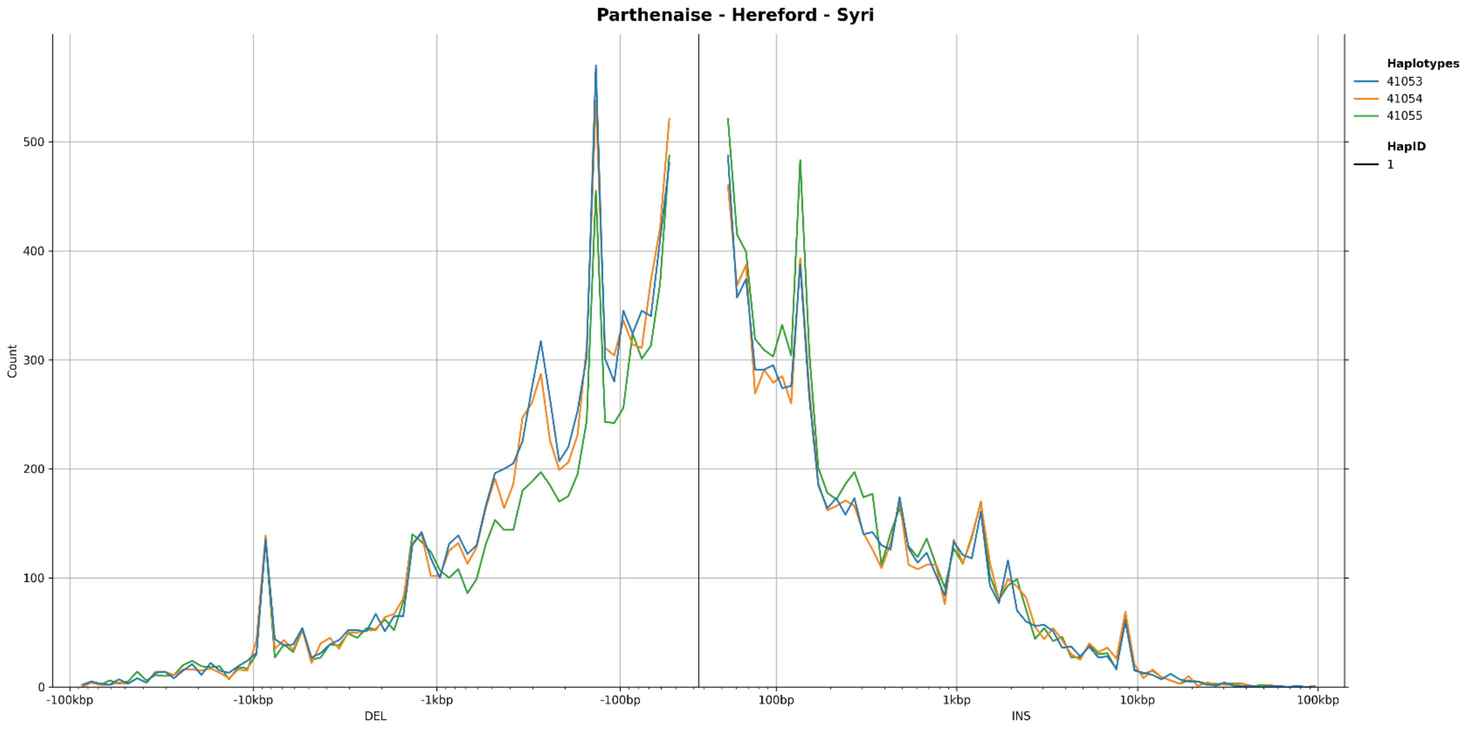


**Figure S24** Size distribution of SVs classified as deletions (left) and insertions (right), identified using **a** Minigraph, and **b** SyRI for the Parthenaise breed

**a**

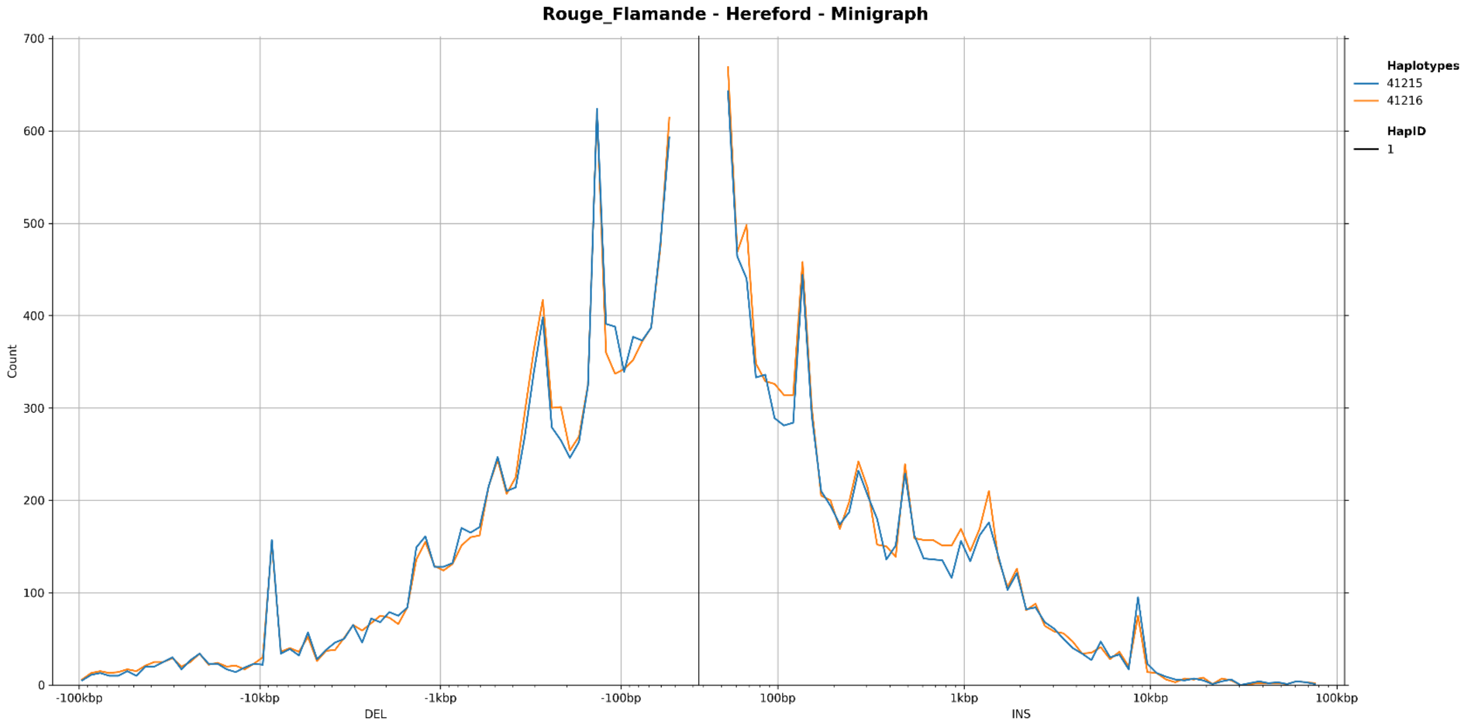


**b**

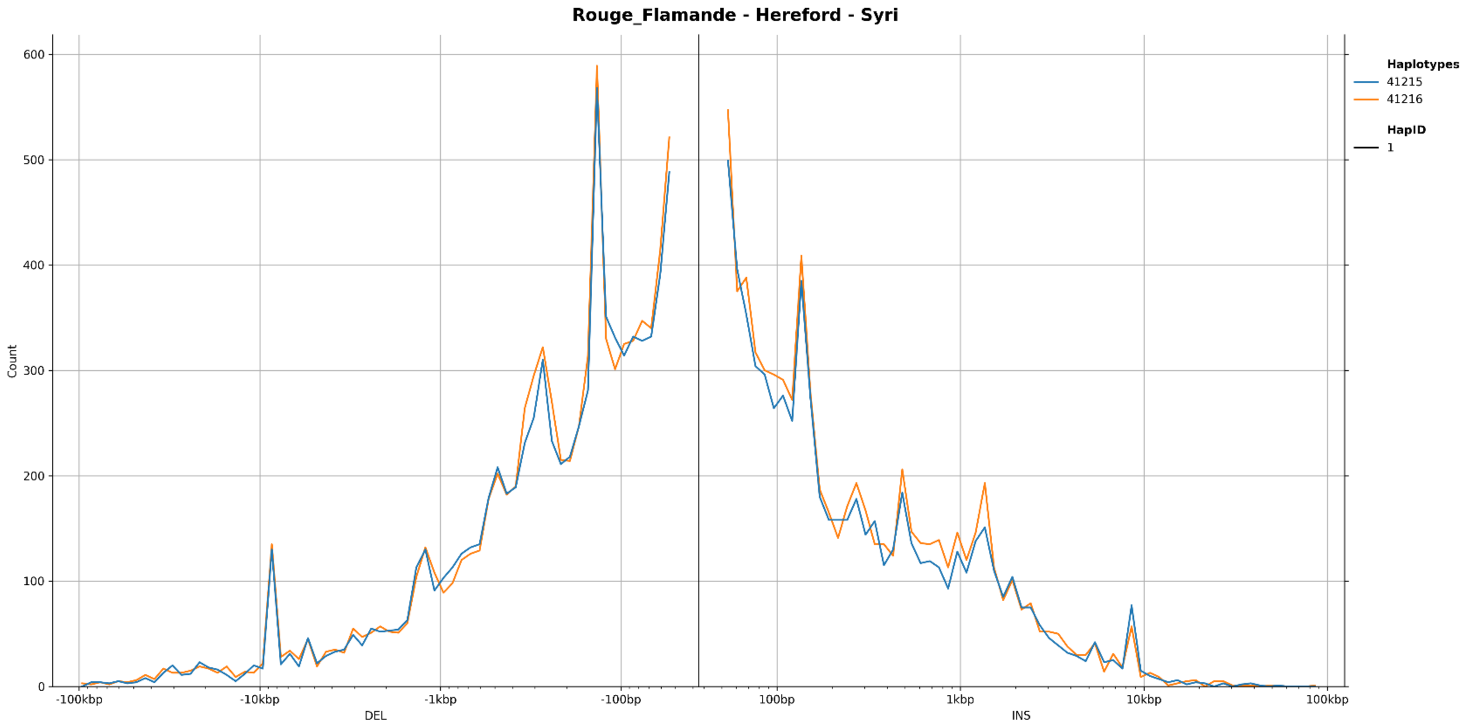


**Figure S25** Size distribution of SVs classified as deletions (left) and insertions (right), identified using **a** Minigraph, and **b** SyRI for the Rouge Flamande breed

a

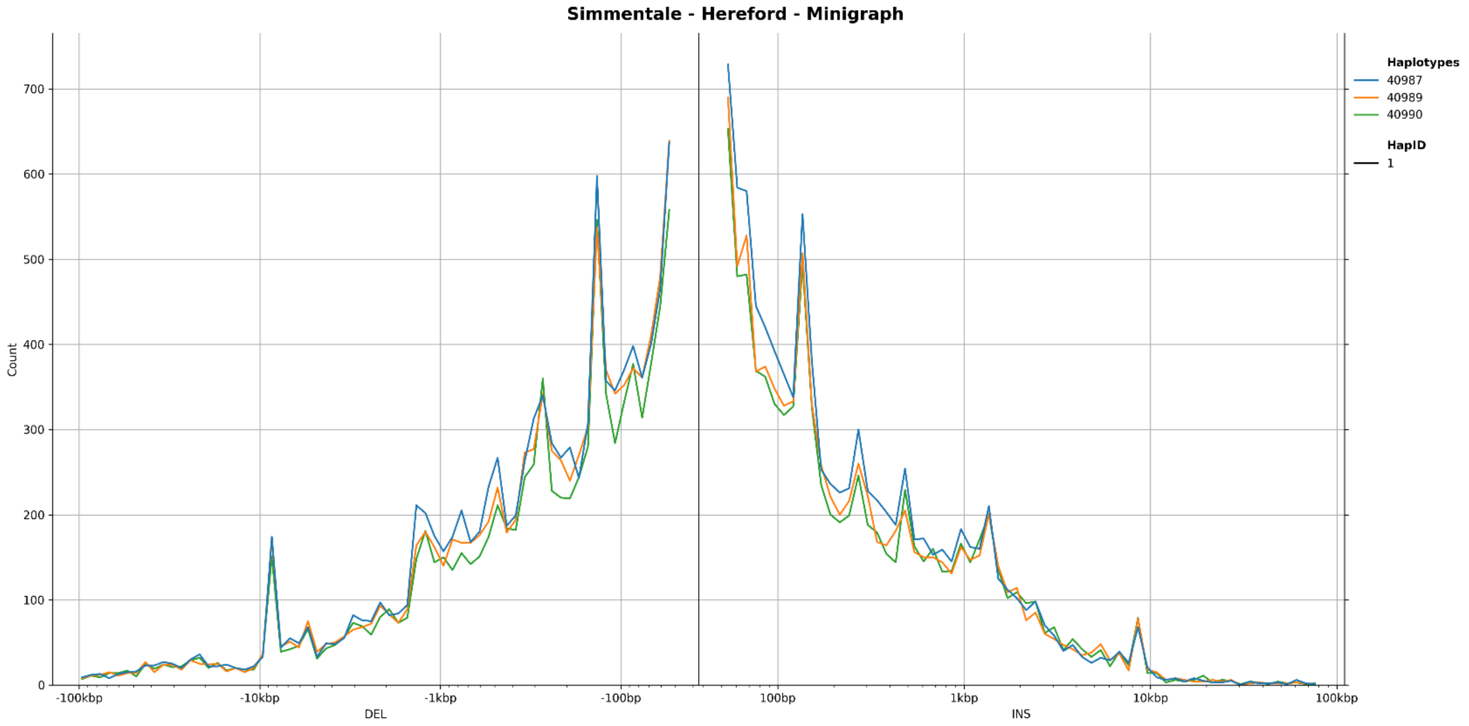


b

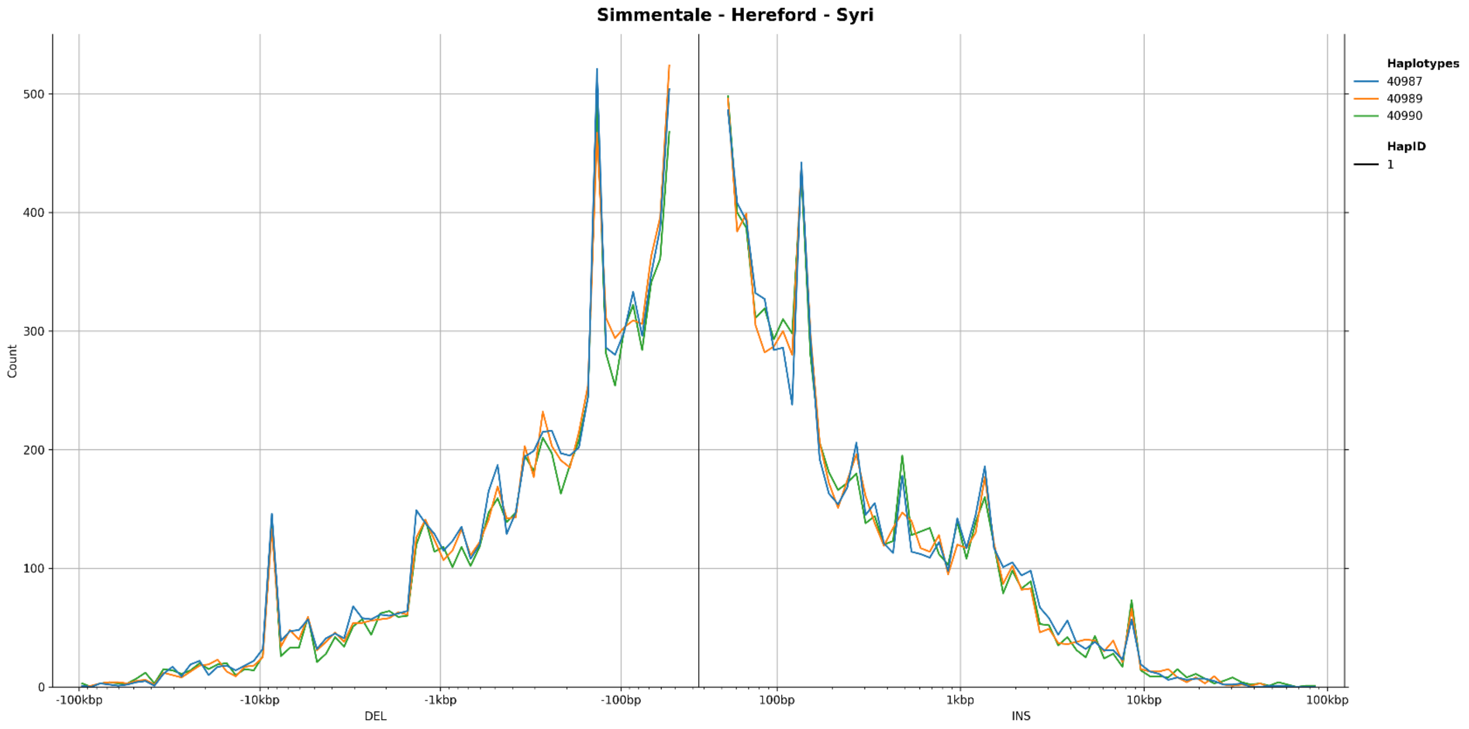


**Figure S26** Size distribution of SVs classified as deletions (left) and insertions (right), identified using **a** Minigraph, and **b** SyRI for the Simmentale breed

**a**

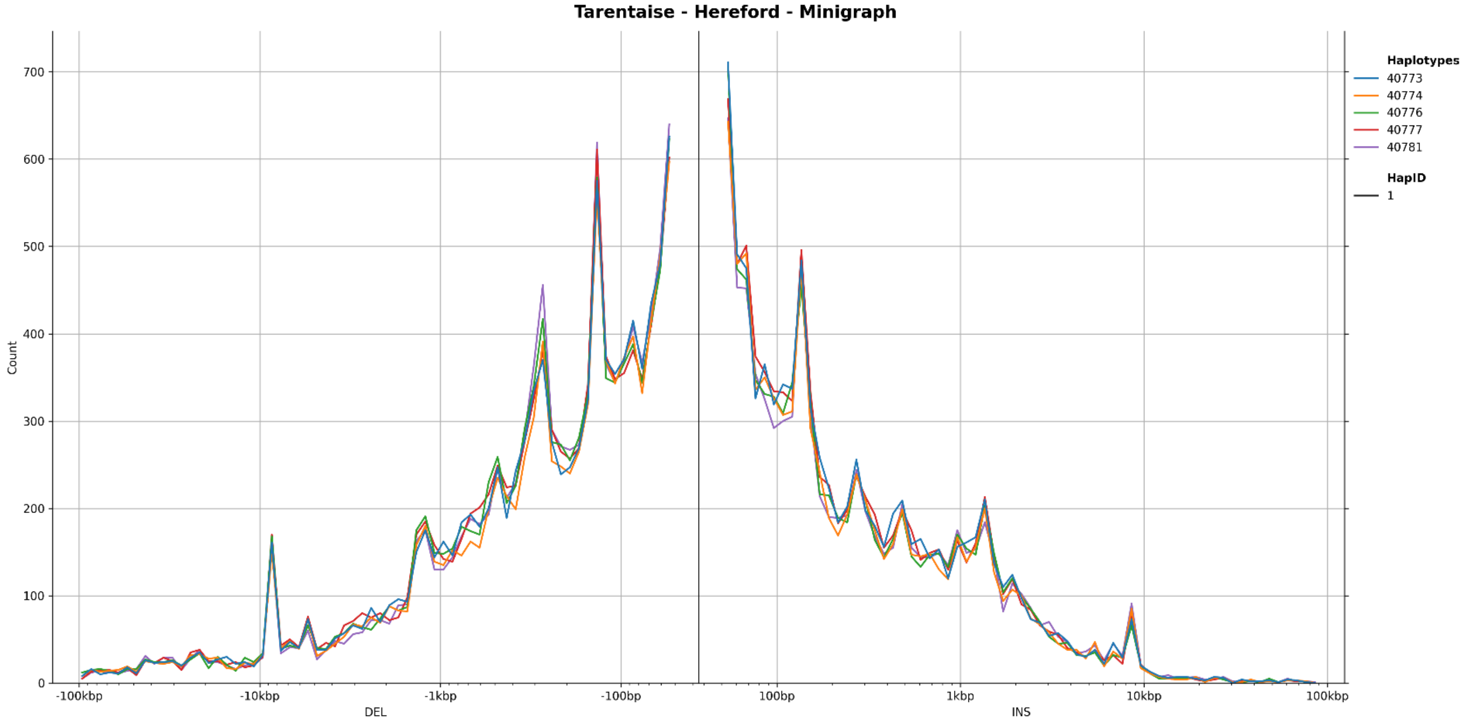


**b**

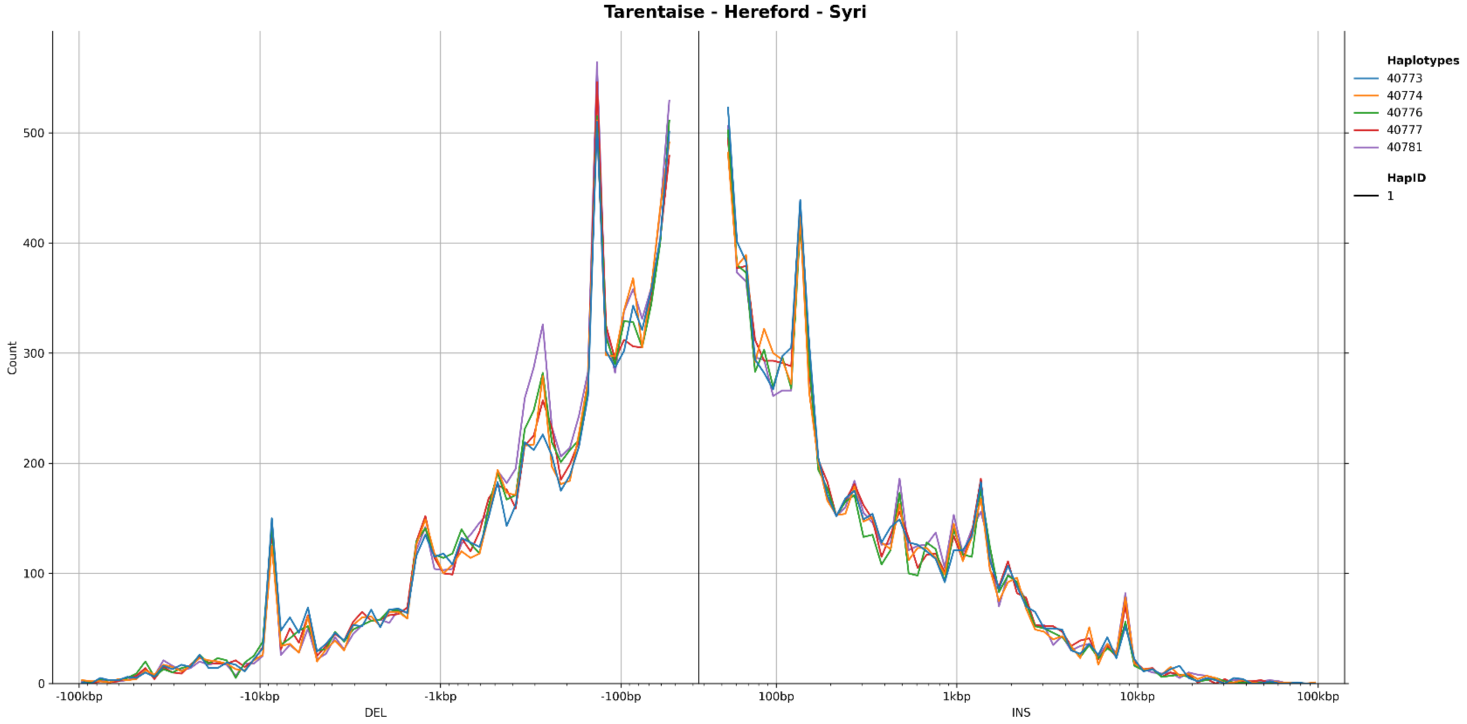


**Figure S27** Size distribution of SVs classified as deletions (left) and insertions (right), identified using **a** Minigraph, and **b** SyRI for the Tarentaise breed

**a**

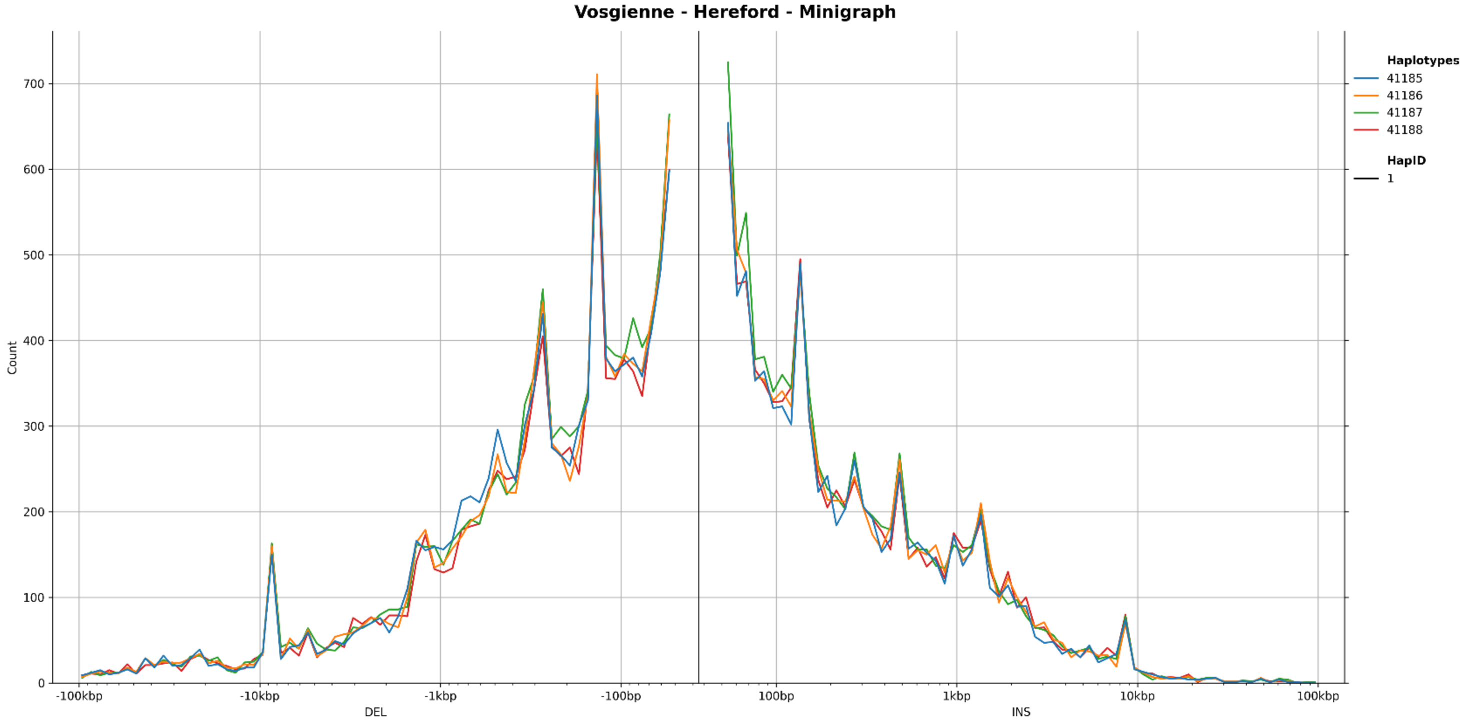


**b**

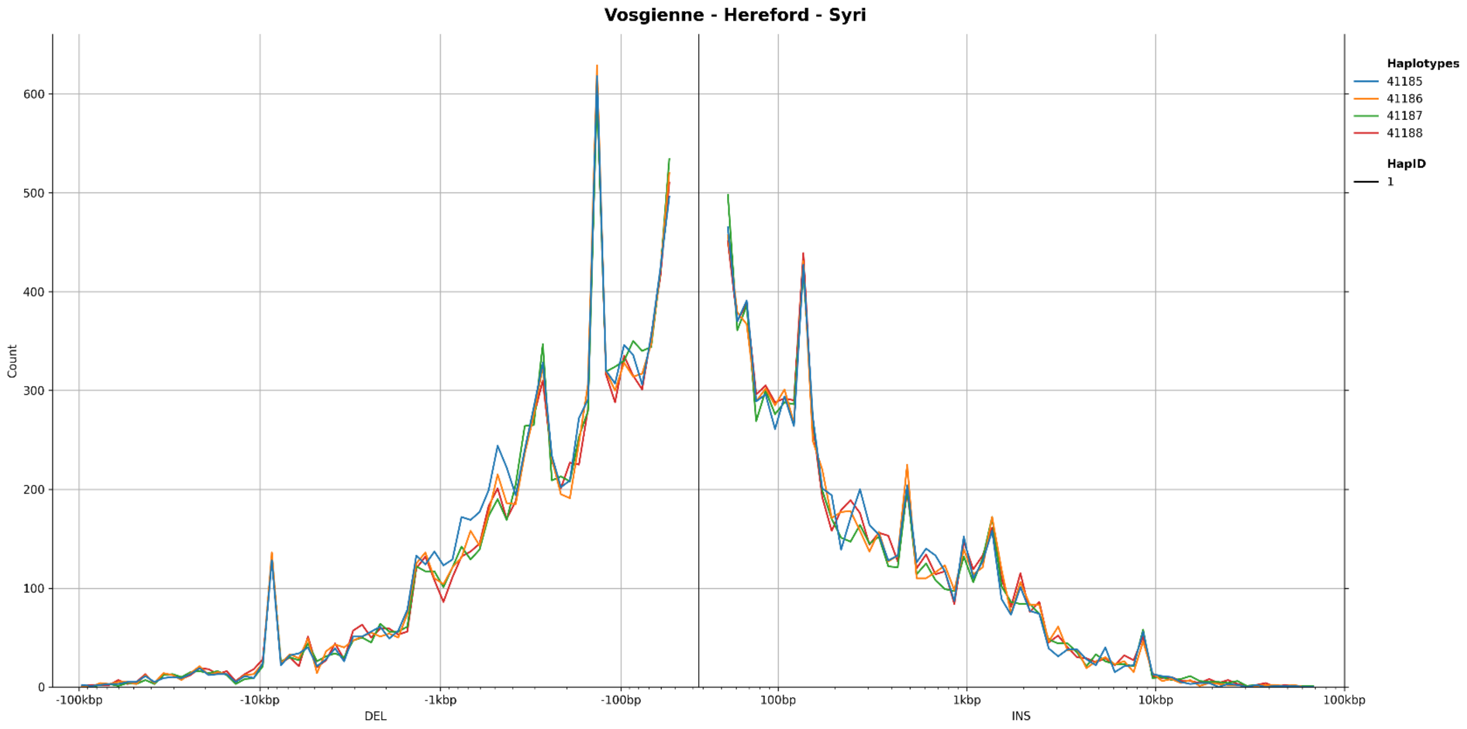


**Figure S28** Size distribution of SVs classified as deletions (left) and insertions (right), identified using **a** Minigraph, and **b** SyRI for the Vosgienne breed
