## Supplementary material for "Application of a French cattle pangenome, from structural variant discovery to association studies on key phenotypes": Access_to_supplemental_files: Additional file 5 Figures S29-S41.pdf

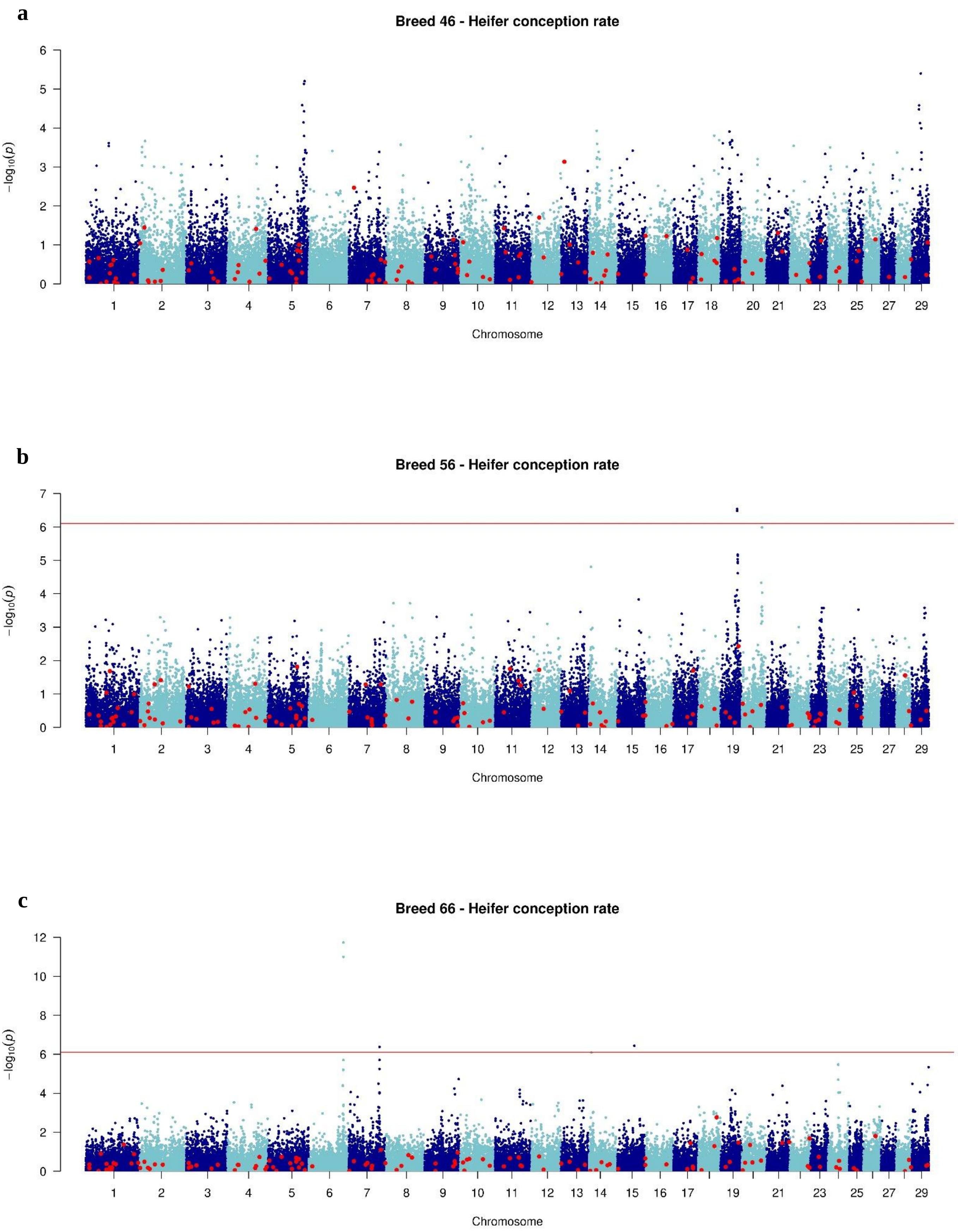


**Figure S29** Manhattan plot of GWAS analysis: -$\log_{10}$(P) values plotted against the positions of *Bos taurus* autosomes for variants associated with heifer conception rate in **a** Montbéliarde, **b** Normande, and **c** Holstein bulls

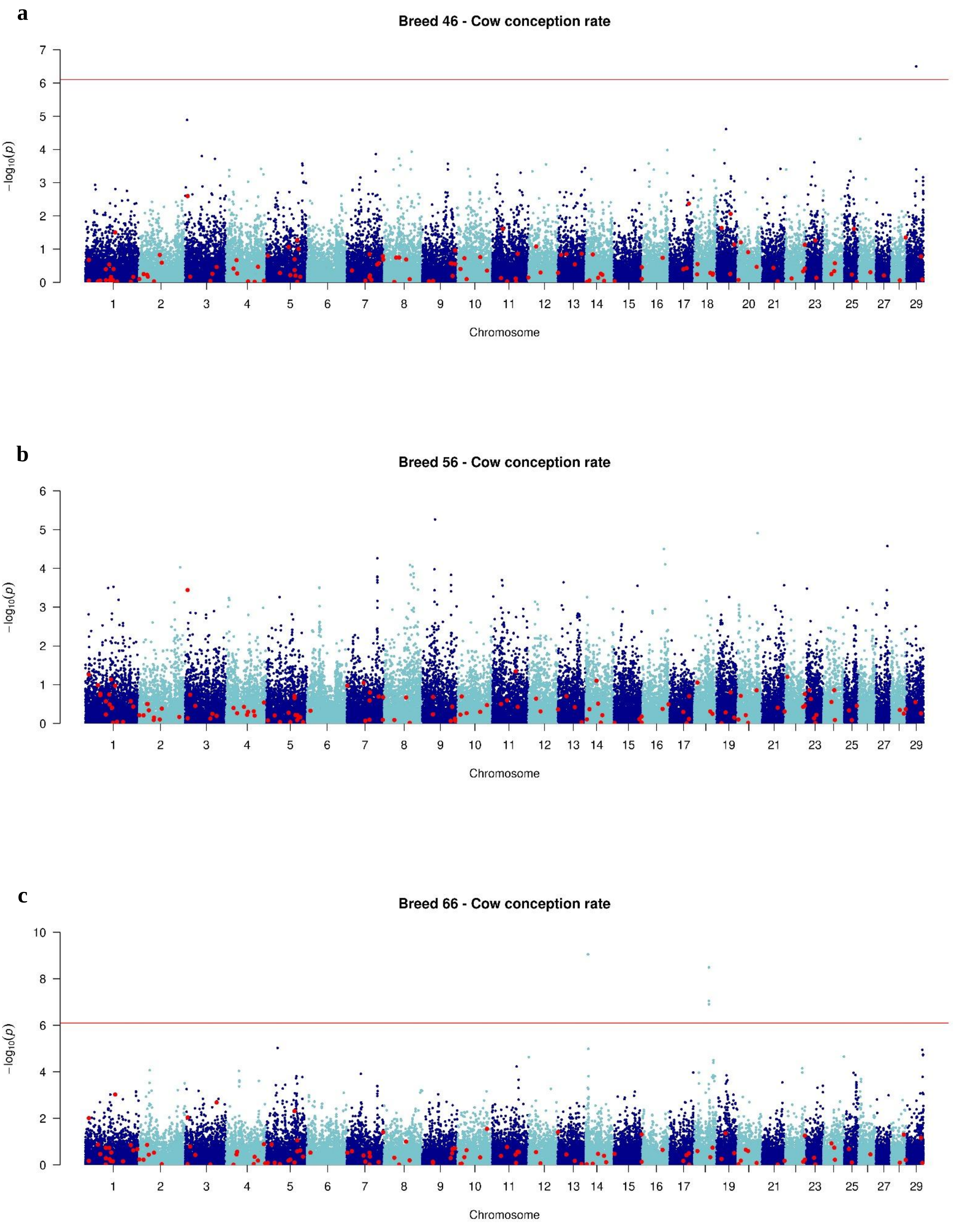


**Figure S30** Manhattan plot of GWAS analysis: -$\log_{10}$(P) values plotted against the positions of *Bos taurus* autosomes for variants associated with cow conception rate in **a** Montbéliarde, **b** Normande, and **c** Holstein bulls

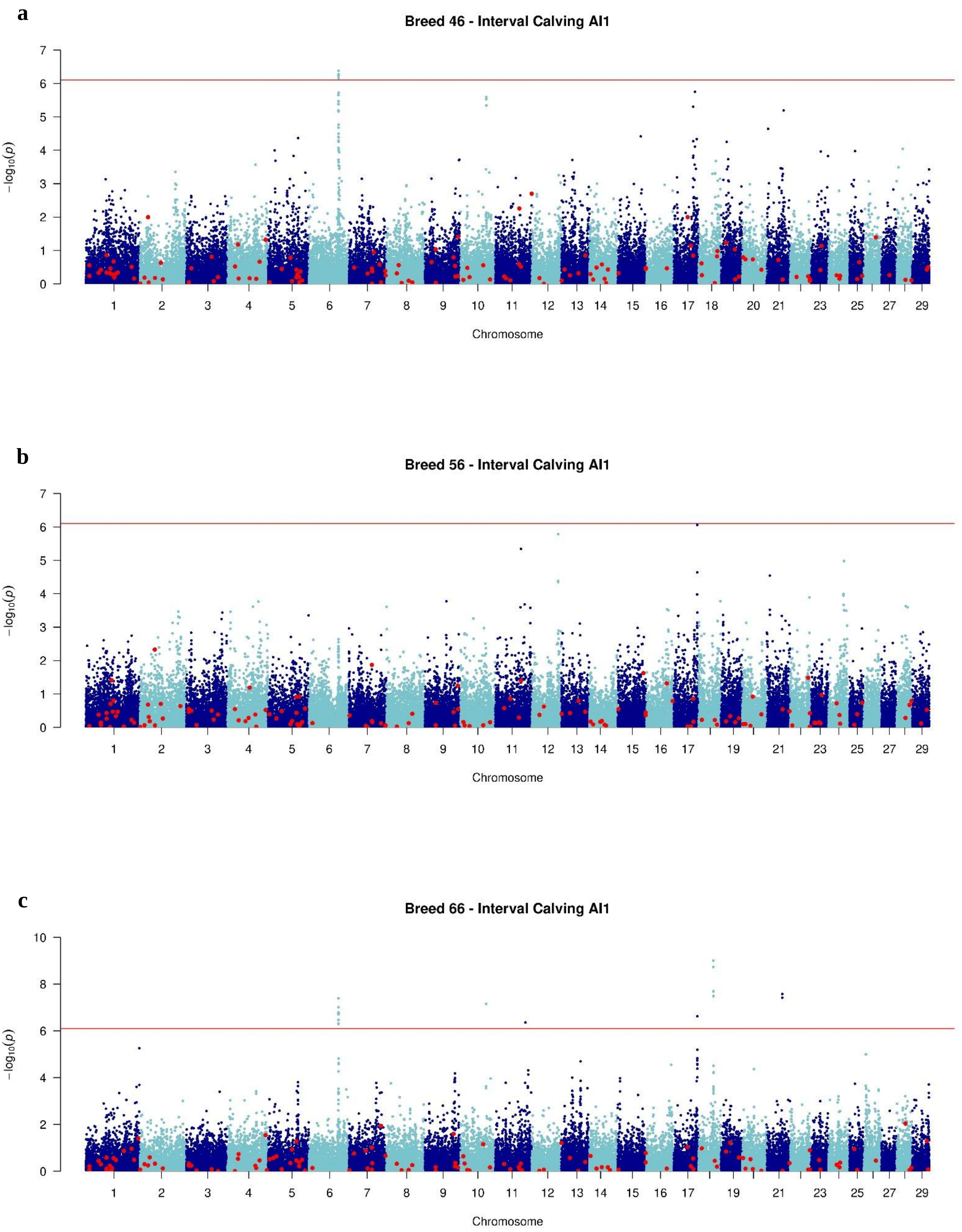


**Figure S31** Manhattan plot of GWAS analysis: -$\log_{10}$(P) values plotted against the positions of *Bos taurus* autosomes for variants associated with interval calving-AI1 in **a** Montbéliarde, **b** Normande, and **c** Holstein bulls

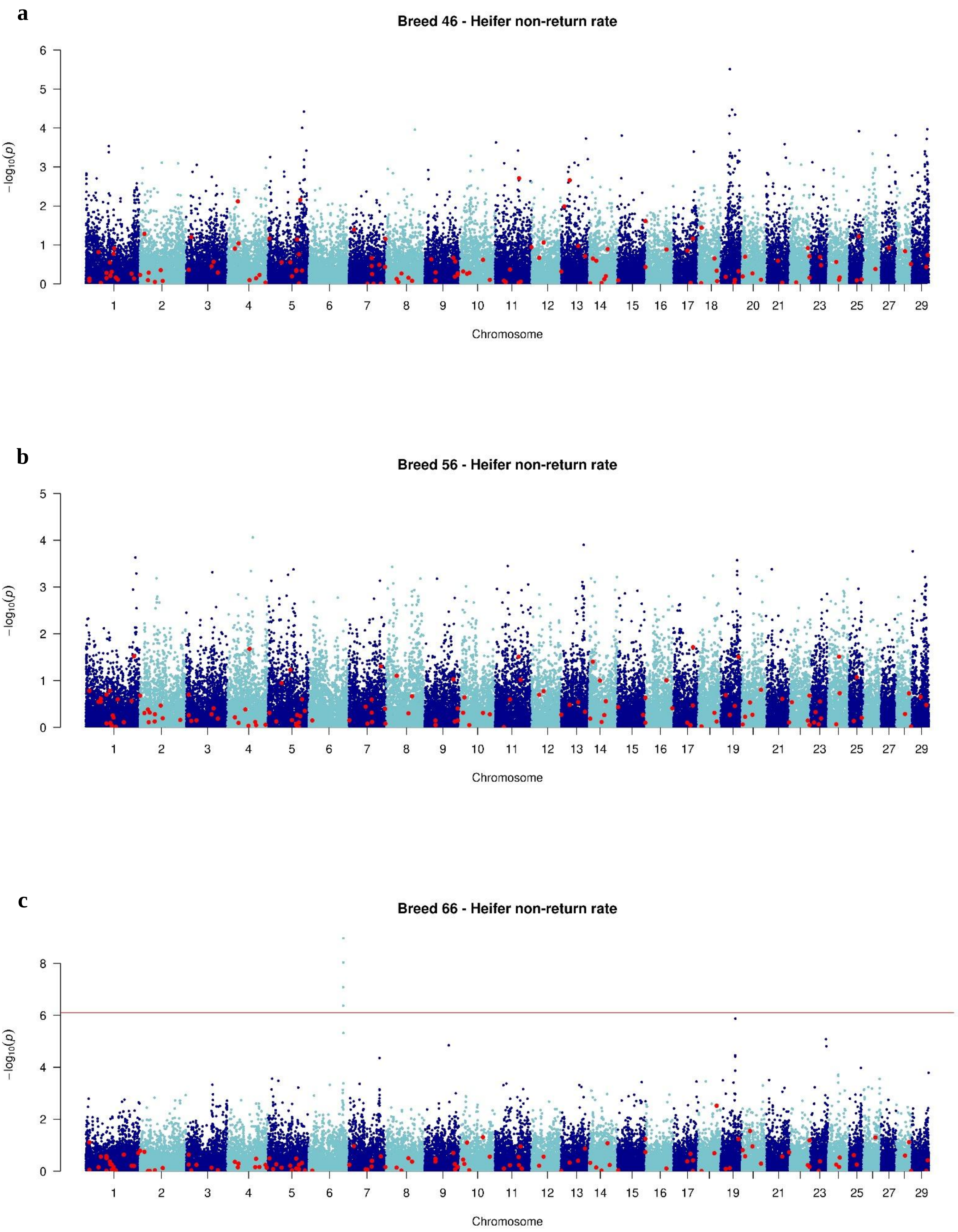


**Figure S32** Manhattan plot of GWAS analysis: -$\log_{10}$(P) values plotted against the positions of *Bos taurus* autosomes for variants associated with heifer non-return rate in **a** Montbéliarde, **b** Normande, and **c** Holstein bulls

**a**

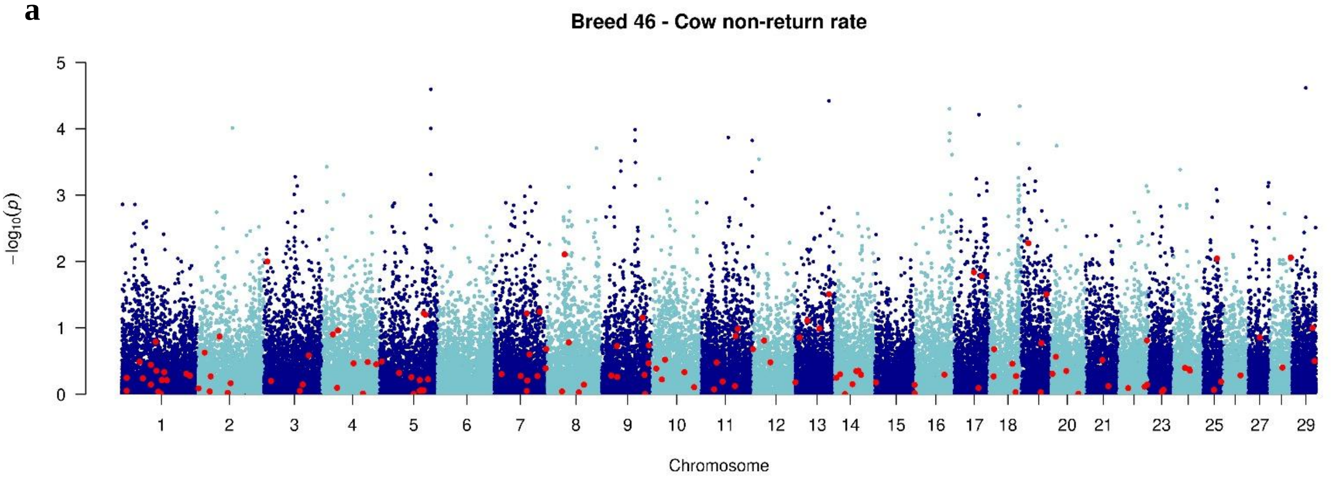


**b**

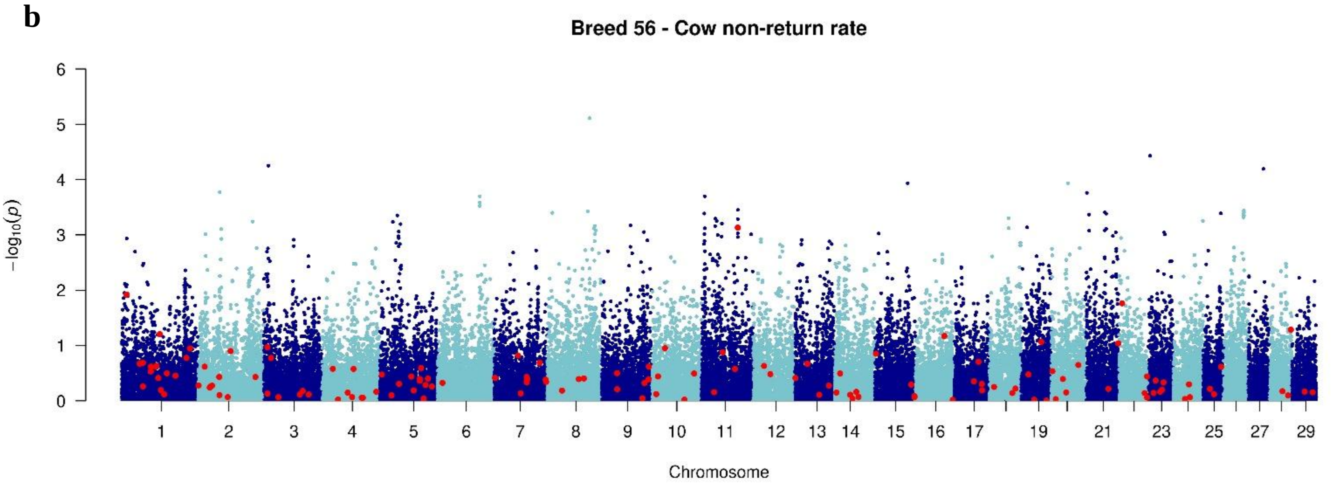


**c**

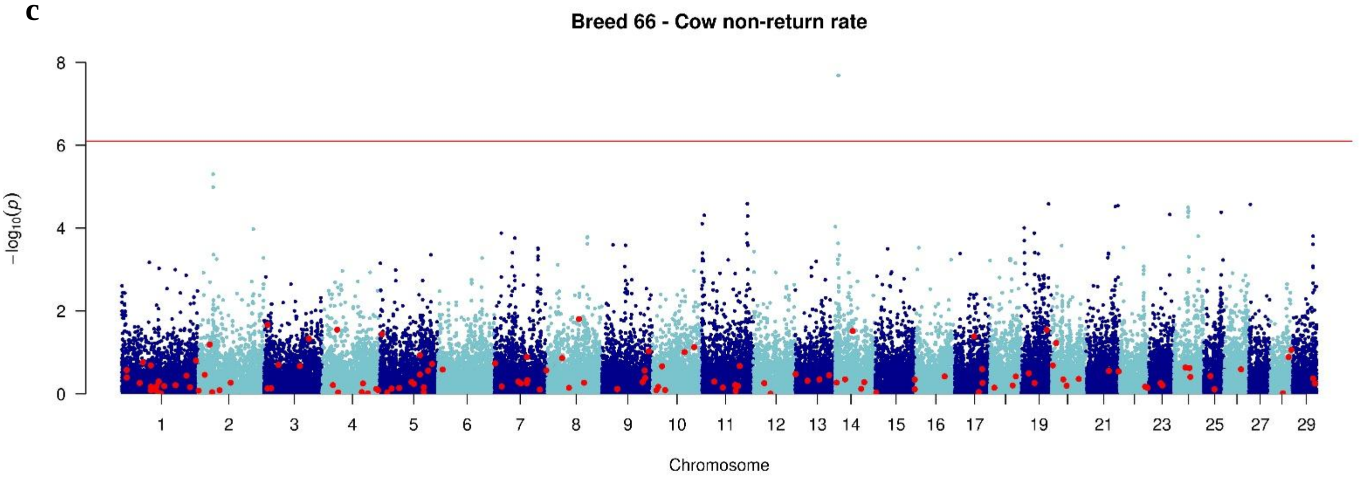


**Figure S33** Manhattan plot of GWAS analysis: -log$_{10}$(P) values plotted against the positions of *Bos taurus* autosomes for variants associated with cow non-return rate in **a** Montbéliarde, **b** Normande, and **c** Holstein bulls

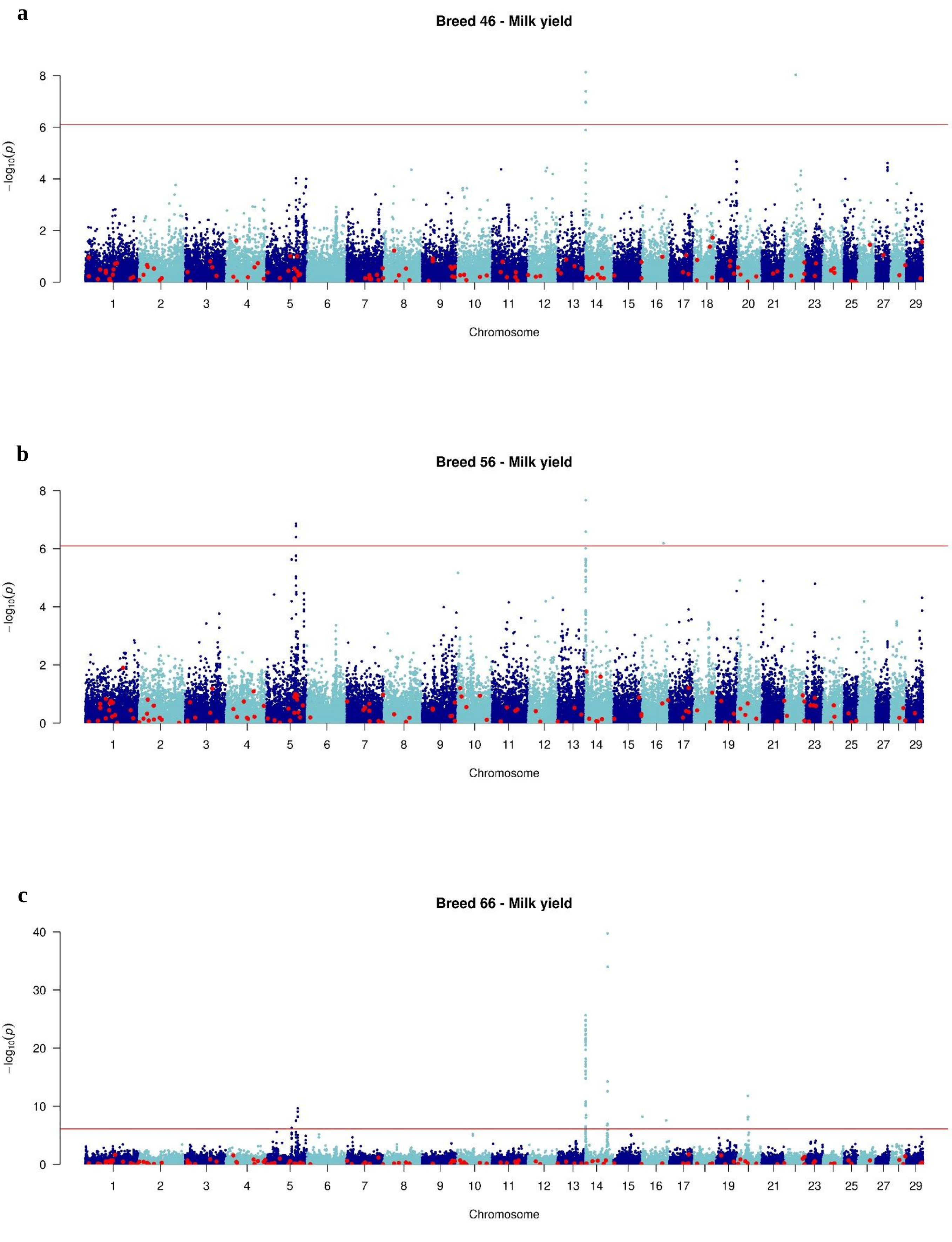


**Figure S34** Manhattan plot of GWAS analysis: -log$_{10}$(P) values plotted against the positions of *Bos taurus* autosomes for variants associated with milk yield in **a** Montbéliarde, **b** Normande, and **c** Holstein bulls

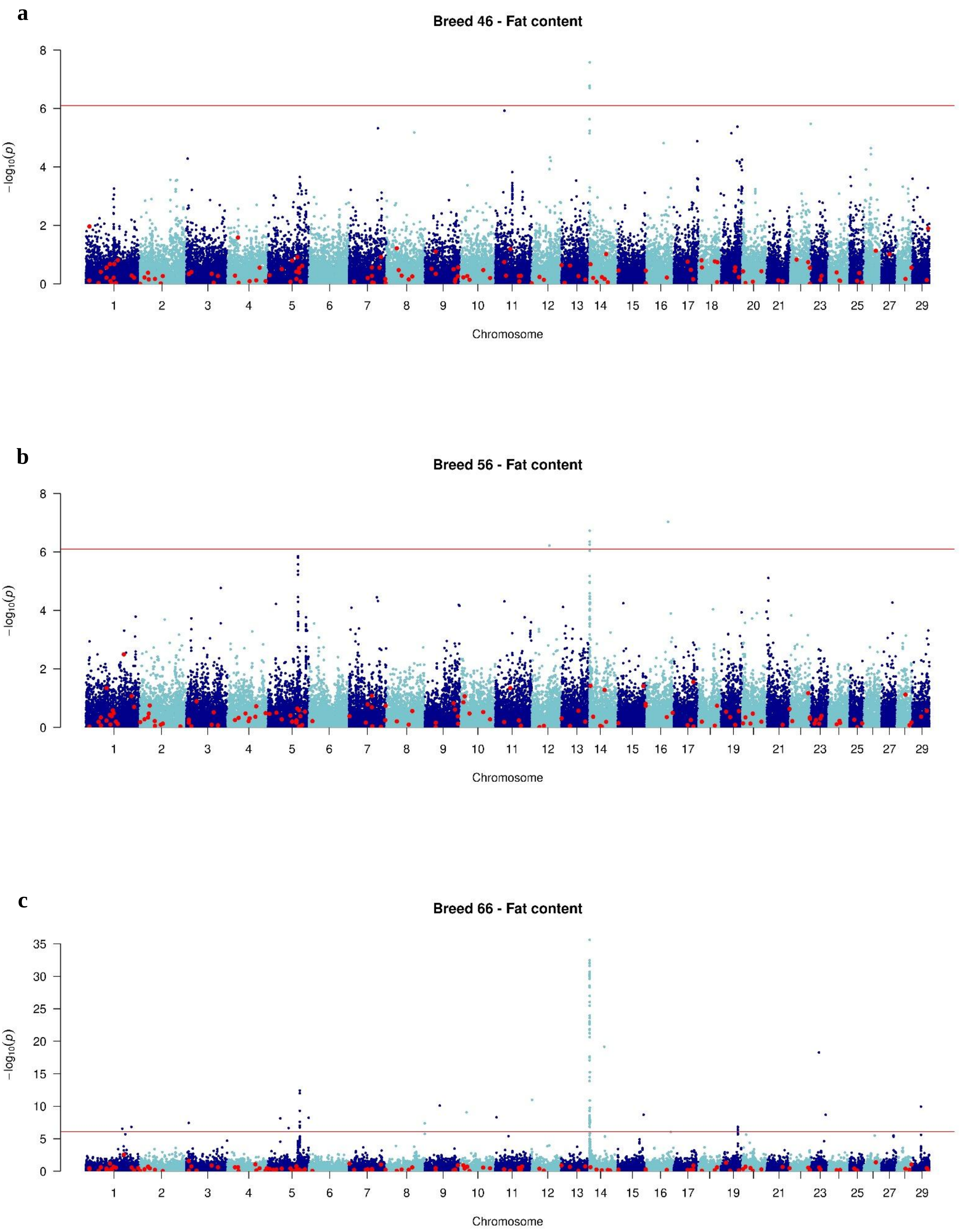


**Figure S35** Manhattan plot of GWAS analysis: -$\log_{10}$(P) values plotted against the positions of *Bos taurus* autosomes for variants associated with fat content in **a** Montbéliarde, **b** Normande, and **c** Holstein bulls

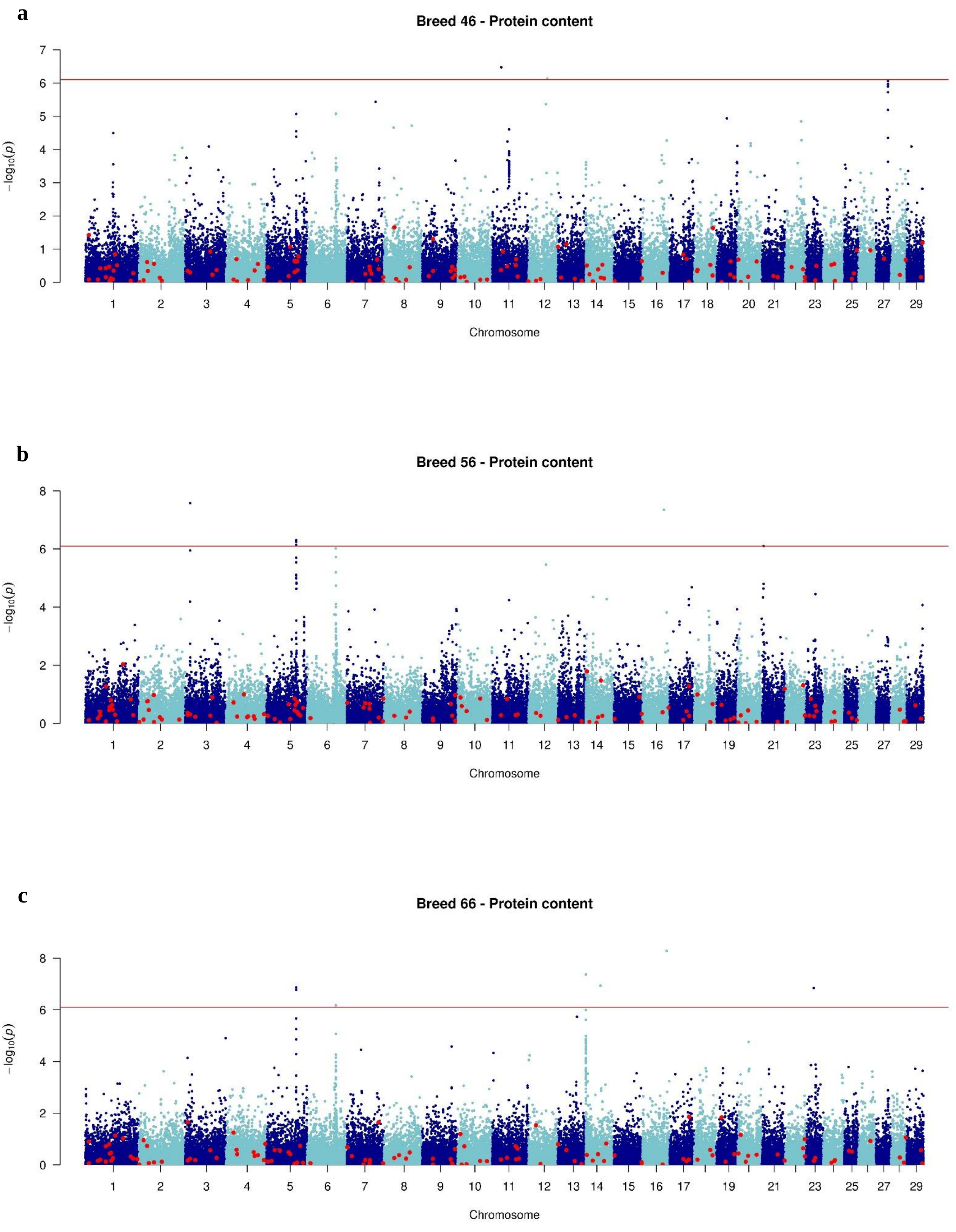


**Figure S36** Manhattan plot of GWAS analysis: -$\log_{10}$(P) values plotted against the positions of *Bos taurus* autosomes for variants associated with protein content in **a** Montbéliarde, **b** Normande, and **c** Holstein bulls

**a**

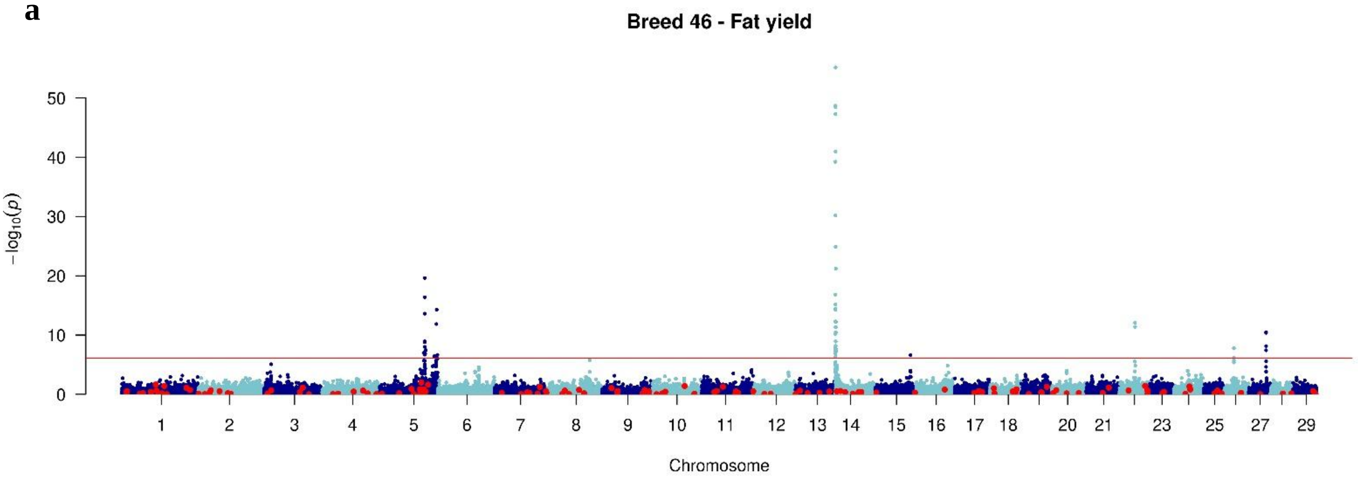


**b**

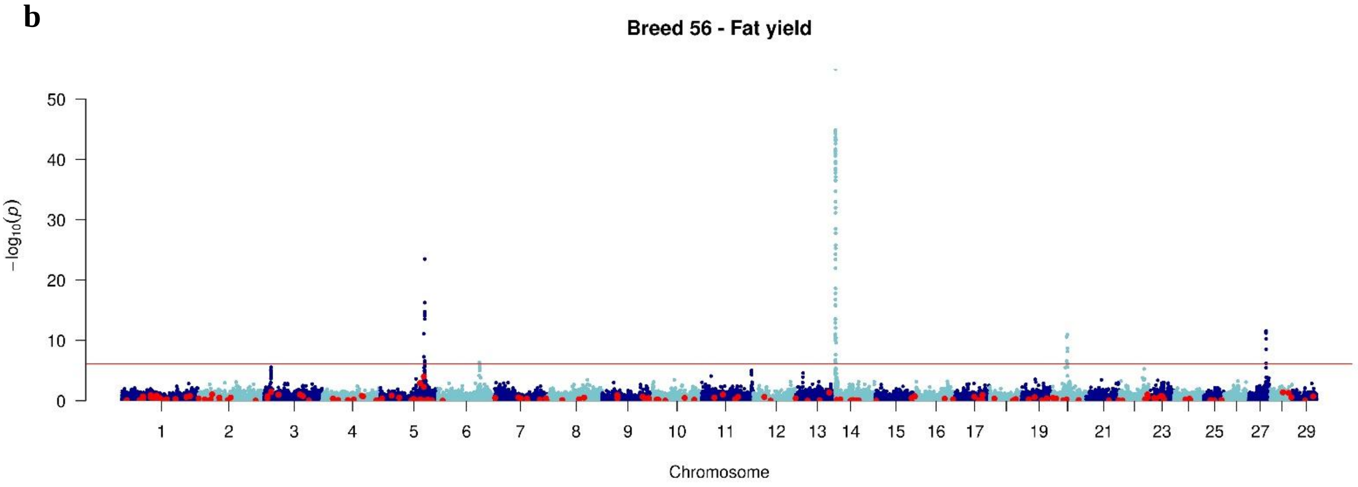


**c**

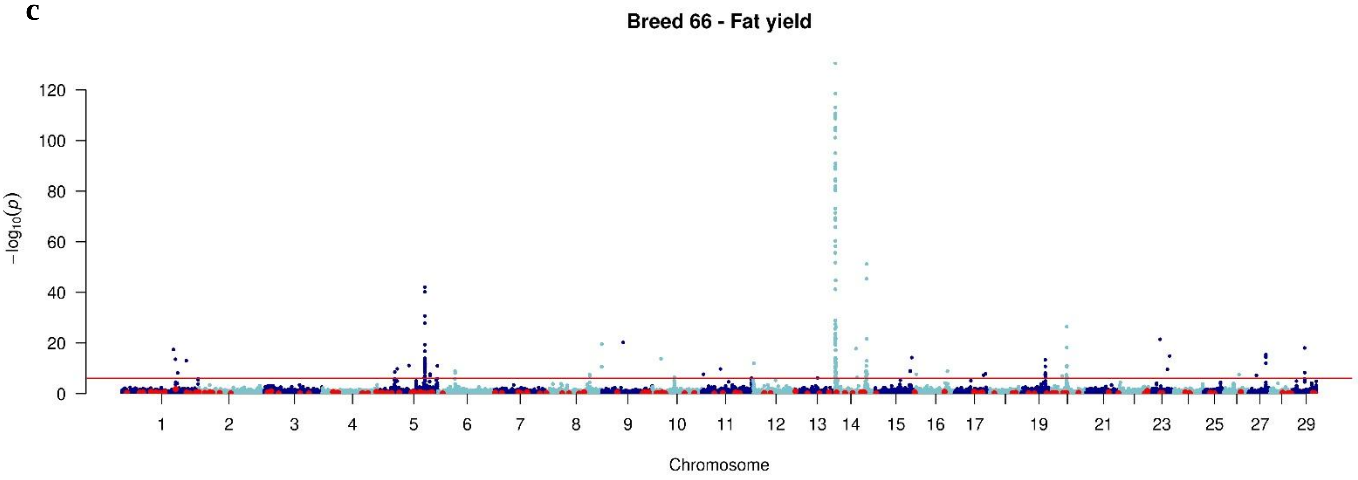


**Figure S37** Manhattan plot of GWAS analysis: -$\log_{10}$(P) values plotted against the positions of *Bos taurus* autosomes for variants associated with fat yield in **a** Montbéliarde, **b** Normande, and **c** Holstein bulls

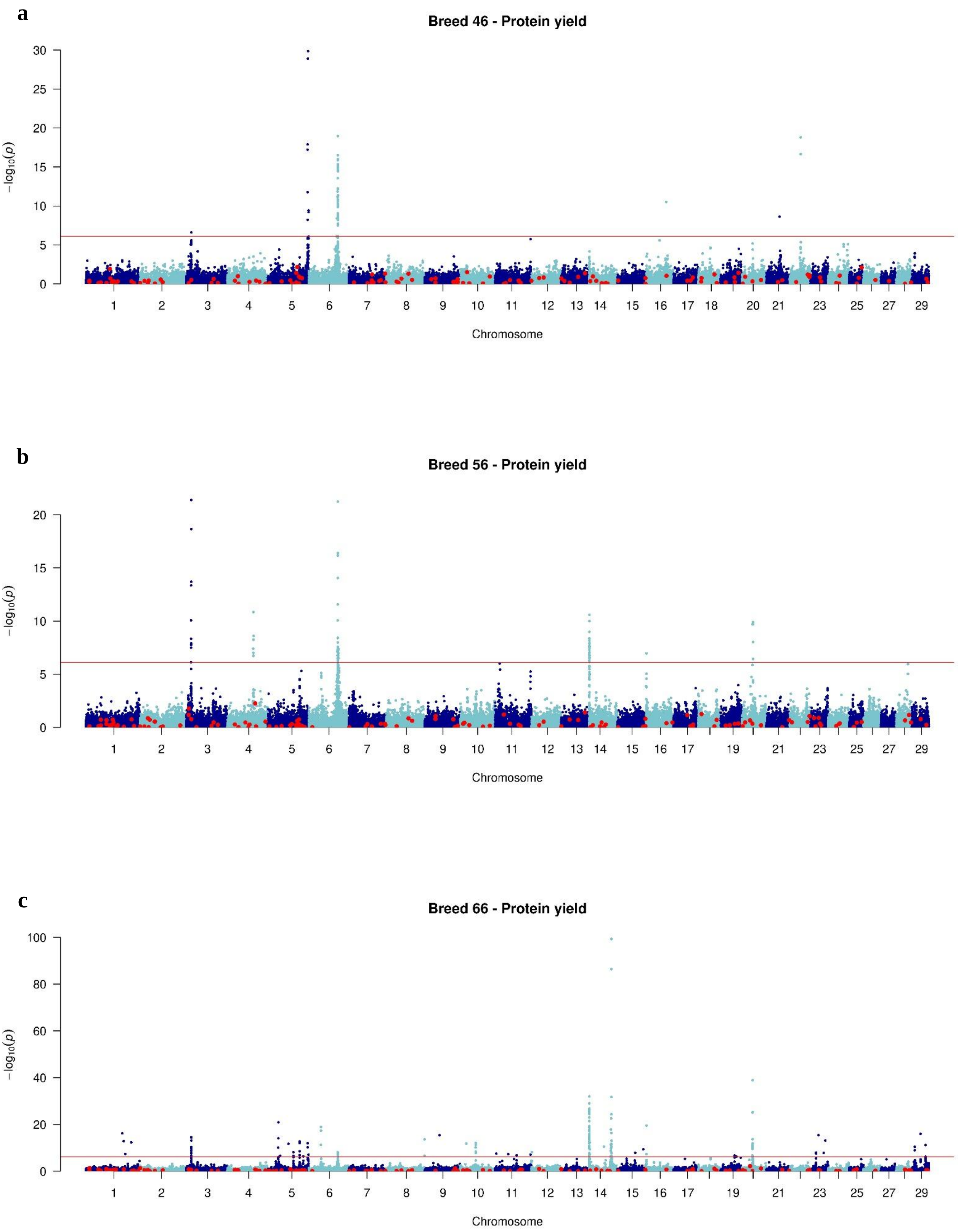


**Figure S38** Manhattan plot of GWAS analysis: -log$_{10}$(P) values plotted against the positions of *Bos taurus* autosomes for variants associated with protein yield in **a** Montbéliarde, **b** Normande, and **c** Holstein bulls

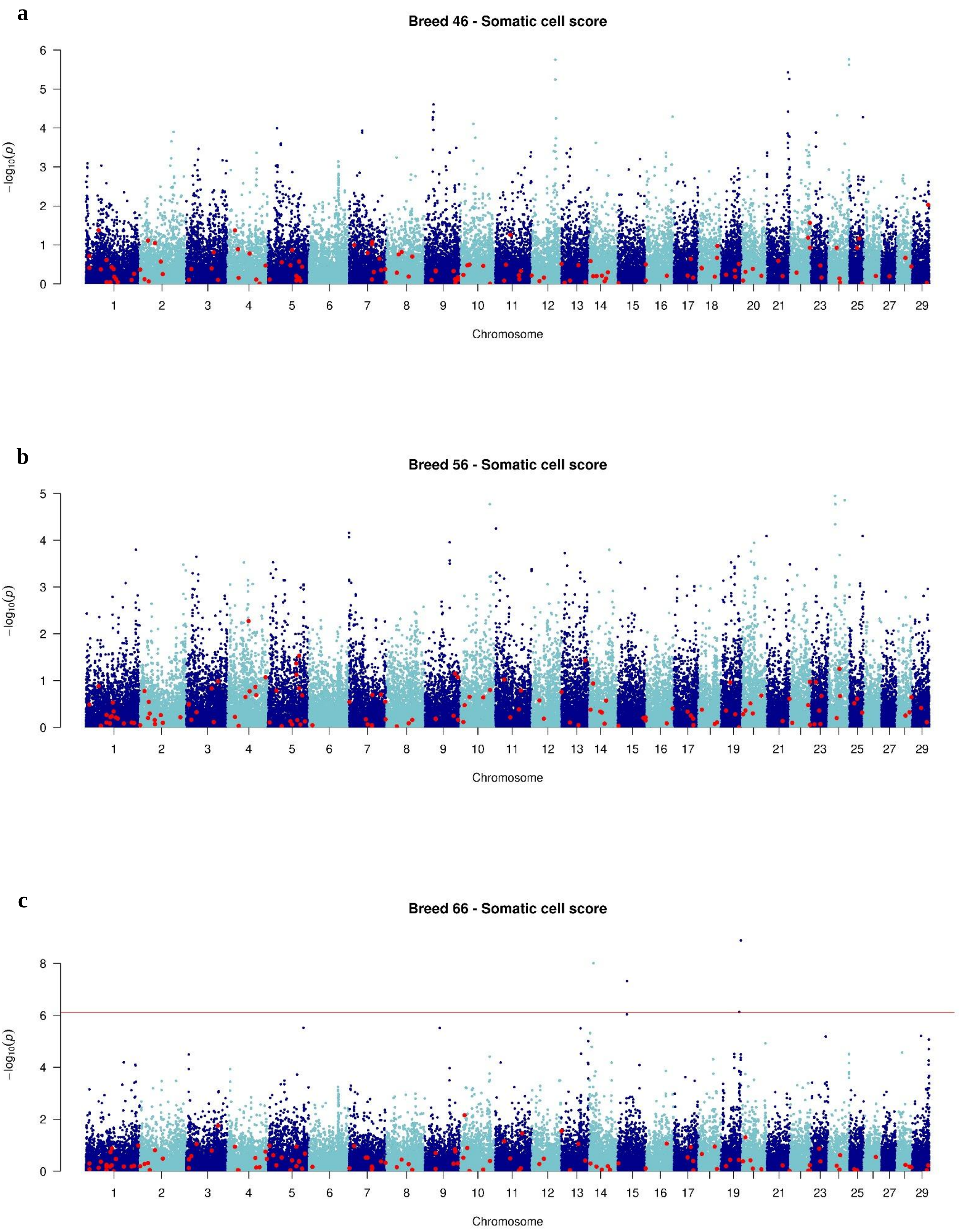


**Figure S39** Manhattan plot of GWAS analysis: -log$_{10}$(P) values plotted against the positions of *Bos taurus* autosomes for variants associated with somatic cell score in **a** Montbéliarde, **b** Normande, and **c** Holstein bulls

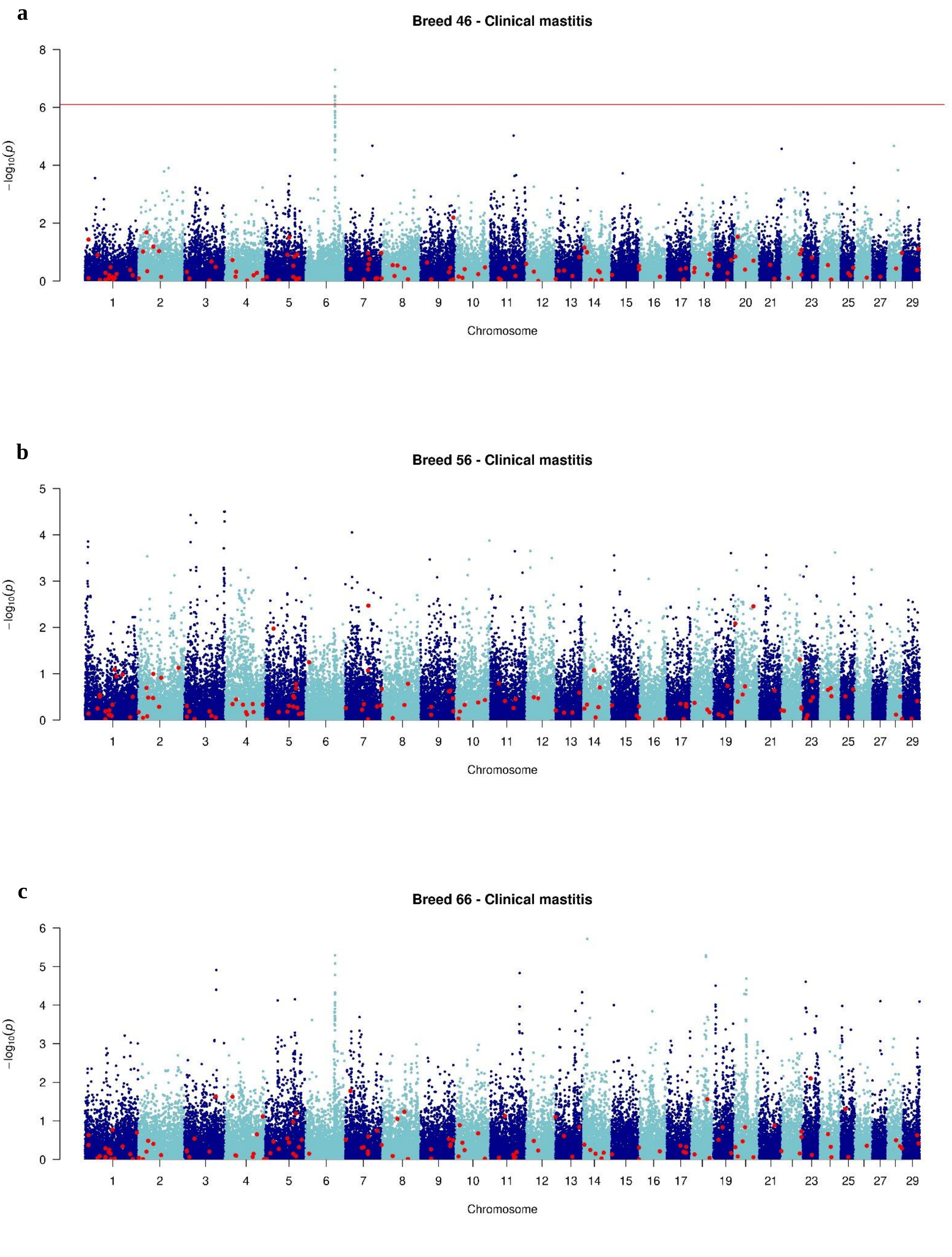


**Figure S40** Manhattan plot of GWAS analysis: -$\log_{10}$(P) values plotted against the positions of *Bos taurus* autosomes for variants associated with clinical mastitis in **a** Montbéliarde, **b** Normande, and **c** Holstein bulls

a

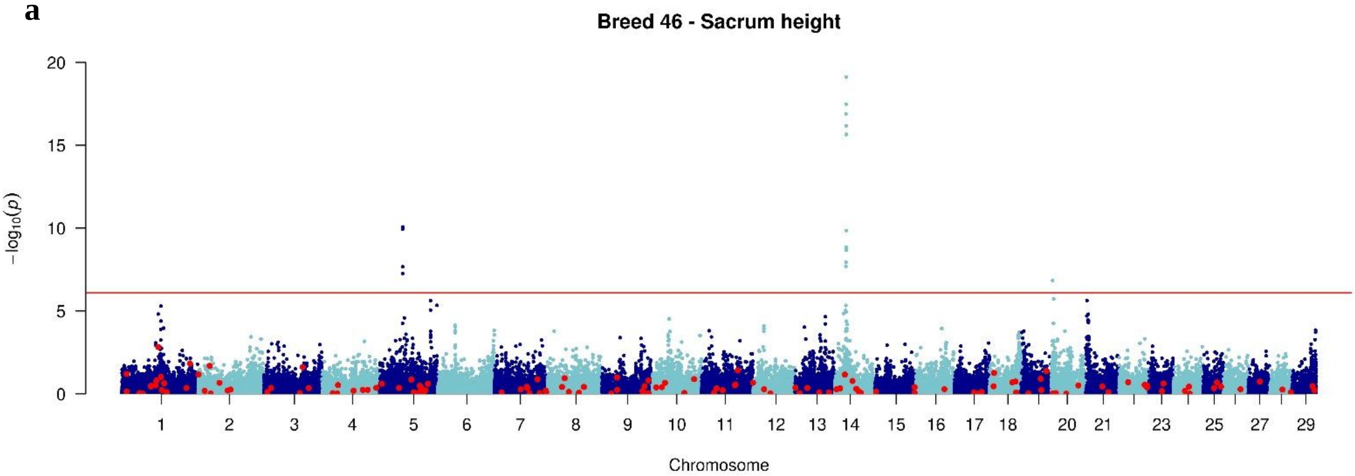


b

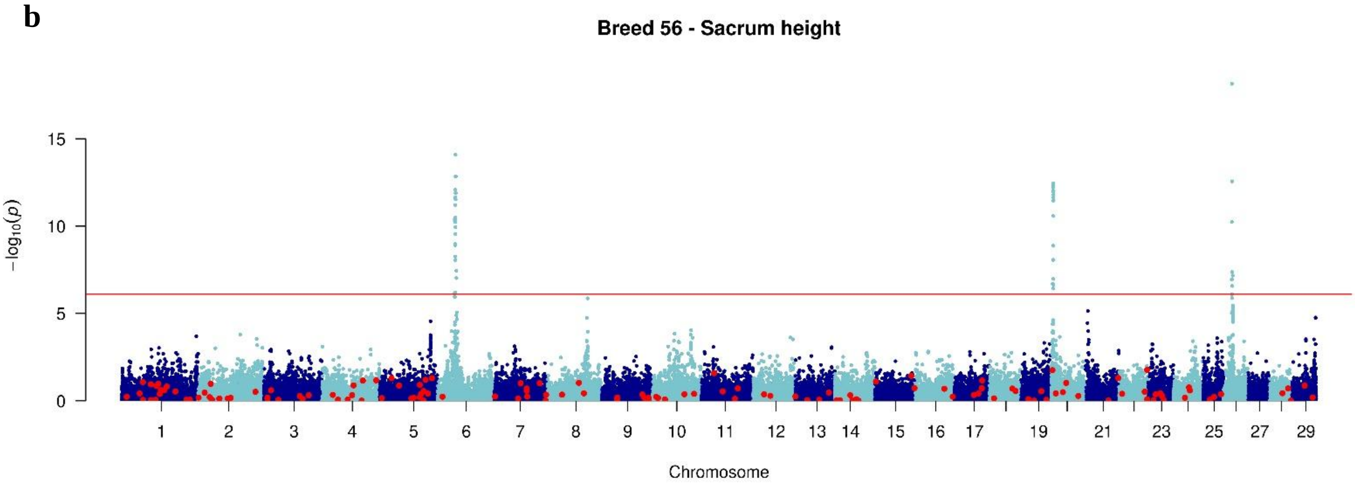


c

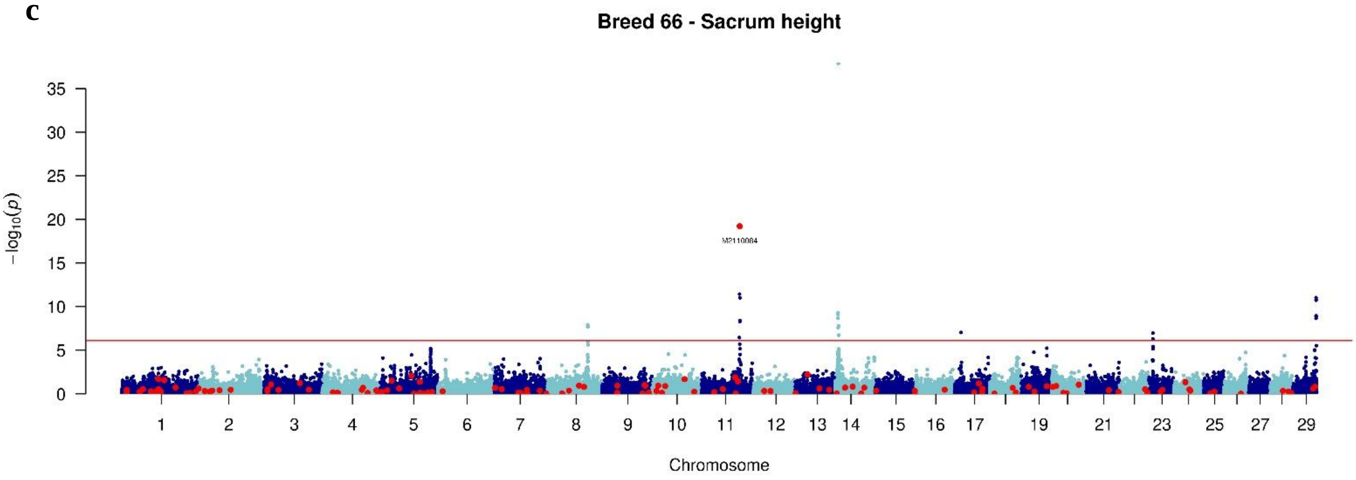


**Figure S41** Manhattan plot of GWAS analysis: -$\log_{10}$(P) values plotted against the positions of *Bos taurus* autosomes for variants associated with sacrum height in **a** Montbéliarde, **b** Normande, and **c** Holstein bulls
